# Learning from human and chemical languages to predict biological function

**DOI:** 10.64898/2026.08.09.743788

**Authors:** Clayton W. Kosonocky, Nikol Kadeřábková, Kangsan Kim, Ayesha J.S. Mahmood, Alexander Dunmyre, Phillip Woolley, Kristi Xing, Daniel Winkler, Tomer Babu, Filip Kadeřábek, Jonathan L. Sessler, Eric V. Anslyn, Edward M. Marcotte, Y. Jessie Zhang, Andrew D. Ellington, Despoina A.I. Mavridou

## Abstract

Understanding how molecular structure encodes biological function remains a grand challenge in drug discovery. Here, we present PubCheF-1, a deep learning model that predicts literature-derived biological function directly from chemical structure. PubCheF-1 was trained on a dataset linking molecules to labels derived from the scientific articles in which they appear, a strategy that connects disparate compounds through the language used to describe their functionalities. When tasked with identifying inhibitors of β-lactamases, including enzymes considered largely refractory to inhibition, PubCheF-1 predicted structurally distinct compounds that collectively have activity against all β-lactamase classes. Furthermore, hit compounds directly bind the enzyme active site, restore antibiotic efficacy in multidrug-resistant high-priority pathogens, and demonstrate potent activity in animal infection models. Together, these findings establish that machine learning-based prediction of biological function derived from the language of scientific literature allows the identification of bioactive molecules at high hit rates, thereby accelerating therapeutic discovery.

## INTRODUCTION

Deep learning methods have recently been successfully applied to small molecule drug discovery [1–3]. Many of these approaches learn directly from protein-ligand structures, reflecting the central role of molecular recognition in drug action [4,5]. However, high-quality co-crystal structures cover only a narrow and biased fraction of relevant chemistry within the vastness of chemical space, and obtaining sufficient additional structures is costly [6,7]. Moreover, structure-based methods primarily model binding affinity, which is often necessary for biological activity but does not fully capture how compounds behave within living systems, from their off-target effects to disease-level activities and potential toxicity. These limitations suggest that complementary approaches are needed for the robust prediction of bioactive small molecules [6].

Large repositories such as PubChem and ChEMBL have catalogued millions of molecules together with the outcomes of the experiments performed on them [8,9]. Because these relationships span a wide range of biological functions, they could, in principle, be used to train a model that predicts how a chemical structure generally affects a biological system. However, unifying disparate measurements is not straightforward. Readouts range from receptor binding affinity to the inhibition of cancer cell growth, and even data for the same endpoint, such as IC50 values, can vary substantially across sources [10]. One potential solution is to bridge experimental outcomes and analyses through the semantics of natural language. Specifically, we hypothesized that the relational structures inherent in both human language and chemical space might be coherently bridged to enable the prediction of a compound’s biological function. Historically, such functional information has been difficult to parse at scale because it is dispersed across unstructured text, with databases such as ChEMBL requiring extensive manual curation [10]. Therefore, we sought to systematically extract high-quality functional annotations from the scientific literature using the summarization capabilities of large language models (LLMs) and to generate a dataset of molecules and labels representing their associated biological functions [11,12]. Using this dataset, we trained PubCheF-1, a deep learning model that can predict the functional profile of any arbitrary molecular structure. We further posited that the functional profiles predicted by PubCheF-1 could be used to discover bioactive molecules, particularly in applications limited by sparse knowledge, low hit rates, and the high cost of experimental scaling.

As a stringent test of this approach (**Figure 1**), we focused on antimicrobial resistance, which constitutes one of the most urgent threats to global health [13,14]. β-Lactam compounds account for approximately two-thirds of clinically prescribed antibiotics [15,16] because of their broad activity and favorable safety profiles [16]. Yet their widespread use has also driven the spread of resistance, contributing substantially to resistance-associated mortality [17]. A major driver of antibiotic resistance is the emergence of β-lactamases, enzymes that hydrolyze the β-lactam ring of the drugs [18] and disseminate rapidly through horizontal gene transfer [19–22]. More than 7,700 β-lactamases belonging to hundreds of phylogenetic families have been identified to date [23], creating extensive sequence and functional diversity that facilitates the rapid emergence of resistance to any newly introduced therapy [21]. Traditionally, β-lactam-resistance is countered by using β-lactamase inhibitors that potentiate the antibiotics [15,24–27]. Although these compounds have transformed clinical treatment strategies over the past 50 years [15], continued evolution and expansion of β-lactamase families increasingly outpace conventional inhibitor discovery efforts [21]. Compounding this problem is the lack of large, high-quality datasets of resistance-breaking compounds and β-lactamase inhibitors [28,29], a sparse knowledge challenge that restricts the application of deep learning approaches to inhibitor discovery [30,31]. Collectively, inhibition of β-lactamases serves as a demanding test of whether a literature-centric deep learning model such as PubCheF-1 can reveal bioactive molecules, particularly against clinically important enzymes for which inhibitor options remain limited, like class B metallo-β-lactamases and class D oxacillinases [24] (**Figure 1**).

**Figure 1.**
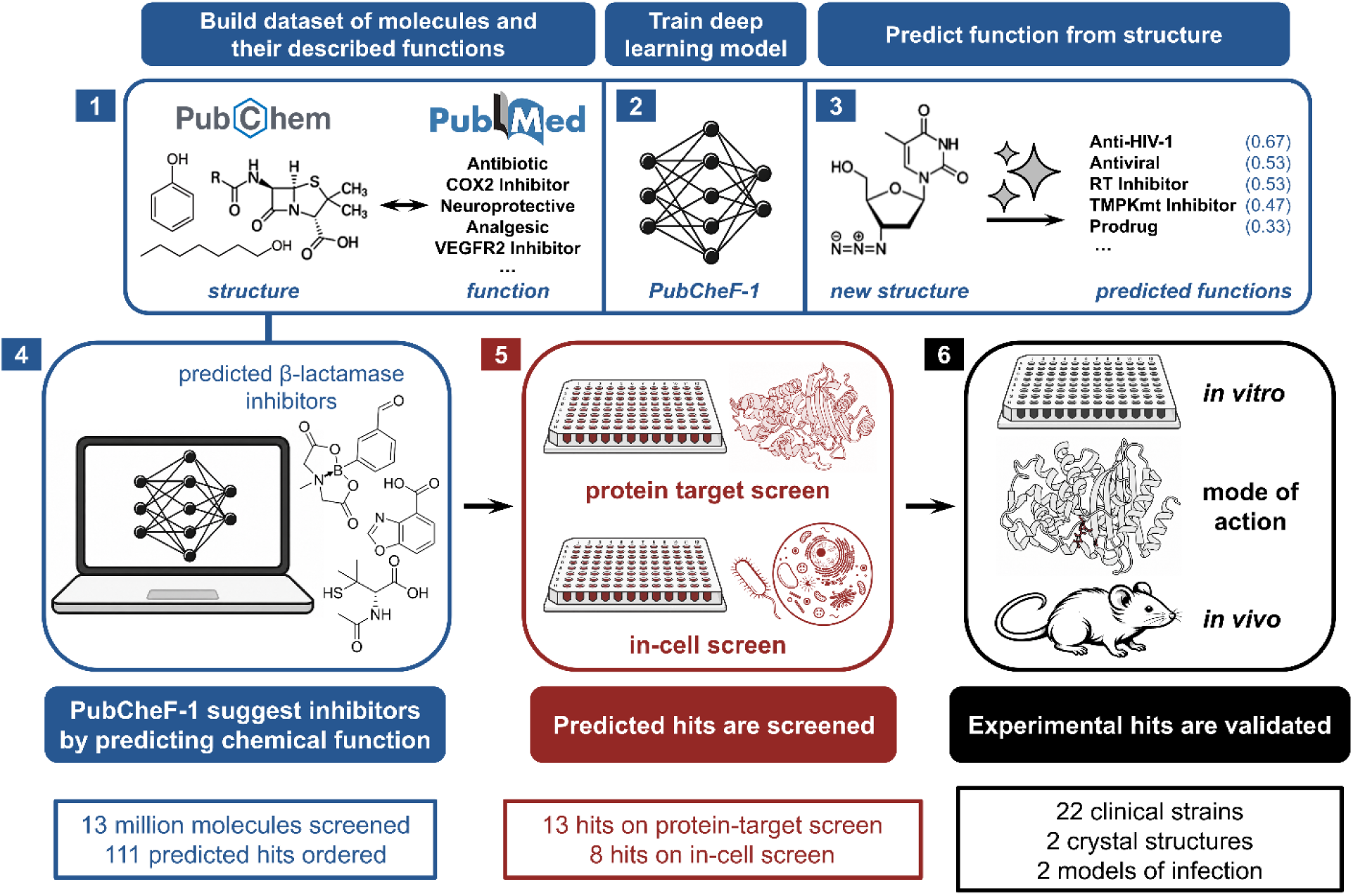
Schematic of the PubCheF framework for the identification and validation of bioactive compounds. Data mining of PubChem and PubMed yields a dataset of chemical structures and their functional annotations **(1)**. This results in the PubMed-derived Chemical Function dataset (PubCheF), which is used to train the PubCheF-1 deep neural network model **(2)**. From a given chemical structure, PubCheF-1 predicts a ranked list of biological functions **(3)**, enabling prioritization of molecules associated with functions of interest **(4)**. Predicted *in silico* hits are triaged for toxicity and novelty before they are subjected to experimental validation to identify compounds with on-target activity and cellular efficacy **(5)**. Experimental hits are then validated using biochemical characterization and mode of action studies, and the best hits are subjected to *in vivo* testing to assess efficacy, biological relevance, and safety **(6)**. Arrows indicate the flow from computational prediction to iterative experimental testing and validation.

## RESULTS

### Building the PubCheF dataset

We sought to create a high-quality dataset of chemical structures paired with labels corresponding to their literature-derived biological functions. In previous work, we used large language models to extract functional descriptors for chemicals from patents to create the Chemical Function (CheF) dataset [11]. Here, we focused on primary scientific literature and implemented new methods to improve dataset quality and scale for downstream modeling and experimental validation (**Figure 2A**).

**Figure 2.**
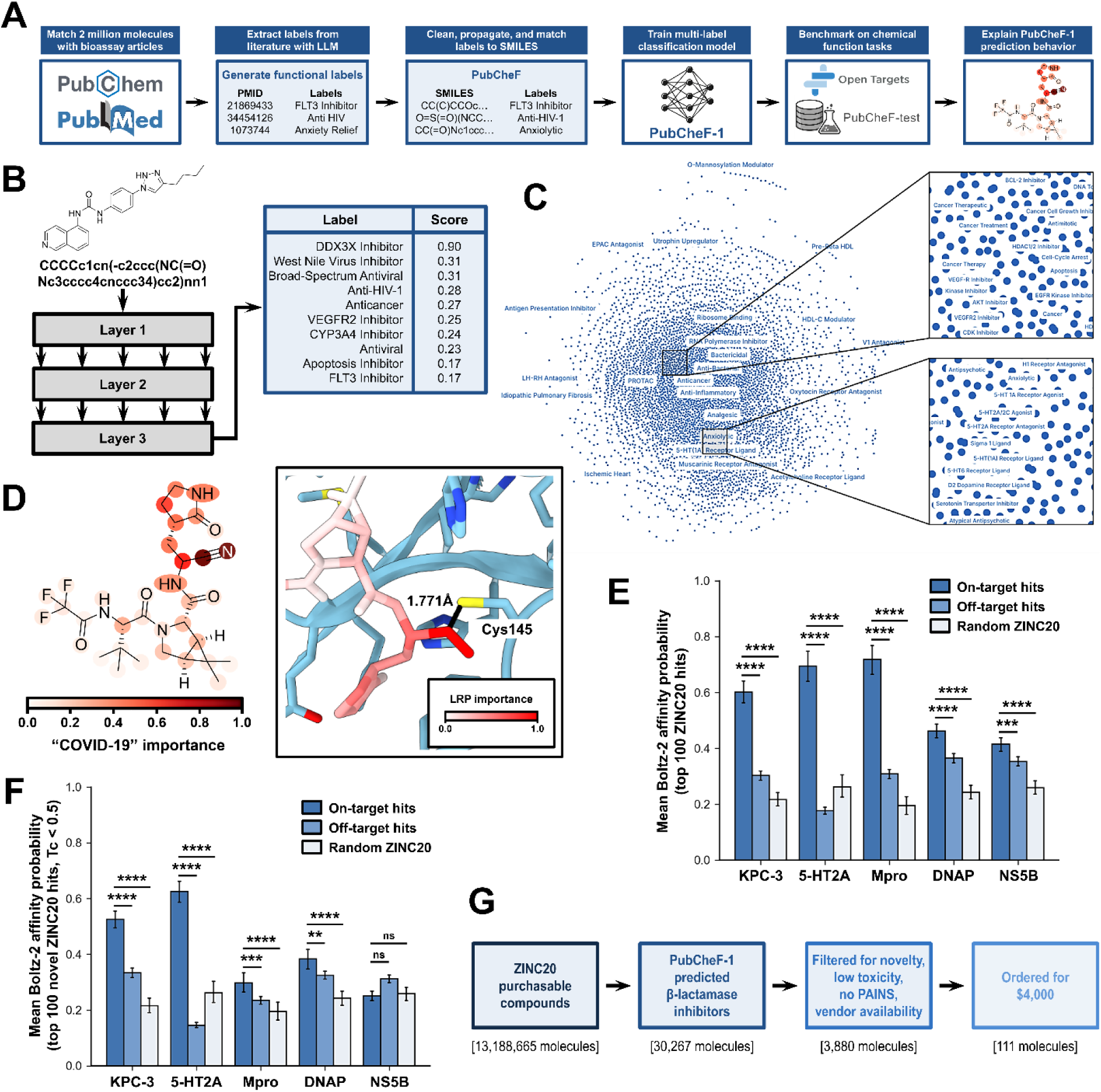
PubCheF-1 predicts inhibitors by modeling the chemical function landscape. (A) Workflow used to create and validate the PubCheF-1 deep neural network model. Titles and abstracts from PubMed articles were fed to a large language model to generate functional labels. These labels were then cleaned, matched to SMILES strings, and propagated to identical structures, resulting in the PubCheF dataset. PubCheF was used to train a multi-label classification model, PubCheF-1, which was benchmarked on both the OpenTargets-indications and PubCheF-test-reduced datasets. A series of explainability analyses were then performed to understand the prediction behavior of PubCheF-1. **(B) PubCheF-1 inference diagram.** Molecules are represented as SMILES strings and input into a RoBERTa model architecture. The classification token embedding is projected to a linear layer with dimensionality equal to the number of classes in the PubCheF dataset. The logits are converted to multi-label classification scores, where each label is assigned a separate normalized score of being applicable to the input molecule. **(C) PubCheF labels organized by molecular co-occurrence.** Labels that occur together frequently are closer together in space, leading to a visually coherent semantic structure. Examples include one cluster of terms related to cancer therapy and another related to neuromodulators. **(D) Model importance for label “COVID-19” is biochemically meaningful**. (**left**) LRP importance for the SARS-CoV-2 main protease (Mpro) inhibitor nirmatrelvir when predicting the “COVID-19” label. Most of the importance is attributed to a warhead moiety that covalently binds to Mpro through Cys145 and inactivates it (PDB:8DZ2; **right**). **(E) Boltz-2 affinity probabilities for PubCheF-1 top predictions to five targets.** For five targets (KPC-3 β-lactamase, 5-HT2A receptor, SARS-CoV-2 Mpro, Taq DNA polymerase, and Hepatitis C Virus NS5B RNA polymerase), Boltz-2 affinity probabilities for the top 100 PubCheF-1 predicted ZINC20 in-stock molecules to each target (on-target hits) were compared to the affinities of the other targets’ ligand sets (off-target hits) and a 100 random ZINC20 molecule control. The on-target hits for all five targets had enriched Boltz-2 affinity probabilities compared to the off-target hits and random ZINC20 molecule control. **(F) Boltz-2 affinity probabilities for PubCheF-1 top novel predictions to five targets.** For the same five targets, Boltz-2 affinity probabilities for the top 100 novel (Tc < 0.50 to any molecule in PubCheF dataset containing the selected-for labels) PubCheF-1 predicted ZINC20 in-stock molecules to each target (on-target hits) were compared to the affinities of the other targets’ ligand sets (off-target hits) and a 100 random ZINC20 molecule control. The on-target hits for four of the five targets had enriched Boltz-2 affinity probabilities compared to the off-target hits and random ZINC20 molecule control. For (E) and (F) statistical analysis was performed using pairwise comparisons between groups with Mann-Whitney U tests with Benjamini-Hochberg correction for multiple comparisons; p < 0.05 (significance, *), p < 0.01 (significance, **), p < 0.001 (significance, ***), p < 0.0001 (significance, ****), p ≥ 0.05 (non-significance, ns). **(G) Using PubCheF-1 to predict β-lactamase inhibitors.** PubCheF-1 was run on the ZINC20 purchasable set of 13.2 million molecules to obtain ∼30,000 strongly predicted β-lactamase inhibitors. These were filtered to 3,880 molecules based on novelty, low predicted toxicity, absence of PAINS substructures, and vendor availability. A total of 111 molecules were then chosen from this set, optimizing for cost, diversity, and PubCheF-1 score.

First, we fine-tuned an LLM on the task of function extraction to improve its precision, recall, and generated vocabulary (**Supplementary Table S1**, **Supplementary File S1**). We then used this fine-tuned LLM to generate functional labels for all PubMed BioAssay-linked articles associated with PubChem [8] using their titles and abstracts (88,000 articles corresponding to 2.2 million molecules). Although many of these labels could have been derived from ChEMBL’s assay and target annotations, working directly from free text removes dependence on structured assay records, allowing the approach to extend, in principle, to sources that lack curated quantitative measurements [11]. We further improved the quality of the generated labels by clustering them based on their language model embeddings and mapping each label to the centroid label of its cluster (see **Materials and methods**). This strategy, applied to the full dataset, correctly preserved each cluster’s functional class and mode of action 100% of the time and, for terms indicating receptors, reflected the correct receptor subtype 92% of the time (n=100 clusters containing two or more terms; **Supplementary File S1**). After additional filtering and refinement (see **Materials and methods**), the final dataset, dubbed PubCheF (PubMed-derived Chemical Function dataset), contained 1,201,302 molecules and 5,125 distinct labels (**Supplementary Figure S2**).

A random subset of PubCheF was then manually validated to assess the accuracy of its label assignments (n=100 molecules, 186 molecule-label pairs; see **Materials and methods**). This showed that the molecules were indeed studied for their assigned functions in 98.9% of the pairs and were reported to be active in 89.7% of them (**Supplementary File S1**). In 99.5% of the pairs, either the molecule or a close derivative of it that differed in only a few atoms was active for the label, suggesting that, at a minimum, PubCheF captures molecular function with high fidelity at the scaffold level. The full PubCheF dataset is available in **Supplementary File S2** (available for download at https://doi.org/10.5281/zenodo.21108754).

### Training and evaluating the PubCheF-1 model

While the PubCheF dataset contains known chemical structure-function relationships, we sought to extend these relationships to enable the prediction of function from arbitrary molecular structures. To do so, we fine-tuned a transformer-based chemical language model, ChemBERTa-77M-MLM [32], on the PubCheF dataset to predict function directly from SMILES strings [33] in a multi-label classification setting (**Figure 2B**; see **Materials and methods**). The resulting model, dubbed PubCheF-1, substantially outperformed an RDKit [34] fingerprint-based logistic regression model trained on the same data (validation set PR-AUC: 0.57 vs 0.30; **Supplementary Figure S3A**). An ablation study of PubCheF-1 showed that masked language model pre-training, focal loss [35] instead of binary cross entropy loss, and ensembling across three independently trained models each improved the macro-averaged validation-set PR-AUC (**Supplementary Figure S3A**).

We created two chemical function benchmark datasets to evaluate PubCheF-1 against existing molecular annotation models: PubCheF-test-reduced (10.3K molecules) and OpenTargets-indications [36] (6.7K molecules), both of which are provided in **Supplementary File S1**. Existing annotation models differ substantially in their prediction formats: the CheF classifier [11] predicts functional labels using a different vocabulary from the one used by PubCheF-1, whereas MolT5 [37] generates free-text descriptions autoregressively. These differences make deterministic comparisons of functional accuracy between predictors difficult to perform at scale. We resolved this by creating an evaluation method that uses a judge LLM [38] to automatically score the confusion matrix, from either free-text output or thresholded class predictions, upon which metrics like precision can be calculated (see **Materials and methods**). This approach showed strong agreement with human evaluation on a set of 300 predictions spanning all three models and both benchmark datasets (row-wise F1: Pearson r=0.89, mean absolute difference 0.03; **Supplementary Table S4**, **Supplementary File S1**).

Using this evaluation method, PubCheF-1 strongly outperformed CheF and MolT5 on both benchmark datasets, achieving an F1 score of 0.53 on PubCheF-test-reduced versus 0.15 and 0.05 for MolT5 and CheF, and 0.12 on OpenTargets-indications versus 0.05 and 0.07. On OpenTargets-indications, F1 was low for all models because many of its molecules carry hundreds of indications that inflated the occurrence of false negatives. When evaluating by hit-rate and macro-precision, which avoid this penalty, PubCheF-1 again ranked first (0.30 and 0.17, versus ≤0.18 and ≤0.08 for the other models; **Supplementary Table S5**, **Supplementary File S1**).

To determine whether PubCheF-1’s performance results from its dataset rather than its architecture or training scale, we trained an identical model (same architecture, hyperparameters, and three-seed ensemble) on a bioactivity dataset derived from ChEMBL 33 [9] (see **Materials and methods**). This dataset is of similar size to PubCheF (1.33M molecules and 6,588 targets) and shares 52% of PubCheF’s molecules and 62% of its scaffolds. PubCheF-1 outperformed the ChEMBL-based model on both benchmarks, more than tripling its F1 score (0.530 vs 0.158 on PubCheF-test-reduced and 0.115 vs 0.033 on OpenTargets-indications; **Supplementary Table S5**). Given the high overlap of molecular structures between PubCheF and the ChEMBL-derived dataset, this difference suggests that the supervision signal from literature-derived annotations is advantageous for predicting broad biological function.

Given the high performance of PubCheF-1, we set out to understand the mechanisms by which the model assigns function. First, we found that individual label performance of the non-ensembled PubCheF-1 model improved when trained on all labels jointly rather than on each label in isolation (n=10 random labels; **Supplementary Figure S3B**). When we visualized the PubCheF dataset labels based on their connectivity to other labels sharing the same molecules, an intuitive semantic structure became apparent (**Figure 2C**). This suggests that the language we use to describe biological effects is chemically coherent, and hence the label co-occurrences for a given molecule provide useful context that allows the model to predict function more accurately. Consistent with this interpretation, the weights of the classifier layer were correlated with co-occurrence for the most common labels; for example, the top label, “Anticancer”, had a 0.64 Mantel test correlation (**Supplementary Figure S3C**). Next, we performed an input ablation study, retraining the non-ensembled PubCheF-1 model on progressively corrupted inputs, to understand which input features drive its predictions (**Supplementary Figure S3D**, see **Materials and methods**). Removing chirality from the inputs before training had little impact on validation set performance (0.56 PR-AUC achiral input vs. 0.56 PR-AUC chiral). Shuffling the token order resulted in a model that retained some, though significantly reduced, predictive capacity (0.17 PR-AUC). Randomizing the embeddings into the classifier layer caused the model to be non-functional (0.00 PR-AUC). The residual performance following token shuffling suggests that the composition of SMILES strings, which includes tokens for bonds, branch points, and ring closures, is distinct enough for a model to estimate a molecule’s location in chemical space without explicit ordering. Consistent with this, we found that 84.5% of the molecules in PubCheF had unique token count vectors. We then tested the extent to which these count vectors could be used to estimate location in chemical space by computing their correlation to fingerprint Tanimoto distances. We computed all-by-all Tanimoto distance matrices for both RDKit fingerprints and token count vectors across the 10,311-molecule PubCheF-test-reduced set, and a Mantel test between the two yielded a Pearson correlation of 0.46 (p=0.0001), in agreement with existing literature [39]. Thus, while token composition positions molecules only approximately within chemical space, their explicit structural ordering is necessary for accurate function prediction, as evidenced by the performance increase from 0.17 to 0.56 PR-AUC (**Supplementary Figure S3D**). Finally, we used layer-wise relevance propagation (LRP) [40] to attribute each prediction to the input atom tokens, quantifying their importance for predicting a given function. Anecdotal examples suggest that PubCheF-1 has converged upon known structure-function relationships for SARS-CoV-2 protease inhibition (**Figure 2D**), 5-HT2A receptor agonism, and prodrug modifications (**Supplementary Figures S3E-G**). More systematically, we found that across the PubCheF-1 test set the model preferentially used large, connected substructures enriched in heteroatoms to predict labels (**Supplementary Figure S3H-J**).

### *In silico* hit discovery with PubCheF-1

Having established that PubCheF-1 can predict function from structure, often through chemically meaningful relationships, we next assessed whether its predictions could be used for prospective bioactive molecule discovery. As an initial *in silico* assessment, we explored its ability to predict inhibitors for five different protein targets: KPC-3 β-lactamase, 5-HT2A receptor, SARS-CoV-2 main protease (Mpro), Taq DNA polymerase (DNAP), and Hepatitis C Virus NS5B RNA polymerase. These targets were chosen based on their differing mechanisms of action and varying abundances and representation in the PubCheF-1 vocabulary. First, we generated PubCheF-1 predictions of chemical function across the entire ZINC20 in-stock dataset of 13.2 million molecules [41] and created a website to visualize these predictions (see https://www.pubchef.org). We then selected molecules that had high scores for labels relevant to each of the five targets, creating two sets of 100 molecules for each target: a set of the top 100 highest-scoring molecules, and a set of the top 100 highest-scoring “novel” molecules, determined by having less than 0.50 RDKit fingerprint Tanimoto similarity to any molecules in the PubCheF dataset that were annotated with the selected-for labels. For the top and novel arms, we obtained Boltz-2 affinity probabilities for each of the hit molecule sets to their respective targets (called “on-target hits”), each hit molecule set to the other targets (called “off-target hits”), and for a set of 100 randomly chosen ZINC20 available molecules (called “random ZINC”). Among the top-ranked hits, all five targets scored significantly higher for their on-target hits compared to the off-target hits (KPC-3: 0.60 vs 0.30, 5-HT2A: 0.69 vs 0.18, Mpro: 0.72 vs 0.31, DNAP: 0.46 vs 0.36, and NS5B: 0.41 vs 0.35; BH-corrected Mann-Whitney p ≤ 2.7×10⁻⁴ for all comparisons), and to the random ZINC molecules (p ≤ 1.6E-14) (**Figure 2E**). Among the top-ranked novel hits, four of the five targets scored significantly higher for the on-target hits compared to the off-target hits (KPC-3: 0.52 vs 0.33, 5-HT2A: 0.63 vs 0.15, Mpro: 0.30 vs 0.23, and DNAP: 0.38 vs 0.32; BH-corrected Mann-Whitney p ≤ 2.7×10⁻³ for all comparisons), and to the random ZINC molecules (p ≤ 1.4E-7) (**Figure 2F**). NS5B was the only target for which the novelty-filtered hit set failed to show significant enrichment, likely because its much larger representation in PubCheF (7,765 molecules versus ≤2,306 for the other targets) caused the fingerprint-based novelty filter to exclude many otherwise plausible candidate scaffolds. Overall, PubCheF-1’s agreement with the structure-based Boltz-2 model suggests that it may be capable of identifying bioactive molecules across a broad range of biological functions, potentially including inhibitors missed by Boltz-2.

### Predicting β-lactamase inhibitors for experimental validation

To understand if PubCheF-1’s predictions are meaningful in a real-world biological context, we focused on the discovery of β-lactamase inhibitors. β-lactamases comprise multiple enzyme classes with distinct catalytic mechanisms, many of which remain poorly addressed by currently available inhibitors [15,20,26]. Moreover, the continued evolution and dissemination of β-lactamases under antibiotic selection pressure [19–22] creates an ongoing need for new resistance-breaking therapeutics. Finally, our *in silico* analyses suggested that β-lactamase inhibitor discovery represented a suitable prospective test of PubCheF-1. The model performed well on KPC-3 inhibitor prediction (**Figure 2E,F**), and β-lactamase-related functional labels were represented by approximately 2,300 molecules in the PubCheF dataset, providing sufficient training signal for meaningful prediction.

We used PubCheF-1 to predict structurally distinct and readily purchasable candidate β-lactamase inhibitors (**Figure 2H**). From the above ZINC20 in-stock predictions, we identified 30,267 compounds relevant to β-lactamase inhibition as those having ≥0.20 PubCheF-1 score on at least one label containing the substring “lactam” (corresponding to 0.33 precision and 0.65 recall in the test set at this cutoff). Then, following a prioritization strategy similar to that used by Wong et al. [42], we filtered for structural novelty (≤0.50 RDKit fingerprint Tanimoto similarity to known β-lactamase inhibitors in PubCheF), low predicted toxicity to HepG2, HSkMC, and IMR-90 cells (<0.20 for each model in [42]), absence of Pan-Assay Interference Substructures [43], and availability from preferred vendors. This resulted in 3,880 candidate molecules (**Supplementary File S3**; see **Materials and Methods**). From this set, and with a budget of $4,000, we selected and purchased 111 compounds, optimizing for low cost, structural diversity, high PubCheF-1 score, and minimal Brenk flags [44]. For selected PubCheF-1-predicted compounds that were unavailable for purchase, the closest commercially available analogs were ordered instead.

### Screening predicted β-lactamase inhibitors

To experimentally evaluate the predicted inhibitor compounds, we first established a protein target screening platform (**Figure 3A**) using diverse, purified β-lactamase enzymes (**Supplementary Figure S4A**). Specifically, we selected the class A carbapenemase KPC-3, the class B3 metallo-β-lactamase L1_#3637_, the class C cephalosporinase EC-2, and the class D oxacillinase OXA-10 (**Supplementary Tables S6** and **S7**). These clinically important enzymes represent all four major structural Ambler classes of β-lactamases and include targets for which effective inhibitors remain limited, like the metallo-β-lactamase L1_#3637_ and the oxacillinase OXA-10 [15,24–27]. Purified proteins were incubated with nitrocefin, a chromogenic β-lactam substrate [45], in the presence of individual PubCheF-1-predicted compounds, and inhibition was quantified by monitoring nitrocefin hydrolysis relative to carrier controls (**Supplementary File S4**). Candidate hits were defined as compounds producing ≥20% inhibition. This threshold is consistent with previous early-stage discovery campaigns and nitrocefin-based β-lactamase inhibitor screens, which commonly employ relatively permissive activity cutoffs (8-15% [46–49]). Compounds were only considered hits if the observed inhibition was also statistically significant relative to the corresponding carrier control.

**Figure 3.**
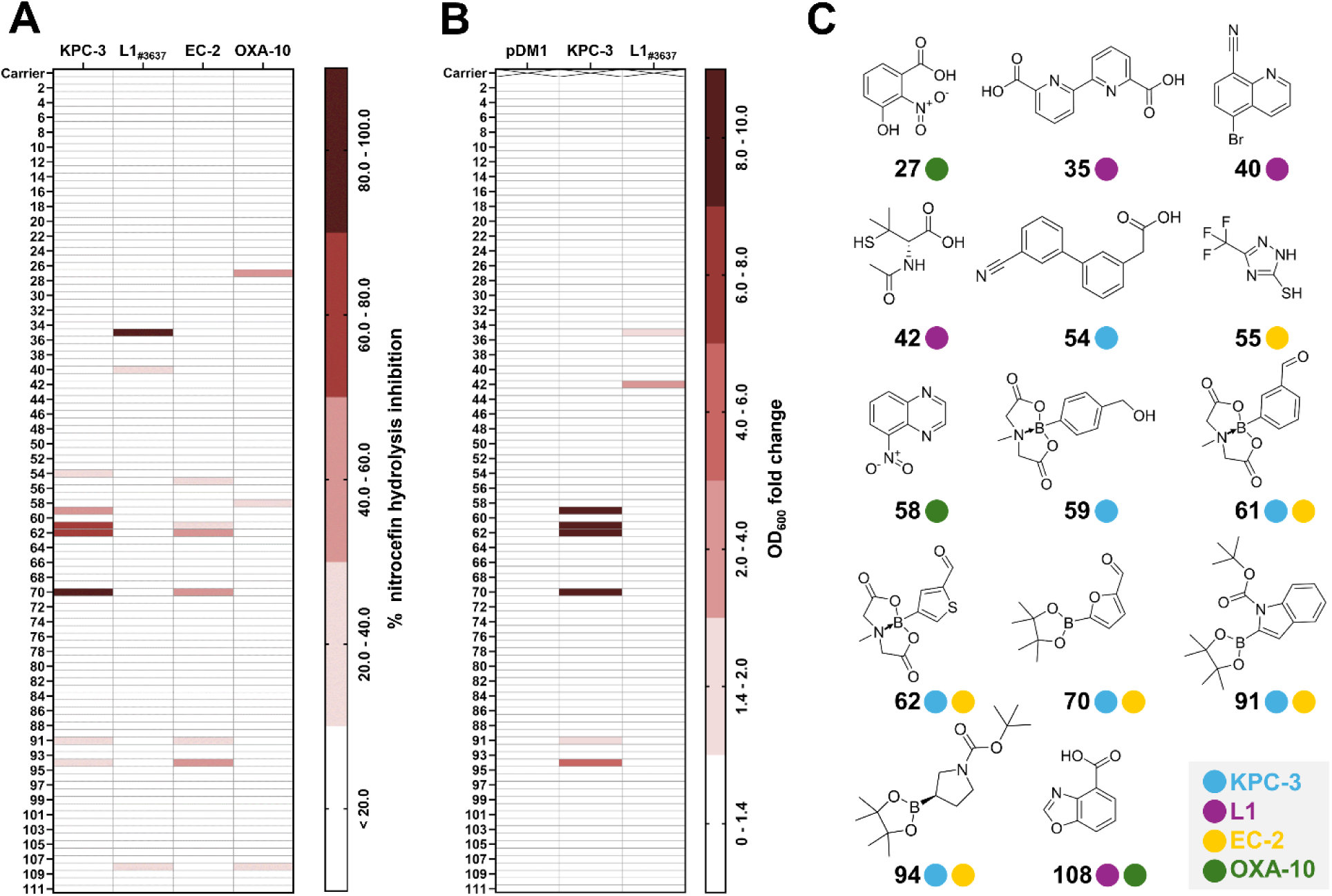
Protein target and in-bacteria screening of the PubCheF-1 candidate compounds identifies molecules with inhibitory activity against β-lactamase enzymes. **(A)** 13 of the 110 tested compounds predicted by PubCheF-1 (compound #90 arrived damaged and was not tested) caused a significant reduction in β-lactamase activity, corresponding to an overall screening hit rate of 11.8%; the most active compounds achieved >80% reduction in nitrocefin hydrolysis by purified and phylogenetically diverse β-lactamase enzymes. Data shown represent the percentage reduction of nitrocefin hydrolysis in a reaction containing the candidate compound compared to a reaction containing the carrier control. Four representative β-lactamases were used: the class A serine-β-lactamase KPC-3, the class B metallo-β-lactamase L1_#3637_ (harboring a C-terminal StrepII tag), the class C serine-β-lactamase EC-2 (as an N-terminal fusion with a His_8_-SUMO solubility tag), and the class D serine-β-lactamase OXA-10 (harboring a C-terminal StrepII tag). Hits are defined as candidate compounds that cause statistically significant activity loss ≥20%. Inhibition strength is signified by color intensity; the darker the red color the greater enzyme inhibition, while compounds resulting in no significant reduction in enzyme activity are shown in white. Statistical analysis was performed in GraphPad Prism v10.6.1 using a one-way ANOVA and significance was defined as p < 0.05. Inhibitor hits include: #54 (p<0.0001; significance, ****), #59 (p<0.0001; significance, ****), #61 (p<0.0001; significance, ****), #62 (p<0.0001; significance, ****), #70 (p<0.0001; significance, ****), #91 (p=0.0009; Significance, ***), and #94 (p<0.0001; significance, ****) for KPC-3; #35 (p<0.0001; significance, ****), #40 (p<0.0001; significance, ****), and #108 (p=0.0002; significance, ***) for L1_#3637_; #55 (p=0.0005; significance, ***), #61 (p<0.0001; significance, ****), #62 (p<0.0001; significance, ****), #70 (p<0.0001; significance, ****), #91 (p<0.0001; significance, ****), and #94 (p<0.0001; significance, ****) for EC-2; #27 (p<0.0001; significance, ****), #58 (p<0.0001; significance, ****), and #108 (p<0.0001; significance, ****) for OXA-10. Raw and analyzed data and statistical analyses can be found in **Supplementary File S4**. **(B)** 8 of the 110 tested compounds predicted by PubCheF-1 caused a significant reduction in bacterial survival of *E. coli* MC1000 (K-12 strain) expressing either the serine-β-lactamase KPC-3 (middle column) or the metallo-β-lactamase L1_#3637_ (right column), when exposed to sub-MIC levels of the β-lactam antibiotic ceftazidime; no candidate compounds showed bactericidal activity against the empty vector control (pDM1; left column). Molecules with activity in this screen include all of the hits identified in the protein target screen shown in (A), except for compound #54 (KPC-3 hit) and compound #40 (L1_#3637_ hit) that likely cannot access the Gram-negative periplasm. Compound #42 was not a hit in the protein target screen but exhibited inhibitory activity against L1_#3637_-expressing bacteria. Neither the empty-vector control nor the carrier control for compound #42 showed bacterial killing (**Supplementary Figure S5G-I**), therefore it is unlikely that the observed activity is due to off-target effects. Together, the target and in-bacteria screens yielded a total hit rate of 12.7% (14 compounds). Data shown represent the fold change of OD_600_ of cells exposed to 32 µg/mL of the candidate compounds in the presence of ∼0.3x MIC of ceftazidime for each strain over the OD_600_ of cells exposed only to ∼0.3x MIC of ceftazidime (48 µg/mL ceftazidime for KPC-3 or L1_#3637_ and 0.01 µg/mL for pDM1; relevant MIC values are shown in **Supplementary Figure S4B**). The schematic of the experimental setup is available in **Supplementary Figure S4C**. Hits are defined as candidate compounds that cause a statistically significant fold change difference greater than 1.4. This corresponds to reproducible reduction in OD_600_ above the noise of the pDM1 empty vector results; carrier controls are shown in **Supplementary Figure S4D**. Inhibition strength is indicated by color intensity; the darker the red color the greater the decrease in optical density, while compounds resulting in no change in bacterial survival are shown in white. Statistical analysis was performed in GraphPad Prism v10.6.1 using a one-way ANOVA and significance was defined as p < 0.05. Inhibitor hits include: #59 (p<0.0001; significance, ****), #61 (p<0.0001; significance, ****), #62 (p<0.0001; significance, ****), #70 (p<0.0001; significance, ****), #91 (p=0.0031; significance, **), #94 (p<0.0001; significance, ****), #35 (p<0.0001; significance, ****), and #42 (p<0.0001; significance, ****). Raw and analyzed data and statistical analyses can be found in **Supplementary File S5**. **(C)** Chemical structures of candidate compounds exhibiting activity against class A KPC-3 (blue), class B L1_#3637_ (purple), class C EC-2 (yellow), and class D OXA-10 (green) β-lactamases; compounds identified from both the target and in-bacteria screens are shown (ChemDraw 23.1.2). Notably, PubCheF-1 identified three monosubstituted organoboron compounds (indicated by #70*, #91*, and #94* and shown in **Supplementary Figure S4E**). Because these non-cyclized analogs were not commercially available, the closest structurally related compounds, the cyclized analogs #70, #91, and #94, were selected for purchase and experimental testing instead.

The protein target screen identified 13 unique candidate hits (**Figure 3A**). Seven compounds inhibited KPC-3 (#54, #59, #61, #62, #70, #91, and #94), three inhibited L1_#3637_ (#35, #40, and #108), six inhibited EC-2 (#55, #61, #62, #70, #91, and #94), and three inhibited OXA-10 (#27, #58, and #108). Notably, the distribution of these candidate hits reflected known relationships among β-lactamase families. KPC-3 and EC-2, which belong to the more closely related serine β-lactamase classes [50], shared multiple inhibitor hits, whereas the more distant OXA-10 and the mechanistically distinct metallo-β-lactamase L1_#3637_ exhibited more unique inhibition profiles. Consistent with the historical tractability of serine β-lactamases relative to metallo-β-lactamases, the greatest number of hits was recovered against the class A and class C enzymes, followed by the class D and class B enzymes [21]. Together, these results show that PubCheF-1 can successfully identify inhibitors across all four β-lactamase classes, including the enzymes with limited coverage like L1_#3637_ and OXA-10.

Because PubCheF-1 was trained on literature-derived functions rather than biochemical assay measurements, and since many of these functions originate from articles describing resistance breakers in cellular contexts, we hypothesized that compounds identified in the protein target screen would retain activity in more biologically relevant settings. We therefore established an in-bacteria screening platform using isogenic *Escherichia coli* K-12 strains expressing β-lactamases (**Supplementary Tables S7-S10**). We chose to focus on two mechanistically diverse enzymes, the serine β-lactamase KPC-3 that yielded the largest number of high-efficacy hits in the protein target screen and the metallo-β-lactamase L1_#3637_, as a representative of a β-lactamase class with low tractability [24]. Prior to screening, we determined ceftazidime MIC values for the β-lactamase-producing strains and an empty-vector control strain to define antibiotic concentrations that permitted robust bacterial growth while sensitively reporting β-lactamase inhibition (**Supplementary Figure S4B**). Candidate compounds were then screened in the presence of sub-inhibitory ceftazidime concentrations (∼0.3x of the MIC value of each strain), and activity was assessed by measuring compound-dependent reductions in bacterial growth relative to carrier controls (**Supplementary Figure S4C, Supplementary File S5**). Control experiments confirmed that neither the compound carriers nor the screening conditions significantly affected bacterial survival (**Supplementary Figure S4D**).

The in-bacteria screen identified eight unique compounds that potentiated β-lactam activity against β-lactamase-producing strains (**Figure 3B**). Importantly, only three candidate hits from the protein target screen failed to retain activity in the cellular assay (#40, #54, and #108), demonstrating a low attrition rate despite the additional permeability, stability, and target-engagement requirements imposed by the bacterial environment. Conversely, the in-bacteria screen identified an additional L1_#3637_ candidate hit (#42) that exhibited no detectable inhibition in the purified protein assay (**Figure 3A**) yet was subsequently validated as a bona fide L1 enzyme inhibitor during follow-up studies (see **Figures 4** and **5**). Collectively, these results demonstrate that PubCheF-1 enriched both for compounds capable of inhibiting purified β-lactamases across all four Ambler classes and for compounds that retain activity in living cells. The resulting hit set encompassed multiple chemical scaffolds and included several boron-containing compounds that among others were selected for further mechanistic characterization (**Figure 3C**).

**Figure 4.**
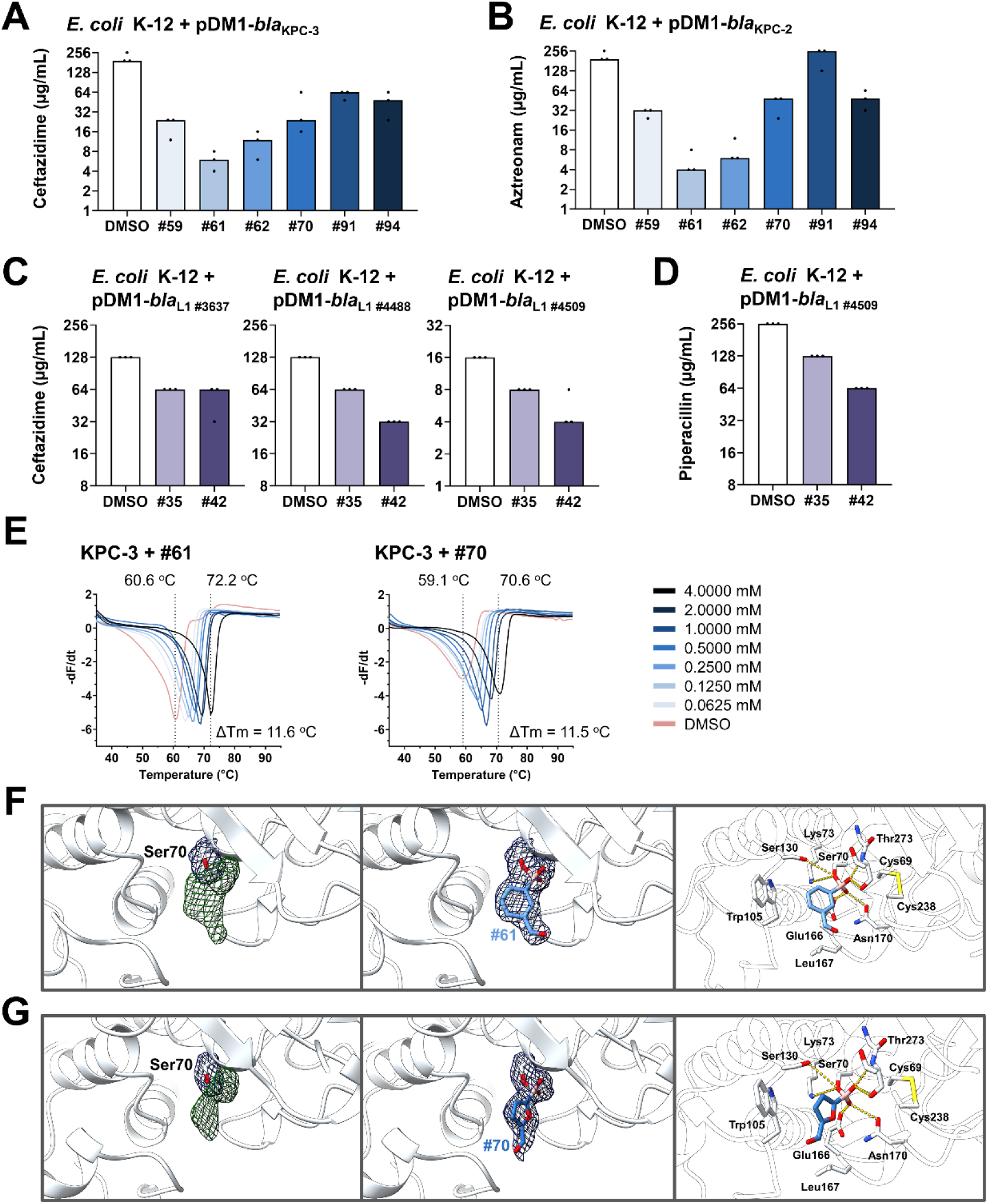
(A-D) Candidate compound hits reduce β-lactam resistance in *E. coli* K-12 expressing serine and metallo-β-lactamase enzymes. **(A)** Ceftazidime MIC values for *E. coli* MC1000 (K-12 strain) expressing the class A serine β-lactamase KPC-3 are reduced in the presence of all hits identified in **Figure 3B** (final concentration of 32 µg/mL; blue bars) in comparison to the DMSO carrier control (added at an equivalent volume; white bar). MIC values remained unchanged for *E. coli* MC1000 when exposed to high concentrations of the inhibitor hits (up to 512 µg/mL, **Supplementary Figure S5A**), and the empty vector (pDM1) control when subjected to a combination of the hits with the non-β-lactam antibiotic gentamicin (**Supplementary Figure S5B**), or a combination of the hits and the β-lactam ceftazidime (**Supplementary Figure S5C**). Furthermore, the gentamicin MIC for *E. coli* MC1000 expressing KPC-3 (pDM1-*bla*_KPC-3_, **Supplementary Figure S5E**) also remained unchanged. Together, these data show that inhibitory activity is specific to the presence of the β-lactamase and not due to off-target bactericidal effects. **(B)** KPC-2 is a prevalent member of the KPC family that differs from KPC-3 by only one amino acid at position 272 but exhibits distinct activity against β-lactam antibiotics [52–54]. Aztreonam MIC values for *E. coli* MC1000 expressing KPC-2 show that, with the exception of compound #91, the inhibitory activity of the KPC-3 inhibitor hits is retained. This activity is confirmed not to be due to off-target effects as in (A) (**Supplementary Figure S5B**, **Supplementary Figure S5D**, and **Supplementary Figure S5F**). **(C)** Ceftazidime MIC values for *E. coli* MC1000 expressing the class B metallo-β-lactamase L1 variants L1_#3637_ (**left**, from *S. maltophilia* ATCC 13637), L1_#4488_ (**middle**, from *S. maltophilia* ERR1974488), and L1_#4509_ (**right**, from *S. maltophilia* ERR1974509) are reduced in the presence of all hits identified in Figure 3B (final concentration of 32 µg/mL; purple bars) in comparison to the DMSO carrier control (added at an equivalent volume; white bar). This activity is confirmed not to be due to off-target effects as in (A) and (B) (**Supplementary Figure S5G**, **Supplementary Figure S5H**, **Supplementary Figure S5I**, and **Supplementary Figure S5K-M**). **(D)** Piperacillin MIC values for *E. coli* MC1000 expressing L1_#4509_ are reduced in the presence of all hits identified in Figure 3B (final concentration of 32 µg/mL; purple bars) in comparison to the DMSO carrier control (added at an equivalent volume; white bar). This activity is confirmed not to be due to off-target effects as in (A-C) (**Supplementary Figure S5G**, **Supplementary Figure S5H**, **Supplementary Figure S5J**, and **Supplementary Figure S5M**). For panels (A-D), graphs show MIC values (μg/mL) from three biological experiments, each conducted as a single technical repeat. Raw MIC data are available in **Supplementary File S6B**. Consistent with convention, error bars and significance assessment are not shown for MIC panels, as MIC values are discrete. **(E-G) Hits #61 and #70 inhibit KPC-3 by binding covalently to the active-site serine residue. (E)** Thermal shift assays show that binding of inhibitor hits #61 and #70 stabilizes KPC-3. First-derivative melting curves (-dF/dT) of purified KPC-3 in the presence of increasing concentrations of the inhibitor hits #61 and #70; both inhibitors induced a concentration-dependent increase in melting temperature (Tₘ), with a maximum ΔTₘ of ∼11.5 °C at 4 mM. X-ray structures of KPC-3·in complex with inhibitor hit **(F)** #61 and **(G)** #70: **(left)** Fo–Fc omit map (green mesh, contoured at +3σ) calculated before ligand modelling. **(middle)** 2Fo–Fc map (grey mesh, contoured at 1.0σ) after ligand fitting. **(right)** Detailed view of ligand-protein interactions; the ligand is shown in blue sticks, the boron atom is colored pink, and key hydrogen bonds are depicted as yellow dashed lines.

**Figure 5.**
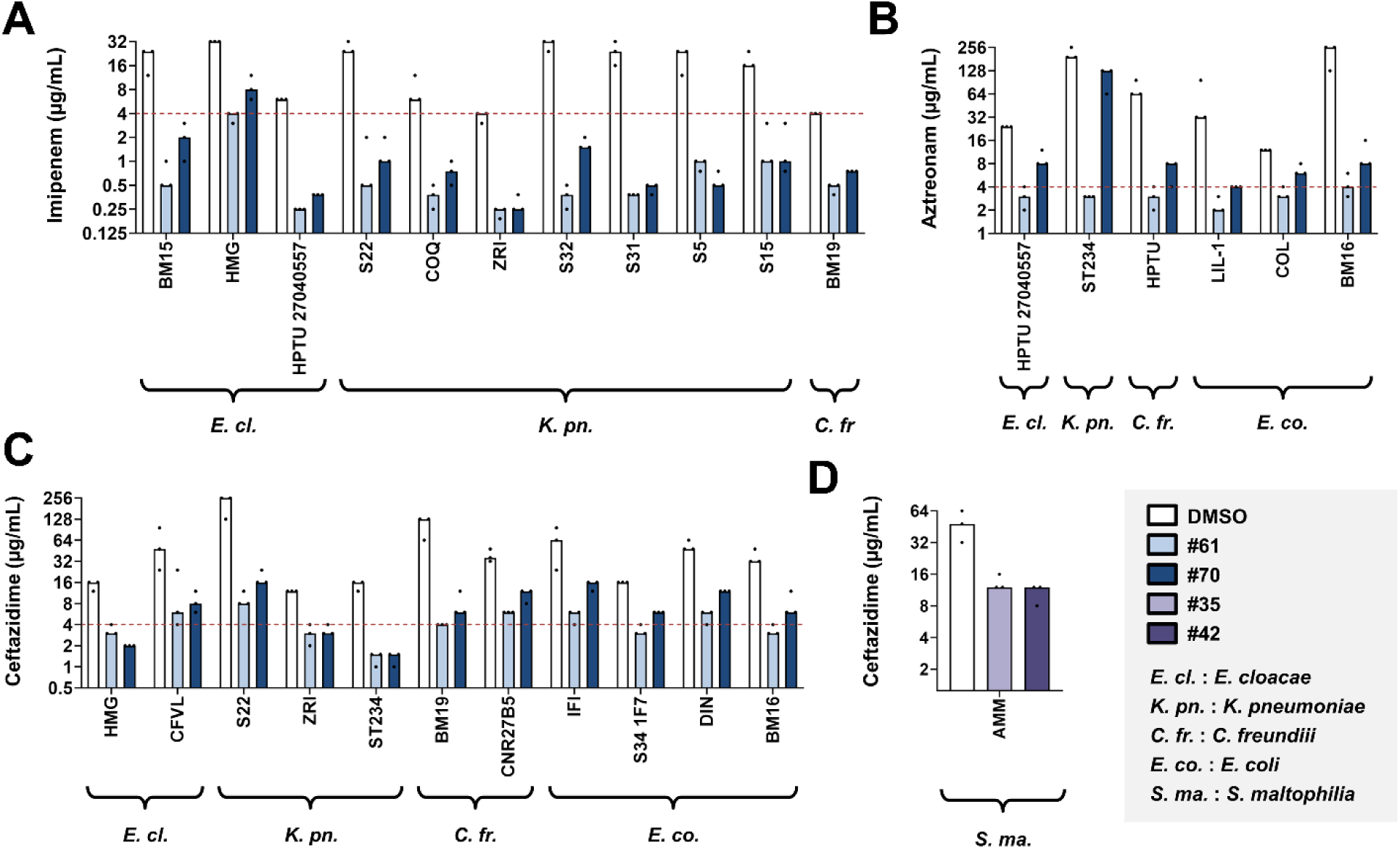
Inhibitor hits sensitize multidrug-resistant clinical isolates to existing β-lactam antibiotics. Inhibitor hits #61 (light blue, 32 µg/mL) and #70 (dark blue, 32 µg/mL) drastically reduce the resistance levels of clinical isolates of *E. cloacae* (*E. cl.*), *K. pneumoniae* (*K. pn.*), *C. freundii* (*C. fr.*), and *E. coli* (*E. co.*) expressing KPC β-lactamases to **(A)** the last-resort β-lactam imipenem, **(B)** the monobactam β-lactam aztreonam, and **(C)** the cephalosporin ceftazidime in comparison to the DMSO carrier control (added at an equivalent volume; white bar). **(D)** Inhibitor hits #35 (light purple, 64 µg/mL) and #42 (dark purple, 64 µg/mL) drastically reduce the resistance levels of a clinical *S. maltophilia* isolate (strain AMM, which we engineered to harbor a deletion of its *bla_L2_* gene,) to ceftazidime in comparison to the DMSO carrier control (added at an equivalent volume; white bar). Graphs show MIC values (μg/mL) from three biological experiments, each conducted as a single technical repeat; red dotted lines indicate the EUCAST clinical breakpoint for each antibiotic, when available. Raw MIC data are available in **Supplementary File S6D,E**. Consistent with convention, error bars and significance assessment are not shown for MIC panels, as MIC values are discrete.

### Validating candidate β-lactamase inhibitor activity

To validate the candidate hits identified in **Figure 3B**, we tested their ability to restore β-lactam activity against *E. coli* K-12 strains expressing β-lactamase enzymes using minimum inhibitory concentration (MIC) assays [51] (**Figure 4A-D**, **Supplementary File S6**). All KPC-3 candidate hits reduced ceftazidime MIC values, with some compounds (#59, #61, #62, and #70) achieving substantial potentiation (**Figure 4A**). To determine whether this activity extended beyond a single KPC enzyme variant, we next evaluated the same compounds against KPC-2, a globally disseminated member of the KPC family that differs from KPC-3 by a single amino acid substitution but exhibits distinct substrate preferences [52–54]. All but one (#91) KPC-3 candidate hits retained activity against KPC-2 and reduced aztreonam MIC values (**Figure 4B**). These results show that the identified compounds target conserved features of KPC-family enzymes rather than a single β-lactamase variant.

We next evaluated inhibitors identified against the metallo-β-lactamase L1_#3637_. We tested the activity of candidate hits #35 and #42 against three enzymes (L1_#3637_ and L1_#4488_ and L1_#4509_) originating from distinct *Stenotrophomonas maltophilia* isolates expressed in *E. coli* K-12. All identified compounds reduced ceftazidime MIC values across the tested variants (**Figure 4C**). Inhibitory activity was also retained when tested using the structurally distinct β-lactam piperacillin (**Figure 4D**), demonstrating efficacy across both enzyme variants and antibiotic substrates.

To determine whether the observed MIC reductions were specifically associated with β-lactamase inhibition, we performed an extensive series of control experiments (**Supplementary Figure S5, Supplementary File S6**). None of the compounds exhibited intrinsic antibacterial activity at the highest tested concentrations (**Supplementary Figure S5A,G**). Similarly, the compounds did not alter susceptibility of empty-vector control strains to either β-lactam antibiotics or the non-β-lactam gentamicin (**Supplementary Figure S5B-D,H-J**), nor did they affect gentamicin susceptibility in β-lactamase-producing strains (**Supplementary Figure S5E-F,K-M**). Together, these results show that the observed potentiation of β-lactam activity requires the presence of the target β-lactamase and is not attributable to general toxicity or increased sensitivity to general antibiotic stress.

### Determining the structural basis of β-lactamase inhibition

Given their potent inhibition against KPC-family β-lactamases (**Figure 4A,B**) and distinct chemical scaffolds (**Figure 3C**), we pursued further biophysical and structural characterization of inhibitor hits #61 and #70. To assess direct interaction with KPC-3, we first examined their effects on enzyme stability using differential scanning fluorimetry (DSF). Incubation of KPC-3 with increasing concentrations of compounds #61 and #70 resulted in clear concentration-dependent increases in melting temperature (T_m_), with maximal thermal shifts (ΔT_m_) of 11.5 °C observed at 4 mM for both compounds (**Figure 4E**). These substantial ligand-induced thermal shifts indicated direct engagement of KPC-3 by both inhibitor hits and motivated subsequent structural characterization.

To elucidate the inhibition mechanism of these compounds at the atomic level, we determined the X-ray crystal structures of KPC-3 in complex with inhibitor hits #61 and #70. As a reference for comparison, we began by solving the apo structure of KPC-3 to define the unliganded active site. The enzyme crystallized in the space group *P3₁* and diffracted to a resolution of 2.40 Å (**Supplementary Table S11**). Each asymmetric unit contained three protein molecules, although the enzyme is monomeric in solution, as confirmed by gel filtration (**Supplementary Figure S6**). The active site was well resolved, including a clearly defined disulfide bond between Cys69 and Cys238 (**Supplementary Figure S7A**). A Fo-Fc omit map calculated from the protein model alone revealed residual electron density corresponding to a bicine molecule from the crystallization buffer positioned adjacent to the active site (**Supplementary Figure S7B**). This observation is consistent with a previously reported KPC-2 structure (PDB 2OV5) [55], in which a carboxyl moiety occupies the same position as the β-lactam carboxylate group during substrate binding.

To obtain the inhibitor-bound complexes, apo KPC-3 crystals were soaked with compound #61 (1 mM) or #70 (10 mM) for varying durations in bicine-free buffer. The resulting crystals diffracted to 2.5 Å and 2.3 Å resolution, respectively (**Supplementary Table S11**). While additional electron densities were evident in the active sites of all three molecules, ligands were modeled only in molecules exhibiting the clearest density (**Figure 4F,G**). The refined ligands exhibited high real-space correlation coefficients (RSCC = 0.98–0.99) and B-factors comparable to the surrounding residues, consistent with high ligand occupancy (**Supplementary Table S12**).

In both complexes, clear positive Fo-Fc difference density was observed contiguous with the Oγ atom of the catalytic nucleophile Ser70 (**Figure 4F,G**). This density was absent in the apo structure and was consistent with covalent attachment of the inhibitor. The electron density unambiguously revealed a boron atom in an sp³ configuration, forming a tetrahedral boronate adduct covalently linked to Ser70 Oγ with a bond distance of 1.5 Å (**Figure 4F,G**). Thus, the structures are consistent with nucleophilic attack of Ser70 on the electrophilic boron center, yielding a covalent enzyme-inhibitor adduct. Notably, no residual density corresponding to the boron-protecting groups was observed, indicating that both compounds undergo deprotection prior to adduct formation. The boronate adduct is stabilized by a conserved hydrogen-bonding network typical of Class A β-lactamases (**Figure 4F,G**). One boronate hydroxyl occupies the oxyanion hole formed by the backbone amides of Ser70 and Thr237, whereas the second is stabilized through interactions with Asn170 and Glu166, consistent with previously described boronic acid transition-state analogs in KPC enzymes [55].

Beyond the covalent linkage, the overall ligand densities were highly similar between the complex of KPC-3 with #61 and #70, reflecting their shared scaffold. The principal structural difference involves the ring system adjacent to the boronate moiety: compound #61 has a phenyl group, whereas compound #70 features a furan. In both structures, an aldehyde substituent projects from the ring system and adopts a planar geometry consistent with extended π-conjugation, stabilizes the electron distribution and increases the electrophilicity of the boron center. The aromatic portion of the inhibitors is positioned against a hydrophobic pocket formed by Trp105 and Leu167, providing additional interactions that likely contribute to ligand binding and orientation within the active site (**Figure 4F,G**).

Since the crystal structures of KPC-3 in complex with the inhibitor hits #61 and #70 revealed that the boron-protecting groups of both compounds were absent in the final enzyme-inhibitor hit complexes, we were prompted to investigate whether deprotection occurs spontaneously in solution or within the enzyme active site. We analyzed compounds #61 and #70 by ^1^H NMR spectroscopy in anhydrous deuterated dimethyl sulfoxide (DMSO-*d*_6_) and in a 50:50 mixture of DMSO-*d*_6_ and D₂O at 37 °C (**Supplementary Figure S8**). In anhydrous DMSO-*d*_6_, both compounds remained fully intact, with spectra consistent with the protected molecules (**Supplementary Figure S8A,B**). Upon exposure of compound #61 to aqueous media, however, the NMR spectra changed over time and hydrolysis of the boron-protecting group was observed (**Supplementary Figure S8C**). Approximately 60% of the starting material had been converted to the corresponding boronic acid after 2 hours (**Supplementary Figure S8D**). In contrast, compound #70 remained stable in the presence of D₂O, showing no evidence of hydrolysis even 24 hours after dissolution (**Supplementary Figure S8E**). These findings indicate that compounds #61 and #70 achieve β-lactamase inhibition through distinct activation mechanisms. Compound #61 functions as a prodrug that undergoes spontaneous hydrolysis in aqueous solution, whereas compound #70 remains protected prior to enzyme binding and likely undergoes deprotection during the formation of the enzyme-inhibitor complex.

### Interpreting PubCheF-1 inhibitor predictions

Although the predicted compounds were selected using generic β-lactamase-related labels, retrospective analysis of the obtained inhibitor hits revealed that PubCheF-1 correctly distinguished between serine- and metallo-β-lactamase inhibition profiles and could have been used to selectively enrich for either one in a forward context (**Supplementary Figure S9**). We then examined the structural features underlying PubCheF-1 predictions using layer-wise relevance propagation. Across the validated inhibitor hits, the model consistently assigned importance to discrete molecular substructures rather than entire chemical scaffolds (**Supplementary Figure S10**). For example, in compounds #61 and #70, the two KPC-family inhibitors selected for structural characterization, the boronate-containing portions of the molecules were highly important for predicting the β-lactamase-related labels. Consistent with these predictions, our structural analyses (**Figure 4F,G**) revealed direct participation of these groups in covalent enzyme binding. These analyses indicate that PubCheF-1 uses chemically meaningful determinants of inhibition to capture distinctions within the broader β-lactamase inhibitor landscape.

To compare our literature-based approach with a structure-based one, we evaluated the ability of PubCheF-1 and Boltz-2 [5] to recover the experimentally validated β-lactamase inhibitor hits from a pool containing confirmed negatives and presumed decoys (**Supplementary Figure S11**). Although Boltz-2 successfully identified some of the validated inhibitors, PubCheF-1 achieved better performance without explicitly modeling the target and at substantially reduced computational cost. This suggests that the structure-function relationships contained in scientific literature may provide a complementary role to structure-based methods.

### Restoring β-lactam efficacy in clinical isolates

To determine whether the validated inhibitor hits retained activity in clinically relevant settings, we evaluated their ability to sensitize multidrug-resistant clinical isolates to existing β-lactam antibiotics (**Figure 5**, **Supplementary File S6**, **Supplementary Table S8**). We focused on compounds #61 and #70, which displayed the strongest activity against KPC-family β-lactamases (**Figure 4A,B**) and compounds #35 and #42, which inhibited L1 metallo-β-lactamases (**Figure 4C,D**).

Compounds #61 and #70 substantially reduced the MIC values of a diverse panel of KPC-producing clinical isolates (*Enterobacter cloacae, Klebsiella pneumoniae, Citrobacter freundii,* and *E. coli*) to diverse β-lactams, including the last-resort carbapenem imipenem and the monobactam aztreonam (**Figure 5A-C**). Notably, some of these clinical strains harbor complex resistance backgrounds that extend beyond a single β-lactamase. Despite the presence of multiple enzymes, the inhibitor hits consistently reduced resistance levels across species and antibiotic classes. In many cases, inhibitor treatment drastically reduced MIC values to below established EUCAST clinical breakpoints, sensitizing the strains to existing antibiotics. To determine whether potentiation was retained at lower inhibitor concentrations, we evaluated representative strain-antibiotic combinations using decreasing concentrations of compounds #61 and #70 (**Supplementary Figure S12A-C**). Both compounds maintained their activity across a broad concentration range. In some cases, potentiation was retained at inhibitor concentrations as low as 2 μg/mL, whereas in others activity persisted across intermediate concentrations before declining at the lower tested doses.

Metallo-β-lactamases remain particularly challenging therapeutic targets because clinically available inhibitors are lacking. We therefore sought to determine whether the L1 inhibitor hits retained activity in the enzyme’s native clinical host. *S. maltophilia* is an extensively drug-resistant emerging pathogen, which is recalcitrant to most β-lactam antibiotics since its L1 and L2 enzymes contribute complementary β-lactam resistance phenotypes [56]. We generated a derivative strain in which the serine β-lactamase gene *bla*_L2_ was deleted to enable direct assessment of L1-dependent inhibitor activity. In this genetically defined clinical strain background, compounds #35 and #42 reduced ceftazidime MIC values relative to the carrier control (**Figure 5D**). Together, these findings highlight that the inhibitors identified by PubCheF-1 retain activity in clinically relevant pathogens and remain collectively effective against both serine and metallo-β-lactamase targets.

### Evaluating inhibitor hit efficacy in insect and animal infection models

To determine whether identified inhibitor hits could potentiate β-lactam activity *in vivo*, we evaluated compounds #61 and #70 in two complementary infection models (**Figure 6**, **Supplementary File S7**) using the multidrug-resistant KPC-producing clinical isolate *K. pneumoniae* ZRI (**Supplementary Table S8**).

**Figure 6.**
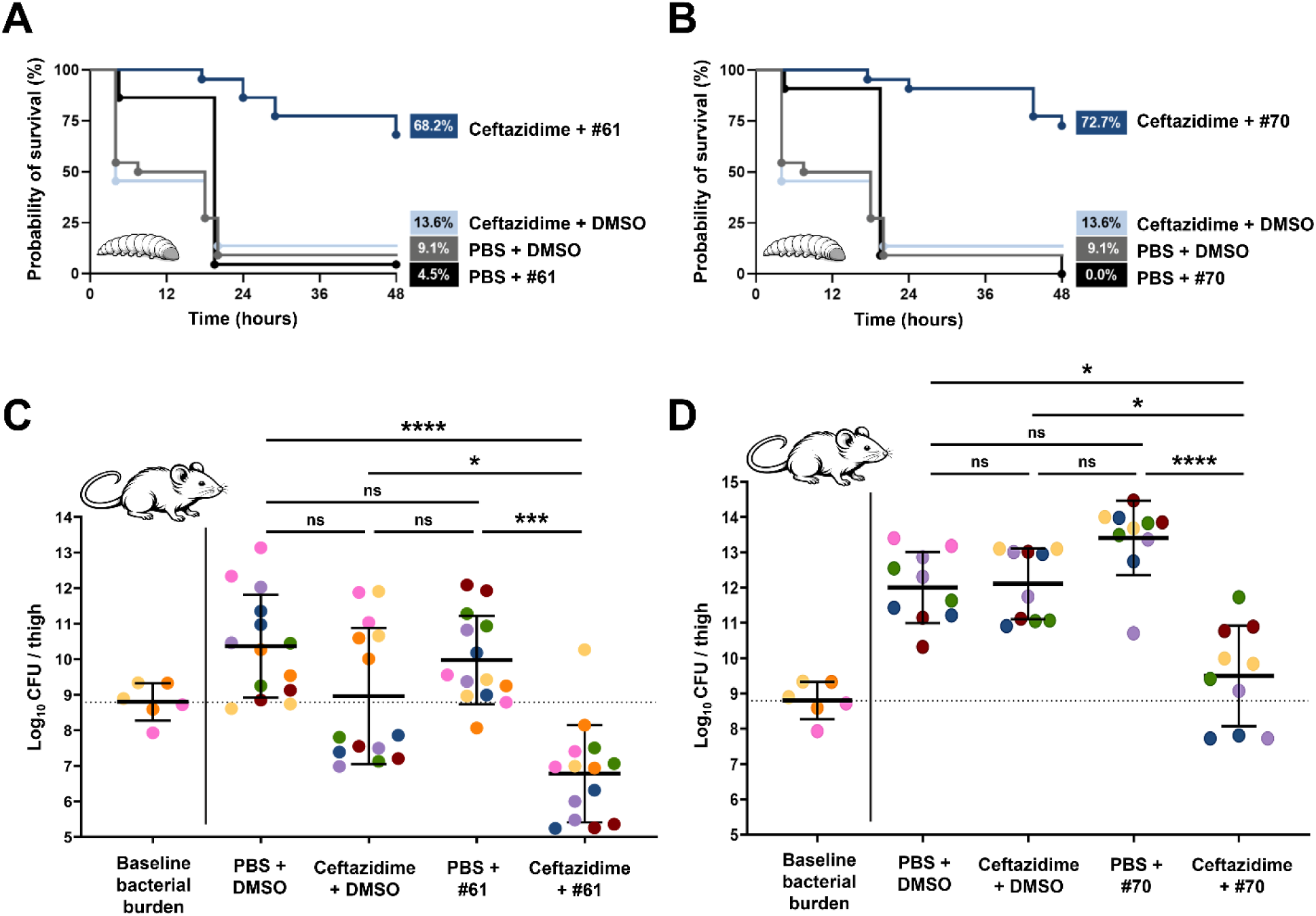
Inhibitor hits #61 and #70 rescue *G. mellonella* larvae from infection and reduce bacterial burden in neutropenic mice. (A, **B)** *G. mellonella* larvae infected with *K. pneumoniae* ZRI show low survival within 48 hours when untreated (9.1% survival, grey line). Treatment with inhibitor hit #61 or #70 (4.5% or 0.0% survival, respectively, black line) or the antibiotic ceftazidime (13.6% survival, light blue line) did not improve survival. In comparison, larvae treated with the antibiotic-inhibitor #61 or #70 combinations show 68.2% or 72.7% survival 48 hours post-infection, respectively (dark blue line), exhibiting a significant improvement in survival compared to all other treatment regimens. The graph shows Kaplan-Meier survival curves of infected *G. mellonella* larvae after different treatment applications; horizontal lines represent the percentage of larvae surviving after application of each treatment at the indicated time point (a total of 22 larvae were used for each condition). The same untreated and ceftazidime-treated groups of larvae were used in both panels (A) and (B). Statistical analysis of this data was performed in GraphPad Prism v10.6.1 using a Mantel-Cox test with the Holm-Šídák correction for multiple comparisons; n=22; p=0.2148 (non-significance) (*K. pneumoniae* ZRI + PBS + DMSO vs. *K. pneumoniae* ZRI + PBS + #61), p=0.2384 (non-significance) (*K. pneumoniae* ZRI + PBS + DMSO vs. *K. pneumoniae* ZRI + PBS + #70), p=0.7838 (non-significance) (*K. pneumoniae* ZRI + PBS + DMSO vs. *K. pneumoniae* ZRI + ceftazidime + DMSO), p=0.2148 (non-significance) (*K. pneumoniae* ZRI + ceftazidime + DMSO vs. *K. pneumoniae* ZRI + PBS + #61), p=0.2479 (non-significance) (*K. pneumoniae* ZRI + ceftazidime + DMSO vs. *K. pneumoniae* ZRI + PBS + #70), p<0.0001 (significance, ****) (*K. pneumoniae* ZRI + PBS + DMSO vs. *K. pneumoniae* ZRI + ceftazidime + #61), p<0.0001 (significance, ****) (*K. pneumoniae* ZRI + PBS + DMSO vs. *K. pneumoniae* ZRI + ceftazidime + #70), p<0.0001 (significance, ****) (*K. pneumoniae* ZRI + ceftazidime + #61 vs. *K. pneumoniae* ZRI + PBS + #61), p<0.0001 (significance, ****) (*K. pneumoniae* ZRI + ceftazidime + #70 vs. *K. pneumoniae* ZRI + PBS + #70), p<0.0001 (significance, ****) (*K. pneumoniae* ZRI + ceftazidime + DMSO vs. *K. pneumoniae* ZRI + ceftazidime + #61), p<0.0001 (significance, ****) (*K. pneumoniae* ZRI + ceftazidime + DMSO vs. *K. pneumoniae* ZRI + ceftazidime + #70). **(C)** Bacterial burden expressed as CFU/thigh in a neutropenic thigh infection model with *K. pneumoniae* ZRI. All groups were infected with *K. pneumoniae* ZRI at an inoculum of ∼2 x 10^5^ CFU/mL, which was equivalent to ∼1.01 x 10^9^ CFU/thigh two hours post-infection, when the first treatment was administered (baseline bacterial burden, left of the solid line, marked by a dotted line). Bacterial burden was assessed for groups of 7 mice (n=7 per group) that were treated with **(1)** the carrier control (PBS + DMSO), **(2)** antibiotic only (ceftazidime + DMSO), **(3)** inhibitor compound only (PBS + #61), or **(4)** combination treatment (ceftazidime + inhibitor hit #61). All treatment regimens, except the combination treatment, show an increase in bacterial burden. By contrast, the combination treatment, ceftazidime alongside the inhibitor hit #61, results in a three-log drop in CFU/thigh in comparison to the carrier control showing antibiotic potentiation. Statistical analysis was performed in GraphPad Prism v10.6.1 using a Kruskal-Wallis test; p>0.9999 (non-significance) (PBS + DMSO vs. #61), p=0.4294 (non-significance) (PBS + DMSO vs. ceftazidime + DMSO), p=0.9550 (non-significance) (#61 vs. ceftazidime + DMSO), p<0.0001 (significance, ****) (PBS + DMSO vs. ceftazidime + #61), p=0.0002 (significance, ***) (#61 vs. ceftazidime + #61), p=0.0314 (significance, *) (ceftazidime + DMSO vs. ceftazidime + #61). **(D)** Bacterial burden expressed as CFU/thigh in a neutropenic thigh infection model with *K. pneumoniae* ZRI. All groups were infected with *K. pneumoniae* ZRI at an inoculum of ∼2 x 10^5^ CFU/mL, which was equivalent to ∼1.01 x 10^9^ CFU/thigh two hours post-infection, when the first treatment was administered (baseline bacterial burden, left of the solid line, marked by a dotted line). Bacterial burden was assessed for groups of 5 mice (n=5 per group) that were treated with **(1)** the carrier control (PBS + DMSO), **(2)** antibiotic only (ceftazidime + DMSO), **(3)** inhibitor compound only (PBS + #70), or **(4)** combination treatment (ceftazidime + inhibitor hit #70). All treatment regimens, except the combination treatment, show an increase in bacterial burden. By contrast, the combination treatment, ceftazidime alongside the inhibitor hit #70, results in a three-log drop in CFU/thigh in comparison to the carrier control showing antibiotic potentiation. Statistical analysis was performed in GraphPad Prism v10.6.1 using a Kruskal-Wallis test; p=0.1930 (non-significance) (PBS + DMSO vs. #70), p>0.9999 (non-significance) (PBS + DMSO vs. ceftazidime + DMSO), p=0.2225 (non-significance) (#70 vs. ceftazidime + DMSO), p<0.0374 (significance, *) (PBS + DMSO vs. ceftazidime + #70), p<0.0001 (significance, ****) (#70 vs. ceftazidime + #70), p=0.0314 (significance, *) (ceftazidime + DMSO vs. ceftazidime + #70). For panels (C) and (D) the two thighs of each mouse are represented by the same color. Raw data for all panels are available in **Supplementary File S7**.

We first assessed efficacy in a *Galleria mellonella* infection model (**Figure 6A,B**). Untreated larvae exhibited high mortality within 48 hours following infection, and survival was not improved by treatment with ceftazidime alone, compound #61 alone, or compound #70 alone. In contrast, co-administration of ceftazidime with either inhibitor resulted in a marked survival benefit, increasing survival from 9.1% in the untreated group to 68.2% and 72.7% for compounds #61 and #70, respectively. Thus, neither the antibiotic nor the inhibitors were effective as monotherapies under these conditions, whereas their combination substantially improved host survival. Injection controls confirmed that the experimental procedure itself did not contribute to larval mortality (**Supplementary Figure S12D**).

We next evaluated compound efficacy in a neutropenic murine thigh infection model (**Figure 6C,D**). Consistent with the results obtained in *G. mellonella*, treatment with ceftazidime alone or inhibitor hits alone failed to reduce bacterial burden in the thigh muscle relative to the carrier control. In contrast, combination treatment with ceftazidime and either compound #61 or #70 resulted in approximately three-log reductions in bacterial burden. These reductions were significant relative to both untreated controls and antibiotic or inhibitor monotherapy, demonstrating robust *in vivo* potentiation of ceftazidime activity.

Together, our results show that the inhibitor hits identified by PubCheF-1 retain efficacy in insect and animal infection models and can restore the activity of an otherwise ineffective β-lactam antibiotic against a multidrug-resistant clinical isolate.

## DISCUSSION

To test whether we could leverage the language of scientific literature for inhibitor discovery at scale, we developed PubCheF-1, a deep learning model trained on chemical structures and their literature-derived functional labels. PubCheF-1 was used to explore the identification of structurally distinct inhibitors targeting β-lactamases, as they represent clinically important antibiotic resistance determinants that remain challenging therapeutic targets. Notably, most of the enzymes included in our screens, and especially the metallo-β-lactamase L1, currently lack effective inhibitors [24,27]. From a set of ∼13.2 million purchasable molecules, PubCheF-1 predicted ∼30,000 of them to be relevant for β-lactamase inhibition. These were down-selected for structural novelty, low toxicity, and low reactivity to 3,880 molecules, from which a candidate set of 110 compounds was experimentally tested. Protein target and in-bacteria screens indicated that 14 compounds indeed functioned as β-lactamase inhibitors, a remarkable hit rate of 12.7% overall (**Figure 3**) [46–48,57,58].

Several inhibitors, including all identified hits against KPC and L1 enzymes in bacteria, remained active across increasingly stringent stages of validation (**Figure 4A-D**) and the most active compounds showed direct target engagement (**Figure 4E-G**), restored β-lactam activity in clinically relevant isolates (**Figure 5**), and retained efficacy *in vivo* (**Figure 6**). In contrast to typical discovery campaigns, where identified compounds are often eliminated because of toxicity, assay interference, or poor development potential [25], both inhibitor hits #61 and #70 were suitable for *in vivo* evaluation and demonstrated potent activity in an animal infection model (**Figure 6**).

This proof-of-principle shows that the relationships between structure and function that are inherently present in decades of scientific literature can be leveraged by deep learning models to understand and predict biological function for inhibitor discovery. This reflects the fact that scientific papers are the primary means by which humans communicate scientific knowledge, and thus deep learning models trained on this same corpus should likewise be able to distill the underlying relationships. Since the semantic structure of scientific language is dense, well-connected, and coherent (**Figure 2C**), training across broad profiles of function ultimately allowed PubCheF-1 to converge upon real-world chemical structure-function relationships (**Figure 2D**, **Supplementary Figure S3F-G**) that would have been difficult to extract from isolated experimental measurements (**Supplementary Figure S3B**). For example, although candidate selection in this study relied solely on labels containing the substring “lactam”, we found that the model was able to distinguish metallo-β-lactamase hits from serine-based ones (**Supplementary Figure S9**). Similarly, PubCheF-1 was able to distinguish the modes of action for the otherwise similar boronate hits. We found that compound #61 functions as a prodrug whereas #70 exposes the active pharmacophore in the KPC-3 active site without requiring prior breakdown (**Figure 4E-G** and **Supplementary Figure S8**) and, accordingly, PubCheF-1 moderately predicted the label “prodrug” for #61 but not for #70. In the future, predicted label profiles may greatly enable more targeted virtual screening by simultaneously prioritizing molecules with favorable functional attributes while deprioritizing molecules with less desirable ones. Furthermore, these labels may be useful not only for hit identification, but also for serving as experimental hypotheses to elucidate mechanisms and potential off-target effects. As such, we have made the PubCheF-1 predicted labels over the ZINC20 purchasable set of 13.2 million molecules freely available and accessible online (https://www.pubchef.org).

It is especially interesting that inhibitors #61 and #70, as well as four more of our hits (**Figure 3C**), were boronate compounds. The boronate moiety has previously proved to be applicable to β-lactamase inhibition, and two such molecules have been successful in advancing into clinical application [59,60]. The PubCheF dataset contained roughly 4,700 compounds containing boron, with roughly 10% of these having been studied for β-lactamase inhibition, all of which have <0.50 RDKit Tanimoto similarity to #61 and #70. More specifically, while the formylphenylboronate breakdown product of #61 does have known β-lactamase inhibitor activity [61] and was present in the PubCheF-1 training data, the protecting group of compound #61 is entirely novel in the realm of β-lactamase inhibition, having only been tested for non-lactamase antibiotic activity [62] and Alzheimer’s treatment [63]. And while there were studied furanylboronate compounds in the PubCheF-1 training data, there were no instances of the formylfuranylboronate breakdown product of compound #70. Moreover, furylboronate alone was studied for β-lactamase inhibition [64] and was found to be inactive. Together, this analysis suggests that PubCheF-1 is able to integrate disparate scientific observations and successfully predict inhibitor activity for chemical structures whose inhibitory potential, while plausibly discoverable by conventional medicinal chemistry, has nonetheless remained undiscovered.

There are of course limitations to this approach. Although PubCheF-1 frequently converged upon medicinal chemistry features consistent with established mechanisms of action, this behavior was not universal across molecules, highlighting the need for improved alignment between model predictions and mechanistic understanding. The functional labels used in the PubCheF dataset would also benefit from a more formal ontological organization to better capture relationships between related functions and to contextualize broad functional categories and labels. Since the development of PubCheF-1 predates several recent advances in deep learning, the performance reported here likely represents a lower bound in the ability to predict function, and additional gains may be realized by integrating functional data from other textual sources of chemical effects such as patient reports, sensory profiles, and chemical patents, with the latter having over 30 million linked molecules in contrast to the <2.2 million linked to scientific articles that were used in this study.

Limitations aside, the high hit rate, low hit attrition and potent activity of the validated inhibitors presented here suggest that the PubCheF-1 model and down-selection framework favor compounds capable of efficient progression through successive stages of drug discovery. Traditionally, β-lactamase inhibitor discovery campaigns rely on high-throughput screening of tens of thousands of compounds. These efforts are often followed by extensive downstream validation and substantial and costly attrition of candidate hits, ultimately resulting in low confirmed hit rates of <5% [47,48,57,58]. We were able to increase that rate, with an investment of only $4,000 in compounds. Given that this approach is orthogonal to structure-based methods, combining the two should greatly accelerate the identification of bioactive molecules across diverse therapeutic areas and across increasingly complex molecular scales, ranging from quantum to clinical properties.

## ACKNOWLEDGEMENTS

This work was supported by the National Institutes of Health under grants R01AI158753 (to D.A.I.M.), R35GM122480 (to E.M.M.), and R35GM148356 (to Y.J.Z.). Additional support was provided by the Welch Foundation through grants F-2250 (to D.A.I.M.), F-1654 (to A.D.E.), F-1515 (to E.M.M.), and F-0018 (to J.L.S.). E.V.A. acknowledges support from the Welch Regent Chair of Chemistry (F-0046) and D.A.I.M acknowledges additional support from the Texas Biologics (TXBio) grant TXB-24-02 from the Cockrell School of Engineering at The University of Texas at Austin and the UT | Portugal Extra Exploratory Project grant 2022.15740.UTA from the Fundação para a Ciência e a Tecnologia, I.P. Computational analyses were performed using the Biomedical Research Computing Facility at UT Austin, Center for Biomedical Research Support. RRID#: SCR_021979. We thank Kevin K. Yang (Microsoft Research) and Sophie Helaine (Harvard Medical School) for their invaluable feedback on the manuscript.

## AUTHOR CONTRIBUTIONS

C.W.K., N.K., K.K., A.J.S.M., E.M.M., Y.J.Z., A.D.E., and D.A.I.M. designed the research. C.W.K. performed the majority of the computational work described in this study with assistance from P.W. and K.X. C.W.K. and D.W. developed the PubCheF-1 Explorer website. N.K. performed protein production, protein target and in-bacteria screening, hit validation, and clinical strain panel experiments. K.K. performed protein production, inhibitor hit binding experiments, and determined crystal structures. A.J.S.M. assisted with in-bacteria screening and performed insect and mouse infection studies. A.D. performed *S. maltophilia* genetics and assisted with mouse infection studies. T.B. and J.L.S. performed NMR analyses. F.K. assisted with protein production. E.V.A. provided expertise in boron chemistry. C.W.K., N.K., K.K., A.J.S.M., E.M.M., Y.J.Z., A.D.E., and D.A.I.M wrote the manuscript with input from all authors. E.M.M., Y.J.Z., A.D.E., and D.A.I.M. supervised the project.

## DECLARATION OF INTERESTS

C.W.K., N.K., K.K., A.J.S.M., E.M.M., Y.J.Z., A.D.E., and D.A.I.M are inventors on a patent application covering compounds described in this study, assigned to The University of Texas at Austin. All other authors declare no competing interests.

## MATERIALS AND METHODS

### COMPUTATIONAL METHODS

#### Text extraction, large language model fine-tuning, and validation

For each PubMed article, the title and abstract were used as input to a large language model (LLM) to generate functional labels for the described molecule(s). The system prompt was “*You are an organic chemist summarizing scientific literature*” and the user prompt was “*Return a set of a few (1-5) 1-3 word descriptors that best describe the chemical or pharmacological function(s) of the molecule described by the given article. Be concise, as specific as possible, but not necessarily comprehensive (choose a small number of great descriptors). For example, prefer ‘Dihydrofolate Reductase Inhibitor’ over ‘Enzyme Inhibitor’. Follow the syntax ‘{descriptor_1} / {descriptor_2} / {etc}’, writing ‘NA’ if nothing is provided. DO NOT BREAK THIS SYNTAX. The following is the article info:*” followed by the title and abstract or the article. Three OpenAI LLMs, gpt-3.5-turbo-0613, gpt-4-turbo, and gpt-3.5-turbo-0613 fine-tuned on a set of 100 curated non-test examples, were evaluated on chemical function extraction over a test set of 100 articles, the results of which were validated manually to obtain precision, recall, and vocabulary size.

#### PubCheF dataset creation

The PubChem CID - PubMed PMID dataset (accessed 1/11/24) was filtered to include only PMIDs in the BioAssay category. For molecules with over 1,000 linked PMIDs, only 1,000 randomly selected articles were used. The fine-tuned gpt-3.5-turbo-0613 model was chosen to generate labels for all remaining PubMed articles, producing 24,328 labels. OpenAI’s text-embedding-3-large model was used to obtain text embeddings of all labels. These were DBSCAN [65] clustered to merge semantically similar terms to further consolidate the vocabulary. To evaluate the optimal clustering parameters, a test set of 1,618 labels grouped into 92 clusters of common biochemical function was created (**Supplementary File S1**). The DBSCAN parameters that maximized F1 score while maintaining >0.90 precision were chosen (**Supplementary Table S2**). Each label was then mapped to the label corresponding to its cluster’s centroid. Labels were then additionally merged based on shared alphanumeric string, those indicating no function were excluded, and those from PubMed articles that linked to more than 100 molecules were also excluded to reduce noise introduced from high-throughput screens (**Supplementary Figure S1A**). Only the 30 most common labels for each molecule were retained to reduce noise introduced from extremely well-studied molecules. Labels were then propagated between molecules with the exact same RDKit [34] molecular fingerprint (1.0 Tanimoto similarity) to mitigate differences introduced by salts or charge. Although this removed chirality, it had minimal impact on model performance (**Supplementary Figure S3D**). The final dataset was validated for 186 distinct molecule-label pairs by manually examining the activity data reported in each pair’s associated manuscript.

#### Co-occurrence graph labeling

Cosmograph (cosmograph.app) was used to plot the connectivity of labels in the PubCheF dataset by molecular co-occurrence (two labels occurring in the same molecule), ignoring edge weights. The following Cosmograph simulation values were used: 0.4 gravity, 2 repulsion, 2 repulsion theta, 0.03 link strength, 20 minimum link distance, 0.85 friction, and high resolution.

#### PubCheF-1 model, ablations, and training

The PubCheF dataset was preprocessed to remove molecules that tokenized with the ChemBERTa-77M-MLM tokenizer (an element-wise tokenizer) to over 509 tokens. Labels occurring in fewer than 50 molecules were also excluded to remove infrequent examples; this cutoff was arbitrarily chosen (**Supplementary Figure S1B**). The dataset was partitioned into training, validation, and test sets (80/10/10) using a scaffold split. Two models were trained on this dataset: **(1)** a logistic regression model with RDKit fingerprints as input (C=0.001, max_iter=1000, OneVsRestClassifier), and **(2)** the chemical language model ChemBERTa-77M-MLM. ChemBERTa-77M-MLM is a RoBERTa-style model [66] trained on 77 million chiral RDKit-canonicalized PubChem molecules using a masked language modeling objective. This model has three layers and a hidden dimension of 384. At the time of use, the ChemBERTa-77M-MLM tokenizer hosted on HuggingFace was not functioning as intended, as it tokenized molecules at the character level rather than the element level (e.g., Cl was tokenized as [C, UNK] rather than [Cl]). This behavior was inconsistent with the reported vocabulary and training procedure. A custom tokenizer was therefore implemented to reproduce the intended tokenization scheme. Then, a hyperparameter search was conducted to optimize validation loss after 20 epochs, varying model hyperparameters as well as loss function (cross entropy vs. focal loss), and the use of model ensembling. The best-performing configuration, dubbed PubCheF-1, was a three-model ensemble trained with focal loss using the hyperparameters listed in **Supplementary Table S3**. For the ten models trained on a single random label shown in **Supplementary Figure S3D**, a non-ensembled PubCheF-1 (random seed 42) for binary classification was used with the same hyperparameters.

#### ChEMBL dataset creation and model training

To ablate the PubCheF dataset, an equivalent model was trained on a ChEMBL-derived target prediction dataset. We used ChEMBL 33 (June 2023), released contemporaneously with the PubCheF dataset (Jan. 2024), so that the training datasets were temporally similar. Following the criteria used for the Polypharmacology Browser PPB3 [67], a compound was labeled as active against a target if it had an IC50, EC50, GI50, Ki, Kd, potency value ≤10µM or percent inhibition >50%. Labels were positive only to provide a multi-label prediction task. We retained molecules of ≤80 non-hydrogen atoms and targets with ≥5 active molecules over all target types (single proteins, protein complexes, protein families, cell lines, organisms, and others). SMILES were canonicalized with RDKit and molecules exceeding 510 tokens were removed. The final dataset comprised 1,329,817 molecules and 6,588 targets. Molecules were then partitioned into training, validation, and test sets (80/10/10) using Bemis-Murcko scaffolds. To prevent data leakage in the PubCheF-test-reduced benchmark, scaffolds were assigned to their respective folds in the PubCheF training set and then greedily filled until the expected ratios were attained. We used identical architecture and hyperparameters as PubCheF-1, changing only the training data. This model is referred to as “PubCheF-1 (ChEMBL)”.

#### PubCheF-test-reduced and OpenTargets-indications benchmarks

To reduce bias and redundancy for benchmarking, the PubCheF test set was filtered from 120,131 to 10,311 molecules by ensuring that no molecules had greater than 0.50 RDKit fingerprint Tanimoto similarity to any other molecule in the set [68]. The OpenTargets-indications set contained 6,677 molecules. While some overlap between the models’ training data and these datasets cannot be excluded, the prediction tasks differed substantially, with the PubCheF benchmark being based on literature-derived functional annotations and OpenTargets-indications focused on therapeutic indications. To enable benchmarking against generative models such as MolT5, classifier models (CheF, PubCheF-1, and PubCheF-1 (ChEMBL)) were converted into discrete term predictions using a score threshold calculated by applying a distributed score cutoff selected to maximize label-specific F1 score on the PubCheF validation set and CheF test set, respectively [69]. MolT5 was trained on the OEChem canonicalization scheme present on PubChem, rather than the RDKit scheme used for CheF and PubCheF-1, and so inputs were converted accordingly to obtain predictions. The judge LLM, gpt-4-turbo-2024-04-09 was used to create the confusion matrix for each model’s predictions across both benchmarks.

For CheF, PubCheF-1, and PubCheF-1 (ChEMBL), the evaluation system prompt was “*You are a skilled medicinal chemist evaluating a model’s predictions of a molecule against the ground-truth*” and the user prompt was “*The following are predicted attributes of about the said molecule: ‘{ predictions }’, The following are the true attributes of this molecule: { actual } Ignore all claims about the molecule’s structure and only focus on its functional attributes. Compute the TP/FP/FN of the predicted attributes given the true attributes. Finish your response with the exact form: ‘TP:a,FP:b,FN:c’*”. For MolT5, the system prompt was “*You are a skilled medicinal chemist evaluating a model’s predictions of a molecule against the ground-truth*” and the user prompt was “*Break this prediction down into a list of the individual claims about the said molecule: ‘{ predictions }’, The following are the true attributes of this molecule: ‘{ actual } Ignore all claims about the molecule’s structure and only focus on its functional attributes. Compute the TP/FP/FN of the predicted claims given the true attributes. Finish your response with the exact form: ‘TP:a,FP:b,FN:c’*”. The confusion matrix was parsed from the generated text, from which we computed F1, hit-rate, and macro precision. Hit-rate was the fraction of molecules with at least one true positive. Macro precision was the mean across molecules of TP/(TP+FP), computed over each model’s predicted set; molecules with no predictions were assigned a precision of 0. Molecules whose confusion matrix could not be parsed were excluded, leaving 10,283 molecules for PubCheF-test-reduced and 6,661 for OpenTargets-indications. For 300 examples comprising 50 predictions from three models (CheF, PubCheF, and MolT5) across both the PubCheF-test-reduced and OpenTargets-indications benchmarks, the metrics computed by the judge LLM were compared to those obtained by author K.X. Both benchmark datasets are provided in **Supplementary File S1**.

#### Input ablation study

A non-ensembled PubCheF-1 model (random seed 42) was re-trained on the following variants of the PubCheF dataset: SMILES with random RDKit canonicalization order for each occurrence, achiral SMILES, PubCheF without label propagation, PubCheF without label propagation with achiral SMILES, PubCheF in which the SMILES strings were shuffled at the token level, and PubCheF in which the input was ignored and replaced with a random vector before the classification layer each forward pass. These were compared by their validation set macro averaged PR-AUC.

#### Token composition vs. fingerprint Tanimoto distance Mantel tests

Pairwise molecular similarity was first computed from RDKit fingerprints using the Tanimoto coefficient and converted to a distance matrix as one minus similarity. SMILES strings were then tokenized (with [cls], [sep], [pad], and [mask] tokens excluded) and each molecule was represented as a token-count vector (token multiplicity retained, token order ignored), and pairwise similarity was calculated using generalized Tanimoto similarity on the count vectors; distances were again defined as one minus similarity. A Mantel test (Pearson; 9,999 permutations; scikit-bio) was performed to quantify the correspondence between the fingerprint distance matrix and the token composition distance matrix across the 10,311 molecules in the PubCheF-test-reduced dataset.

#### Model weight vs. co-occurrence correlations

Molecular co-occurrence counts were converted to a pseudo-distance metric by normalizing the joint co-occurrence count by the sum of the total counts of both elements and subtracting the resulting value from 1:

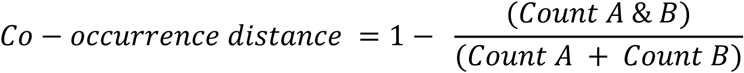

A Mantel test (skbio; 9,999 permutations) was used to compute the Pearson correlation between the model weights of the classification layer of one of the PubCheF-1 models (seed 42; exalted-sweep-1) and the label co-occurrence distance. This was computed for each of the following top-K label matrices: 50, 100, 200, 500, 1000, 2000, 3000, and 4000.

#### Layer-wise relevance propagation

The method from Chefer et al. was adapted for PubCheF-1 [40]. For a given label and molecule, importance values were averaged across each model in the PubCheF-1 ensemble. Values corresponding to elements were kept and then min-max normalized.

In the analysis to systematically understand where PubCheF-1 places importance, LRP values were computed for each molecule in the PubCheF test set using one randomly selected ground-truth label. For each molecule, the ‘important’ substructures were defined as the atoms above the knee point of the cumulative sum of the atom-wise importance values sorted in ascending order. These atoms were converted into connected substructures using RDKit, from which we determined the average substructure size, the number of substructures, and their heteroatom content (excluding H and C). These metrics were then compared with those obtained from randomly selected atom sets of equal size to assess whether PubCheF-1 preferentially concentrated importance on a smaller number of larger, heteroatom-enriched substructures.

#### PubCheF-1 *in silico* hit discovery analysis

PubCheF-1 predictions over the purchasable set of ZINC20 compounds (2024-04-30) were used to select molecules for five targets: KPC-3 β-lactamase (**Supplementary Table S7**), 5-HT2A receptor (PDB: 6A93), SARS-CoV-2 main protease (Mpro; PDB: 7CAM), Taq DNA polymerase (PDB: 1TAQ), and Hepatitis C Virus (HCV) NS5B RNA polymerase (PDB: 3FQK). A molecule was considered a predicted hit if it was high-scoring for any of the labels used for the given target. The labels used for each target spanned different selection criteria and were as follows:

#### KPC-3 β-lactamase

Any label containing the substring “lactam” (case-insensitive), i.e., “Beta-Lactamase Inhibitor”, “Metallo-Beta-Lactamase Inhibitor”, “Beta-lactamase Inhibitor”, “AmpC Beta-lactamase Inhibitor”, “Beta-Lactam Adjuvant”, “Beta-Lactamase”. This set of labels corresponded to 2,009 molecules in the PubCheF dataset.

#### 5-HT2A Receptor

“5-HT2A Receptor Ligand”. This label corresponded to 1,209 molecules in the PubCheF dataset.

#### SARS-CoV-2 Mpro

“3CL Protease Inhibitor”, “SARS-CoV-2 3CL Protease Inhibitor”, “SARS-CoV Mpro Inhibitor”, “Main Protease Inhibitor”, “3CL(pro) Inhibitor”, “Mpro Inhibitor”, “SARS-CoV-2 Main Proteinase Inhibitor”. This label set corresponded to 2,306 molecules in the PubCheF dataset.

#### Taq DNA Polymerase

“DNA Polymerase Inhibitor”. This label corresponded to 744 molecules in the PubCheF dataset.

#### HCV NS5B RNA polymerase

“HCV NS5b Polymerase Inhibitor”, “NS5B RdRp Inhibitor”, “HCV RdRp Inhibitor”. This label set corresponded to 7,765 molecules in the PubCheF dataset.

The hits were partitioned into two sets: top hits (unfiltered) and novel hits (those with RDKit fingerprint Tanimoto coefficient < 0.50 to anything in the original PubCheF dataset with the target labels). The hits were then filtered to remove molecules containing a run of ≥8 acyclic carbons, as long-chain fatty acids represent a known source of false-positive predictions owing to their frequent annotation in scientific literature. For each of the five targets, the 100 highest-scoring molecules passing these criteria were selected from each prediction set (1,000 molecules in total), together with 100 molecules randomly sampled from the ZINC20 purchasable set as a control.

Using Boltz-2 (v2.2.1, seed 42), all ligands were co-folded with each of the five targets (6,000 total co-folds) and affinity probabilities were obtained. RDKit failed to generate a 3D conformer for two molecules in the novel set (one KPC-3 hit and one control; 2,990/3,000 co-folds were completed) and twelve in the top-ranked set (four 5-HT2A hits, seven NS5B hits, one control; 2,940/3,000 co-folds were completed). To assess whether PubCheF-1 predictions were target-specific, we compared the average Boltz-2 affinity probabilities for compounds against their predicted targets with those obtained for all cross-target combinations and for randomly selected ZINC20 control molecules.

#### β-Lactamase inhibitor prediction and down-selection

Molecules from the purchasable set of ZINC20 were classified as possible β-lactamase inhibitors if they were predicted with ≥0.20 PubCheF-1 score for labels that contained the substring “lactam”, including the labels “Beta-Lactamase Inhibitor”, “Metallo-Beta-Lactamase Inhibitor”, “AmpC Beta-Lactamase Inhibitor”, “Beta-Lactam Adjuvant”, and “Beta-Lactamase”, producing 30,267 molecules. These were then filtered for novelty (≤0.50 RDKit fingerprint Tanimoto similarity to a molecule in the PubCheF dataset whose labels contained the substring “lactam”), low toxicity (≤0.20 probability of cell toxicity from the HepG2, HSkMC, and IMR-90 models used in [42]), absence of Pan-Assay Interference Substructures (PAINS), and availability at MolPort or AKSci vendors. Brenk substructures were not filtered out as this would have removed desirable mechanisms such as metal binding. From the final set of 3,880 molecules, we chose a set of molecules semi-arbitrarily, optimizing cost (maximum expenditure of approximately $4,000), structural diversity, high PubCheF-1 score, and minimal Brenk flags.

#### PubCheF-1 comparison to Boltz-2

To create a lightweight evaluation set, KPC hits #54, #59, #61, #62, #70, #91, and #94, L1_#3637_ hits #35, #40, #42, and #108, and the remaining 100 non-active PubCheF-1 predictions were added to a set of presumed negatives created from 1,084 randomly selected molecules in ZINC20 with less than 0.5 RDKit fingerprint Tanimoto similarity to any molecule in the PubCheF dataset containing the substring “lactam”. To determine scores for PubCheF-1, the highest score of any label containing “lactam” was used. For Boltz-2, protein-ligand complexes were predicted for both KPC-3 and L1_#3637_ (**Supplementary Table S7**) and the maximum average affinity score between both enzyme structures was used. Compounds 38 and 77 were excluded from the evaluation set as they failed to co-fold in Boltz-2.

#### PubCheF-1 Explorer website

Predictions across the ZINC20 purchasable dataset for all labels in the PubCheF-1 vocabulary are available at the PubCheF website, https://www.pubchef.org. This website contains only predictions above a 0.10 predicted score from the PubCheF-1 model. Users can filter and sort by specific functional labels, apply structural similarity filters, download top-ranking compounds, and visit PubChem pages for the contained molecules. This dataset is also available for download at https://doi.org/10.5281/zenodo.21108754. This website has been made available as an open resource for the community to facilitate the discovery of potentially useful functions for readily purchasable molecules.

### EXPERIMENTAL METHODS

#### Reagents and bacterial growth conditions

Unless otherwise stated, chemicals and reagents were acquired from Sigma Aldrich, Fisher Scientific and VWR, growth media were purchased from Fisher Scientific, VWR, and Oxoid, and antibiotics were obtained from Melford Laboratories and Fisher Scientific. Tested candidate compounds, identified through the PubCheF-1 model, were acquired from Aaron Chemicals, AK Scientific, AstaTech, ChemBridge, ChemDiv, CombiBlocks, and Otava; predicted compound #90 arrived damaged and was not tested.

Lysogeny broth (LB) (10 g/L NaCl) and agar (1.5% w/v) were used for routine growth of all organisms at 37 °C with shaking at 220 RPM, as appropriate. Mueller-Hinton (MH) broth and agar (1.5% w/v) were used for Minimum Inhibitory Concentration (MIC) assays and for compound screening assays. Growth media were supplemented with the following, as required: 0.25 mM Isopropyl β-D-1-thiogalactopyranoside (IPTG), 12.5 μg/mL tetracycline, 50 μg/mL kanamycin, 100 μg/mL ampicillin, 50 μg/mL streptomycin (for cloning purposes), and 4000 μg/mL streptomycin (for genetic manipulation of *S. maltophilia* clinical isolates).

#### Construction of plasmids and bacterial strains

Bacterial strains, plasmids and oligonucleotides used in this study are listed in **Supplementary Tables S8**, **S9**, and **S10** respectively. DNA manipulation was conducted using standard methods. KOD Hot Start DNA polymerase (Merck) or Phusion High-Fidelity DNA Polymerase (Thermo Fisher Scientific) were used for all PCR reactions according to the manufacturer’s instructions, oligonucleotides were synthesized by Sigma Aldrich, and restriction enzymes were purchased from New England Biolabs. DNA g-blocks for *bla*_EC-2_, *bla*_L1 #4488_, and *bla*_L1 #4509_ were purchased from Twist Biosciences. All plasmids and mutant strains were DNA sequenced (Sanger sequencing, Eurofins; whole-plasmid or whole-genome sequencing, Plasmidsaurus) and confirmed to be correct before use. More specifically:

#### pDM1-based plasmids

DNA fragments encoding L1_#4488_, and L1_#4509_ (**Supplementary Table S6, Supplementary Table S7**) were amplified (P1-P4) and cloned into the pDM1 vector using SacI/KpnI.

#### pet28-based plasmids

DNA fragments encoding KPC-3 and EC-2 (**Supplementary Table S7**) were amplified (P5-P8), and the pET28-His_8_-SUMO-3C vector backbone was linearized by inverse PCR (P9-P11). The resulting fragments were assembled using an in-house Gibson assembly mix.

#### Genomic mutants

The *S. maltophilia* AMM *bla*_L2_ mutant was constructed by allelic exchange, as previously described [70]. Briefly, DNA fragments upstream and downstream of the *bla*_L2_ genes were amplified using *S. maltophilia* AMM genomic DNA (primers P12/P13 (upstream) and P14/P15 (downstream)). Following overlap PCR (P12/P15), the mutator fragment was inserted into the BamHI/SpeI sites of the suicide pKNG101 vector, resulting in plasmid pKNG101-*bla*_L2_. Since pKNG101 [64] is not replicative in *S. maltophilia*, the vector is maintained in *E. coli* CC118λpir and mobilized into *S. maltophilia* by triparental conjugation. Transconjugants were selected on MH agar supplemented with streptomycin (4000 μg/mL) and ampicillin (100 μg/mL; the L1 β-lactamase of *S. maltophilia* AMM confers ampicillin resistance even in the absence of *bla*_L2_). Successful transconjugants were confirmed using PCR and mutants were resolved by exposure to 20% w/v sucrose. Colonies carrying a *bla*_L2_ gene deletion were identified via colony PCR (primers P16/P17) and the *S. maltophilia* AMM *bla*_L2_ mutant strain was confirmed by whole-genome sequencing.

#### Minimum inhibitory concentration (MIC) assays

Antibiotic MIC assays were carried out in accordance with the EUCAST recommendations using ETEST strips (BioMérieux) or broth microdilution (BMD), as described in [51]. Unless stated otherwise, all candidate compounds were tested at a final concentration of 32μg/mL. Carrier controls were included at equivalent volumes.

#### ETEST MIC assays

Briefly, overnight cultures of each strain to be tested were standardized to Optical density at 600 nm (OD_600_) 0.063 in 0.85% NaCl (equivalent to McFarland standard 0.5) and distributed evenly across the surface of MH agar plates. ETEST strips were placed on the surface of the plates, evenly spaced, and the plates were incubated for 18-24 hours at 37 °C. MICs were recorded according to the manufacturer’s instructions. All ETEST MIC values were determined in biological triplicate.

#### BMD MIC assays

Briefly, a series of antibiotic concentrations was prepared by two-fold serial dilution in MH broth in a clear-bottomed 96-well microtiter plate (Corning). The strain to be tested was added to the wells at approximately 5 x 10^4^ colony forming units (CFU) per well and plates were incubated for 18-24 hours at 37 °C. The MIC value was defined as the lowest antibiotic concentration with no visible bacterial growth in the wells. All BMD MIC values were determined in biological and technical triplicate.

#### S. maltophilia MIC assays

For *S. maltophilia* AMM *bla*_L2_ MIC assays, the following variations were implemented: MIC assays were performed using ETEST strips in synthetic cystic fibrosis sputum medium (SCFM) as described in [71]. Overnight cultures were standardized to OD_600_ 0.2 in SCFM prior to being distributed evenly across the plate surface. A final concentration of 64 μg/mL of candidate compounds was used. Due to the pale color of *S. maltophilia* growth on SCFM, MICs were recorded at 24 hours and checked again at 48 hours to ensure accuracy.

#### Expression and purification of proteins for the β-lactam hydrolysis screen

The genes encoding KPC-3 and EC-2 without their signal sequences (Δ1–22 for KPC-3 and Δ1–19 for EC-2, **Supplementary Table S7**) were expressed in the cytoplasm of BL21 SHuffle T7 Express or BL21 (DE3), respectively, from a pET28a(+)-derived vector as fusions with an N-terminal His_8_-SUMO tag followed by an HRV 3C protease cleavage site. Notably, when KPC-3 is natively expressed in the bacterial periplasm, it matures through the formation of a disulfide bond that is essential for protein folding and function [54]. To ensure that recombinant, leaderless KPC-3 was correctly folded, we used *E. coli* SHuffle, an engineered strain that promotes cytoplasmic disulfide bond formation [72]. In addition, following affinity purification, the protein was further exposed to a cysteine/cystine redox system in which L-cysteine and L-cystine were mixed at 1:1 in molar ratio in the dialysis buffer as described in [69]. Protein expression was carried out as follows: A 10 mL LB culture supplemented with kanamycin was grown overnight at 37 °C with shaking at 220 RPM and used to inoculate 1 L of Terrific Broth (TB) supplemented with kanamycin. Protein expression was induced with 0.5 mM IPTG at an OD_600_ of 0.7-0.9, and the culture was incubated at 18 °C and 200 rpm for 16 hours. Cells were harvested by centrifugation (4,000 × g, 20 min, 4 °C) and resuspended in lysis buffer (50 mM Tris-HCl pH 8.0, 500 mM NaCl, 20 mM imidazole, 10% glycerol). Cells were lysed by sonication on ice, and the lysate was clarified by centrifugation (27,000 × g, 45 min, 4 °C). The supernatant was applied to a HisPur™ Ni-NTA resin (ThermoFisher Scientific) equilibrated in lysis buffer, washed with the same buffer, and eluted with 50 mM Tris-HCl pH 8.0, 500 mM NaCl, 250 mM imidazole, 10% glycerol. Elution fractions containing KPC-3 or EC-2 were pooled and dialyzed overnight at 4 °C against dialysis buffer (20 mM Tris-HCl pH 8.0, 150 mM NaCl, 0.1 mM L-cysteine, 0.1 mM L-cystine for KPC-3; 20 mM Tris-HCl pH 8.0, 150 mM NaCl for EC-2). For KPC-3 the HRV 3C protease was included in the dialysis buffer to cleave the His_8_-SUMO tag, whereas for EC-2 the HRV 3C protease addition was omitted due to the tendency of EC-2 to aggregate without the solubility tag.

The genes encoding L1_#3637_ and OXA-10 native sequences followed by a C-terminal StrepII tag were expressed in the periplasm from the pDM1 vector [54] using *E. coli* MC1000 cells as follows: Two 5 mL LB cultures supplemented with tetracycline were grown overnight at 37 °C with shaking at 220 RPM, combined and used to inoculate 3.3 L of TB supplemented with tetracycline. Protein expression was induced with 0.5 mM IPTG at an OD_600_ of 0.7-0.9, and the culture was incubated at 37 °C and 205 rpm for 16 hours. Cells were harvested by centrifugation (6,000 × g, 15 min, 4°C) and resuspended in equilibration buffer (50 mM Tris-HCl pH 7.5 (for L1_#3637_) or pH 8.0 (for OXA-10), 150 mM NaCl) containing protease inhibitors (Roche). Cells were harvested, sphaeroplasted as described [73], except that EDTA was omitted, and the periplasmic fraction was applied to Strep-Tactin Sepharose (Iba Lifesciences) in equilibration buffer. The resin was washed with 50 mM Tris-HCl pH 7.5 (for L1_#3637_) or pH 8.0 (for OXA-10), 1 M NaCl and the protein was eluted with 50 mM Tris-HCl pH 7.5 (for L1_#3637_) or pH 8.0 (for OXA-10), 150 mM NaCl, 2.5 mM D-desthiobiotin. Elution fractions containing L1_#3637_ or OXA-10 were pooled and concentrated using an Amicon Ultra Centrifugal Filter with 10 kDa molecular weight cutoff (Millipore Sigma) according to the manufacturer’s instructions.

Protein purity for all four β-lactamase samples was assessed by SDS-PAGE analysis. 10% BisTris NuPAGE gels (ThermoFisher Scientific) and MES/SDS running buffer prepared according to the manufacturer’s instructions were used. Pre-stained protein markers (SeeBlue Plus 2, ThermoFisher Scientific) were included and gels were stained for total protein with SimplyBlue SafeStain (ThermoFisher Scientific) according to the manufacturer’s instructions. Protein concentration was determined using the Pierce BCA Protein Assay Kit (Thermo Fisher Scientific) according to the manufacturer’s instructions.

#### β-Lactam hydrolysis screen (protein target screen)

β-Lactamase activity and inhibition were evaluated using the chromogenic β-lactam substrate nitrocefin (Fisher). Purified β-lactamase enzymes (**Supplementary Table S7**), KPC-3 (0.050 μg/well), L1_#3637_-StrepII (0.088 μg/well), His_8_-SUMO-3C-EC-2 (0.233 μg/well), and OXA-10-StrepII (0.064 μg/well), were incubated with 32 μg/mL of each of the 110 candidate compounds and 200 μM nitrocefin in 100 μL reaction volumes prepared in 100 mM sodium phosphate buffer (pH 7.0). Nitrocefin hydrolysis was monitored at 25 °C by measuring absorbance at 490 nm every 60 seconds for 10 minutes using a BioTek Synergy H1 microplate reader (Gen5 v3.10 software). The amount of nitrocefin hydrolyzed over the 10-minute interval was quantified using a standard curve generated from acid-hydrolyzed nitrocefin standards. Inhibitory activity of each candidate compound was expressed as the percentage of nitrocefin hydrolyzed relative to the corresponding carrier-treated enzyme control. The raw and analyzed data and statistical analyses can be found in **Supplementary File S4**.

#### Bacterial survival screen (in-cell screen)

Bacterial survival in the presence of the 110 candidate compounds in the presence and absence of the β-lactam antibiotic ceftazidime was evaluated using *E. coli* MC1000 strains expressing either the KPC-3 or L1_#3637_ β-lactamase or carrying the empty vector pDM1. MH medium supplemented with 0.25 mM IPTG was prepared with antibiotics, candidate compounds, or carrier controls to generate the following conditions: **(1)** candidate compound (32 μg/mL), **(2)** candidate compound (32 μg/mL) + ceftazidime (0.1 μg/mL for MC1000 pDM1; 48 μg/mL for MC1000 pDM1-*bla*_KPC-3_ and MC1000 pDM1-*bla*_L1#3637_; the amount of ceftazidime used was ∼0.3x the MIC value of each strain; **Supplementary Figure S4B**, **Supplementary File S6A**), **(3)** ceftazidime at the same concentrations listed in (2), **(4)** the carrier control (MeOH or DMSO, at equivalent volumes to those used for compound treatments), **(5)** no additions (plain MH medium). Overnight cultures were normalized to 5 × 10⁴ CFU per well, inoculated into 96-well microtiter plates as outlined in **Supplementary Figure S4C**, and incubated for 20 hours at 37 °C. OD_600_ was then recorded for each well. Inhibitory effects were quantified as the fold change between OD_600_ values from condition (2) (candidate compound + ceftazidime) and condition (3) (ceftazidime alone). Compounds producing a statistically significant fold change greater than 1.4 were classified as inhibitor hits. The survival assay was performed in a single technical repeat and three biological replicates. The raw and analyzed data and statistical analyses can be found in **Supplementary File S5**.

#### Refinement of KPC-3 for binding and structural studies

KPC-3 was further purified to homogeneity for differential scanning fluorimetry (DSF) measurements and protein crystallization. Briefly, the dialyzed KPC-3 protein solution (see above) was applied to a HisPur™ Ni-NTA resin (Thermo Fisher Scientific) to remove the cleaved His_8_-SUMO tag and any fusion protein that remained uncleaved. The flow-through fraction, containing KPC-3 without the His_8_-SUMO tag, was further refined to homogeneity by size-exclusion chromatography using a HiLoad™ 16/600 Superdex™ 75 pg column (Cytiva) equilibrated with 20 mM Tris-HCl pH 8.0, 150 mM NaCl. Peak fractions containing the target protein were pooled, concentrated to 20 mg/mL using a Vivaspin 20 centrifugal concentrator with a 10 kDa molecular cut-off (Satorius) according to the manufacturer’s instructions, snap-frozen in liquid nitrogen, and stored at -80 °C until use.

#### Differential scanning fluorimetry (DSF)

A DSF binding assay was performed using purified KPC-3 at a final concentration of 2 µM in 96-well low-profile PCR plates. Compounds were added in 2-fold serial dilutions at final concentrations ranging from 0 to 4 mM, and the mixture was pre-incubated for 10 min at room temperature in 20 mM Tris-HCl (pH 8.0), 150 mM NaCl. SYPRO Orange dye (Thermo Fisher Scientific), supplied as a 5000× stock in DMSO, was added to a final concentration of 10× in each reaction, and fluorescence was recorded during a continuous temperature increase from 25 °C to 95 °C at a rate of 0.04 °C s^-1^ using a LightCycler 480 instrument (Roche). The melting temperature (T_m_) was determined from the first derivative (-dF/dT) of the fluorescence signal, and curve fitting was performed using the Boltzmann equation in the LightCycler 480 Software (v1.4.1.62). Each DSF binding assay condition was performed in technical triplicate yielding consistent T_m_ values.

#### Protein crystallization and X-ray crystallography

Initial crystallization screening was performed using four commercial screens (Nextal Classics, JCSG+, ProComplex, and PEGs Suite; Nextal Biotechnologies) using the sitting-drop vapor diffusion method at room temperature. Crystallization conditions were further optimized in 24-well plates using the hanging-drop vapor diffusion method. Needle-shaped crystals were obtained from reservoir solutions containing 20-25% w/v PEG 4000 and 0.1 M Bicine (pH 8.5-9.5) and reached full size within five days at 20 °C. For inhibitor soaking, crystals were transferred into solutions containing the same concentration of PEG 4000 without Bicine but supplemented with 1 mM inhibitor #61 or 10 mM inhibitor #70 for 10 min. Apo crystals were cryoprotected with 20-25% w/v PEG 4000 and 0.1 M Bicine (pH 8.5-9.5) reservoir solution supplemented with 30% w/v glycerol. Inhibitor-soaked crystals were cryoprotected using a solution containing 20% w/v PEG 400 and 20-25% w/v PEG 4000, but without Bicine, to avoid potential interference between the buffer and the inhibitors. Individual crystals were snap-frozen in liquid nitrogen and stored until data collection.

X-ray diffraction data were collected at beamline 24-ID-E of the Advanced Photon Source (APS) for apo crystals and at beamline 8.2.2 of the Advanced Light Source (ALS) for inhibitor-soaked crystals. The diffraction data were indexed and integrated using the xia2 pipeline (v3.21.1) implementing DIALS (v3.21) within the CCP4 suite (v9.0.010) using the CCP4i2 graphical interface [74–78]. The structures were solved by molecular replacement using PHASER (v2.8.3) with either the coordinates of KPC-2 (PDB:2OV5) or an AlphaFold-predicted model of KPC-3 as the search model [55,79,80]. Model building and refinement were conducted iteratively with Coot (v0.9.8.96) and Phenix (v1.21.2) packages with twinning operation “h,-h-k,-l”, followed by automated re-refinement with PDB-REDO [81–84]. The quality of the final models was assessed using MolProbity [85]. The refined data confirmed Fo-Fc electron density near the active site, prior to further rounds of refinement with inhibitors. The initial 3D coordinates and geometry restraints of the inhibitors were generated using the eLBOW module in Phenix suite SMILES string (Compound #61: “C1=CC(=CC(=C1)B(O)O)C(=O)[H]”, Compound #70: “C1(=CC=C(O1)B(O)O)C=O”) as an input for eLBOW [81]. The statistics for data collection and structure refinement are summarized in **Supplementary Table S11**.

#### Galleria mellonella survival assay

The wax moth model *G. mellonella* was used for *in vivo* survival assays [86]. Individual *G. mellonella* larvae, weighing approximately 200 mg, were randomly allocated to experimental groups (22 larvae per condition; no masking was used). Larvae were separated into Petri dishes with clean Kimwipe bedding and sprayed with 70% ethanol before equilibration to ambient temperature. An overnight culture of *Klebsiella pneumoniae* ZRI was washed three times with PBS and standardized to OD_600_ 0.025 (equivalent to 5.1 x 10^6^ CFU/mL). 10 µL of the bacterial suspension was injected using an auto-injector into the last right abdominal proleg of each larva. 30 minutes after infection, larvae were injected into the last left abdominal proleg with 10 µL of **(1)** the carrier control (PBS + 1.125% v/v DMSO), **(2)** antibiotic only (ceftazidime at 0.05 mg/kg + 1.125% v/v DMSO), **(3)** inhibitor compound only (PBS + inhibitor #61 or #70 at 5 mg/kg), or **(4)** combination treatment, inhibitor compound + antibiotic (ceftazidime at 0.05 mg/kg + inhibitor #61 or #70 at 5 mg/kg). An additional group of 22 larvae was injected twice with 10 µL of PBS at the same intervals as the test larvae, as an injection control; experiments were discontinued and excluded if mortality was greater than 10% in the injection control. All larvae were incubated at 37 °C and mortality was monitored for 48 hours. Death was recorded when larvae turned black due to melanization and did not respond to physical stimulation.

#### Neutropenic murine thigh infection model

A neutropenic murine thigh infection model was established as previously described [87,88]. Details regarding animal husbandry, ethical approval, induction of neutropenia, infection procedures, and treatment regimens are provided below.

#### Animal husbandry

Female, 8-week-old CD1 IGS outbred mice (Charles River) weighing, on average, 28.05 ± 3.16 g (for experiments with inhibitor hit #61) and 30.73 ± 2.60 g (for experiments with inhibitor hit #70) were used for the neutropenic murine thigh infection model. The mice were maintained and overseen by the Animal Resources Center of The University of Texas at Austin. On the day of the experiments, groups of mice were randomly selected, with a maximum of four mice per cage, where they remained for the duration of each procedure (see below). Following induction of neutropenia, and during the infection, and treatment mice were monitored for signs of dehydration and severe disease symptoms.

#### Ethics statement

The animal experiments were performed under the protocol AUP-2024-00138 approved by The University of Texas at Austin, Institutional Animal Care and Use Committee. The University of Texas at Austin animal management program is accredited by the Association for the Assessment and Accreditation of Laboratory Animal Care, International (AAALAC), and meets National Institutes of Health standards as set forth in the Guide for the Care and Use of Laboratory Animals (DHHS Publication No. (NIH) 85–23 Revised 1996).

#### Neutropenic murine thigh infections and treatment

After weighing the animals, mice were rendered neutropenic on day -4 and day -1 of the infection cycle by intraperitoneal injection of 200 µL of filter-sterilized cyclophosphamide (MP Biomedicals) mixed into 70% PBS and 30% PEG 300 (Sigma-Aldrich) at a final concentration of 150 mg/kg. Overnight cultures of *K. pneumoniae* ZRI were washed three times with PBS and standardized to an OD_600_ of 0.0008 (equivalent to 2 x 10^5^ CFU/mL). Mice were anesthetized with 5% v/v isoflurane (Covetrus) and 100 µL of the standardized bacterial suspension was injected intramuscularly into both hind thighs. Baseline bacterial burdens were determined by euthanizing 3 mice and enumerating thigh CFUs 2 hours post-infection. Treatment was initiated at the same time point (2 hours post-infection) and mice were treated every 6 hours over a period of 24 hours (four treatments in total). For inhibitor hit #61, 7 mice were used per treatment condition and were treated with **(1)** the carrier control (PBS + 7.03% v/v DMSO), **(2)** antibiotic only (ceftazidime at 128 mg/kg + 7.03% v/v DMSO), **(3)** inhibitor compound only (PBS + inhibitor #61 at 7.5 mg/kg), or **(4)** combination treatment (ceftazidime at 128 mg/kg + inhibitor #61 at 7.5 mg/kg). For inhibitor hit #70, 5 mice were used per treatment condition and were treated with **(1)** the carrier control (PBS + 4.68% v/v DMSO), **(2)** antibiotic only (ceftazidime at 128 mg/kg + 4.68% v/v DMSO), **(3)** inhibitor compound only (PBS + inhibitor #70 at 10 mg/kg), or **(4)** combination treatment (ceftazidime at 128 mg/kg + inhibitor #70 at 10 mg/kg). For both inhibitor hits, six hours after the final treatment, mice were euthanized by CO_2_, followed by cervical dislocation. Infected thighs from each mouse were harvested and the muscle was debrided, weighed, and transferred into a Beadbeater vial containing 3 mL of PBS and 2 mL of zirconia/silica beads (1.0 mm diameter, Biospec Products). Tissues were homogenized using a Beadbeater homogenizer (Glen Mills). Homogenized suspensions were serially diluted, plated, incubated for 18 hours at 37 °C, and enumerated.

#### Nuclear Magnetic Resonance (NMR)

All NMR spectroscopy data were collected on a Bruker AVANCE III HD 500 MHz spectrometer. The data were processed using the MestReNova software version 16.0.0. ^1^H NMR chemical shifts were referenced to the individual solvent residual peaks of the respective NMR solvents. Stability studies were conducted in DMSO-d_6_ :D_2_O (1:1 ratio) at 37 °C with ^1^H spectra were recorded at 30-min intervals. Inhibitor #61: ^1^H NMR (500 MHz, DMSO-d_6_) δ 10.05 (s, 1H), 8.11 - 7.52 (m, 4H), 4.39 (d, *J* = 17.2 Hz, 2H), 4.18 (d, *J* = 17.2 Hz, 2H), 2.54 (s, 3H). Inhibitor #70: ^1^H NMR (500 MHz, DMSO-d_6_) δ 9.72 (s, 1H), 7.52 (d, *J* = 3.6 Hz, 1H), 7.25 (d, *J* = 3.5 Hz, 1H), 1.31 (s, 12H).

### ADDITIONAL INFORMATION

#### Statistical analysis of experimental data

The total number of performed biological experiments and technical repeats are mentioned in the figure legend of each display item. Biological replication refers to completely independent repetition of an experiment using different biological and chemical materials. Technical replication refers to independent data recordings using the same biological sample.

All recorded MIC values are displayed in the relevant graphs and the bars indicate the median value. We note that in line with recommended practice, MIC results were not averaged. This should be avoided because of the quantized nature of MIC assays, which only inform on bacterial survival for specific antibiotic concentrations and do not provide information for antibiotic concentrations that lie between the tested values. For all other assays, statistical analysis was performed in GraphPad Prism v10.6.1 using either an Ordinary one-way ANOVA, Unpaired t-test, Mantel-Cox Logrank test, Kruskal-Wallis test, Wilcoxon test, or Mann-Whitney U test, as appropriate. Statistical significance was defined as p < 0.05. Detailed information for the statistical analysis performed for each figure is below:

***Figure 2E*:** Comparisons used one-sided Mann-Whitney U tests (matched > control) with Benjamini-Hochberg correction across targets within each comparison family. Significance: *p ≤ 0.05, **p ≤ 0.01, ***p ≤ 0.001, ****p ≤ 0.0001.

***Figure 2F*:** Comparisons used one-sided Mann-Whitney U tests (matched > control) with Benjamini-Hochberg correction across targets within each comparison family. Significance: *p ≤ 0.05, **p ≤ 0.01, ***p ≤ 0.001, ****p ≤ 0.0001.

***Figure 3A*:** One-way ANOVA. For class A serine-β-lactamase KPC-3: p<0.0001 (#54), p<0.0001 (#59), p<0.0001 (#61), p<0.0001 (#62), p<0.0001 (#70), p=0.0009 (#91), p<0.0001 (#94). For class B metallo-β-lactamase L1_#3637_: p<0.0001 (#35), p<0.0001 (#40), p=0.0002 (#108). For class C serine-β-lactamase EC-2: p=0.0005 (#55), p<0.0001 (#61), p<0.0001 (#62), p<0.0001 (#70), p<0.0001 (#91), p<0.0001 (#94). For class D serine-β-lactamase OXA-10: p<0.0001 (#27), p<0.0001 (#58), p<0.0001 (#108). All other statistical analyses are available in **Supplementary File S4**.

***Figure 3B*:** One-way ANOVA. For class A serine-β-lactamase KPC-3: p<0.0001 (#59), p<0.0001 (#61), p<0.0001 (#62), p<0.0001 (#70), p=0.0031 (#91), p<0.0001 (#94). For class B metallo-β-lactamase L1_#3637_: p<0.0001 (#35), p<0.0001 (#42). All other statistical analyses are available in **Supplementary File S5**.

***Figure 6A*:** Mantel-Cox test with the Holm-Šídák correction for multiple comparisons; n=22; p=0.2148 (non-significance) (*K. pneumoniae* ZRI + PBS + DMSO vs. *K. pneumoniae* ZRI + PBS + #61), p=0.7838 (non-significance) (*K. pneumoniae* ZRI + PBS + DMSO vs. *K. pneumoniae* ZRI + ceftazidime + DMSO), p=0.2148 (non-significance) (*K. pneumoniae* ZRI + Ceftazidime + DMSO vs. *K. pneumoniae* ZRI + PBS + #61), p<0.0001 (significance, ****) (*K. pneumoniae* ZRI + PBS + DMSO vs. *K. pneumoniae* ZRI + ceftazidime + #61), p<0.0001 (significance, ****) (*K. pneumoniae* ZRI + ceftazidime + #61 vs. *K. pneumoniae* ZRI + PBS + #61), p<0.0001 (significance, ****) (*K. pneumoniae* ZRI + ceftazidime + DMSO vs. *K. pneumoniae* ZRI + ceftazidime + #61).

***Figure 6B*:** Mantel-Cox test with the Holm-Šídák correction for multiple comparisons; p=0.2384 (non-significance) (*K. pneumoniae* ZRI + PBS + DMSO vs. *K. pneumoniae* ZRI + PBS + #70), p= 0.7838 (non-significance) (*K. pneumoniae* ZRI + PBS + DMSO vs. *K. pneumoniae* ZRI + ceftazidime + DMSO), p=0.2479 (non-significance) (*K. pneumoniae* ZRI + ceftazidime + DMSO vs. *K. pneumoniae* ZRI + PBS + #70), p<0.0001 (significance, ****) (*K. pneumoniae* ZRI + PBS + DMSO vs. *K. pneumoniae* ZRI + ceftazidime + #70), p<0.0001 (significance, ****) (*K. pneumoniae* ZRI + ceftazidime + #70 vs. *K. pneumoniae* ZRI + PBS + #70), p<0.0001 (significance, ****) (*K. pneumoniae* ZRI + ceftazidime + DMSO vs. *K. pneumoniae* ZRI + ceftazidime + #70).

***Figure 6C*:** Kruskal-Wallis test; p>0.9999 (non-significance) (PBS + DMSO vs. #61), p=0.4294 (non-significance) (PBS + DMSO vs. ceftazidime + DMSO), p=0.9550 (non-significance) (#61 vs. ceftazidime + DMSO), p<0.0001 (significance, ****) (PBS + DMSO vs. ceftazidime + #61), p=0.0002 (significance, ***) (#61 vs. ceftazidime + #61), p=0.0314 (significance, *) (ceftazidime + DMSO vs. ceftazidime + #61).

***Figure 6D*:** Kruskal-Wallis test; p=0.1930 (non-significance) (PBS + DMSO vs. #70), p>0.9999 (non-significance) (PBS + DMSO vs. ceftazidime + DMSO), p=0.2225 (non-significance) (#70 vs. ceftazidime + DMSO), p<0.0374 (significance, *) (PBS + DMSO vs. ceftazidime + #70), p<0.0001 (significance, ****) (#70 vs. ceftazidime + #70), p=0.0314 (significance, *) (ceftazidime + DMSO vs. ceftazidime + #70).

***Figure S3D:*** Pairwise comparisons were performed using independent-samples t-tests. All significant differences yielded p<0.0001 (****); non-significant (ns) comparisons indicate p≥0.05.

***Figure S3H:*** Wilcoxon test; p≤0.0001 (significance, ****) (Random subset vs. Important subset). ***Figure S3I:*** Wilcoxon test; p≤0.0001 (significance, ****) (Random subset vs. Important subset). ***Figure S3J:*** Wilcoxon test; p≤0.0001 (significance, ****) (Random subset vs. Important subset).

***Figure S9A:*** Mann-Whitney U tests with Benjamini-Hochberg correction for multiple comparisons; p≥ 0.05 (non-significance) (Negatives vs. KPC-3 hits), p≥ 0.05 (non-significance) (Negatives vs. L1_#3637_ hits), p≥ 0.05 (non-significance) (KPC-3 hits vs. L1_#3637_ hits).

***Figure S9B:*** Mann-Whitney U tests with Benjamini-Hochberg correction for multiple comparisons; p < 0.0001 (significance, ****) (Negatives vs. KPC-3 hits), p≥ 0.05 (non-significance) (Negatives vs. L1_#3637_ hits), p < 0.01 (significance, **) (KPC-3 hits vs. L1_#3637_ hits).

***Figure S9C:*** Mann-Whitney U tests with Benjamini-Hochberg correction for multiple comparisons; p< 0.01 (significance, **) (Negatives vs. KPC-3 hits), p≥ 0.05 (non-significance) (Negatives vs. L1_#3637_ hits), p≥ 0.05 (non-significance) (KPC-3 hits vs. L1_#3637_ hits).

## Data, code and materials availability

PubCheF-1 predictions over the purchasable set of ZINC20 compounds (2024-04-30) can be found at https://www.pubchef.org and at https://doi.org/10.5281/zenodo.21108754. Code used for creating the PubCheF dataset, training and running the PubCheF-1 model, with LRP, can be found at https://www.github.com/kosonocky/pubchef. The X-ray crystallography data generated in this study have been deposited in the Protein Data Bank (PDB) under the following accession codes: KPC-3 apo structure (PDB:36PH); KPC-3 in complex with inhibitor #61 (PDB:36ZC); KPC-3 in complex with inhibitor #70 (PDB:36ZD); residue numbering in the KPC-3 structures follows the standard Ambler scheme with two one-residue gaps at position 58 and 253, in line with previous literature [89,90]. All other data generated during this study that support the findings are included in the manuscript or the Supplementary Information. All materials are available from the corresponding authors upon request.

## SUPPLEMENTARY INFORMATION FOR

### SUPPLEMENTARY FIGURES

**Figure S1.**
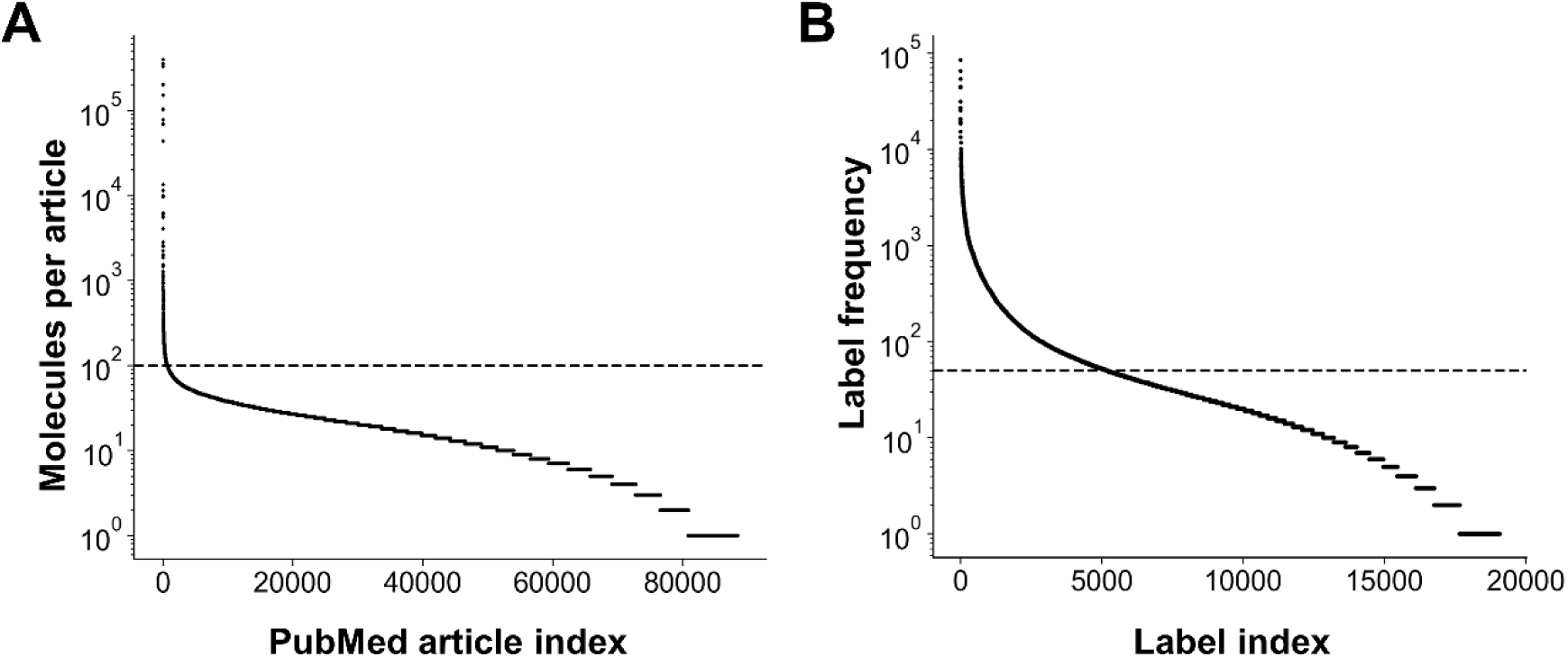
PubCheF dataset statistics. **(A)** Only articles containing fewer than 100 molecules (dashed line) were included in the final dataset. **(B)** Only labels occurring in more than 50 molecules (dashed line) were used for training. The mean number of molecules per label was 425 above this cutoff (included in training) and 16 below the cutoff (excluded from training).

**Figure S2.**
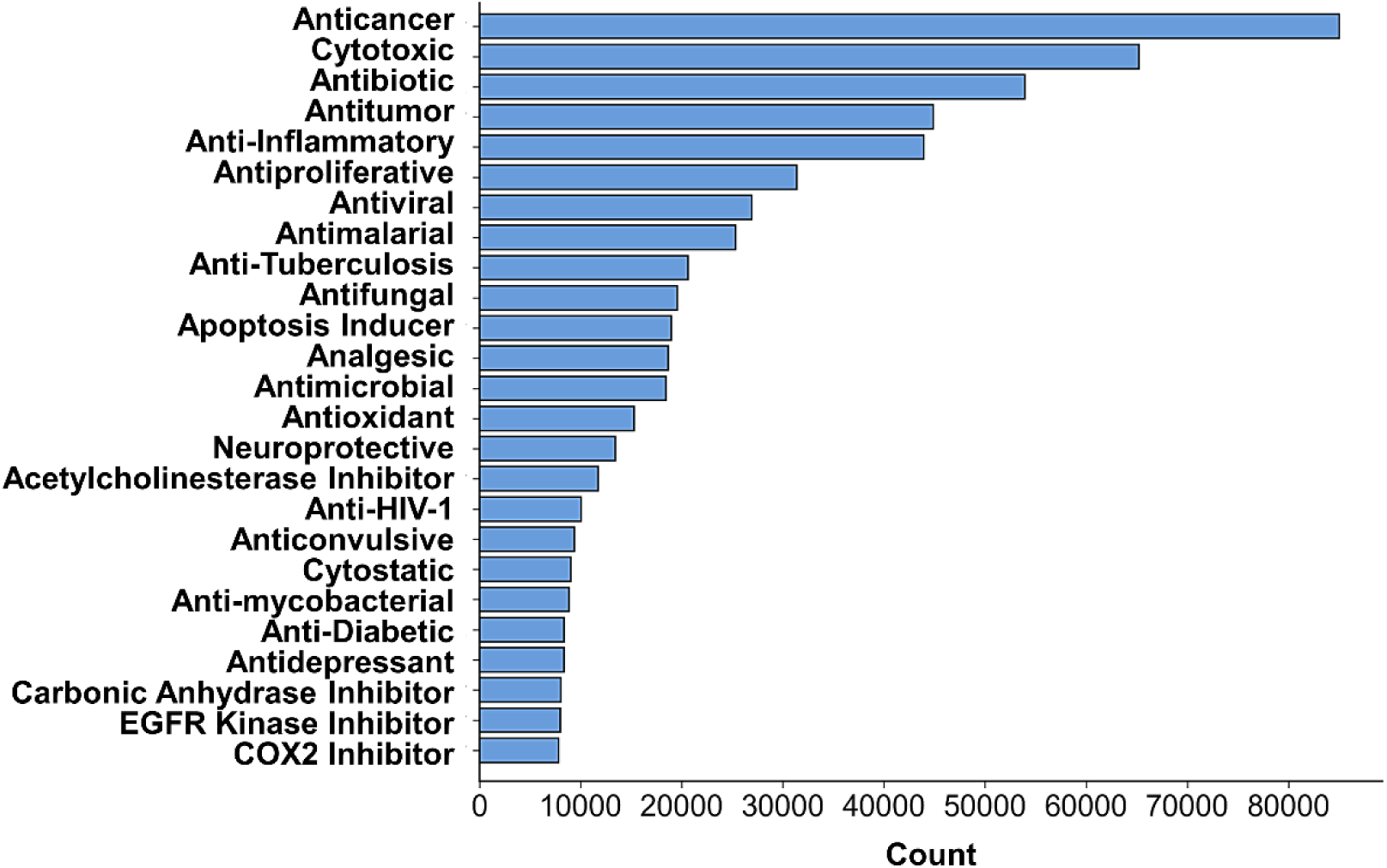
Most frequent labels in the PubCheF dataset. Labels are plotted according to the number of molecules in which they occur.

**Figure S3.**
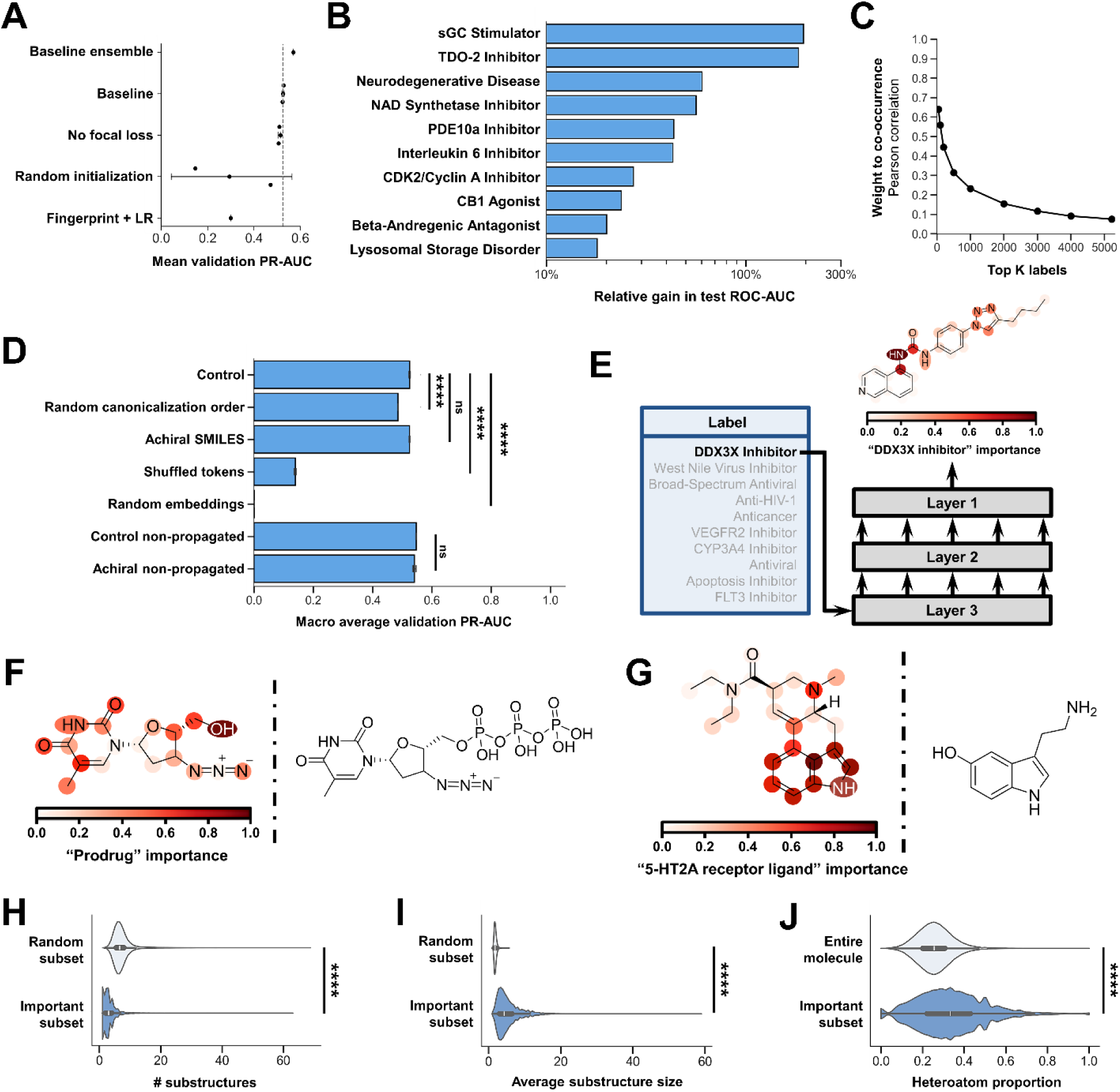
PubCheF-1 model ablations and explainability analyses. (A) Model ablation study using the validation-set macro-averaged PR-AUC. Baseline refers to a single non-ensembled model trained using the parameters in **Supplementary Table S3**. The RoBERTa architecture outperforms simpler models such as logistic regression on RDKit molecular fingerprints, and initialization from a pre-trained ChemBERTa checkpoint outperforms random initialization. Focal loss outperforms cross-entropy loss, which is expected given the label-frequency imbalance (**Supplementary Figure S2**). Ensembling three models trained with different random seeds outperforms a single model. The ensembled model was used throughout the rest of the project. **(B) Effect of single- vs. multi-label training.** For 10 randomly selected labels, PubCheF-1 was re-trained for binary classification of a single label. In all cases, the test-set ROC-AUC was higher when the model was trained on all PubCheF labels than when it was trained only on the label being evaluated, suggesting that multi-label training helps the model gain a general and transferable understanding of chemical function. **(C) Pearson correlations from progressive Mantel tests between the classification-layer weights and label co-occurrence in the dataset.** This indicates that the model may have learned to predict the most frequent labels largely based on the other labels with which they commonly occur, though this trend is diminished for less frequent labels. All points were significant with p < 0.05. **(D) Effect of input representation on validation macro-averaged PR-AUC.** Randomizing canonicalization order for every batch marginally decreases model performance. Removing chirality has no effect on model performance, and this was not due to label propagation. Shuffling token order reduces performance while retaining some predictive power. Randomizing the input embedding to the classifier eliminates predictive performance. Statistical analysis was performed in GraphPad Prism v10.6.1 using independent-samples t-tests; p≤0.0001 (significance, ****), p ≥ 0.05 (non-significance, ns). **(E) Diagram for Layer-wise relevance propagation (LRP)**. LRP calculates which input tokens are most important for predicting a given label. Importance is signified by color intensity; the darker the red color, the greater the weight of the token for prediction. **(F) Importance example for label “Prodrug”**. (**left**) LRP importance for the antiviral zidovudine when predicting the “Prodrug” label. Most of the importance is attributed to a hydroxyl that gets phosphorylated to convert the molecule into its active form (**right**). **(G) Importance example for label “5-HT2A Receptor Ligand”**. (**left**) LRP importance for lysergic acid diethylamide when predicting the “5-HT2A Receptor Ligand” label. Most of the importance is attributed to the indole group that is shared by the receptor’s native ligand, serotonin (**right**). **(H-J) Test set LRP importance analysis**. Across the PubCheF test set, LRP importance was preferentially placed on a small number of large, coherent substructures that are enriched in heteroatoms. Statistical analysis was performed in GraphPad Prism v10.6.1 using a Wilcoxon test; p≤0.0001 (significance, ****).

**Figure S4.**
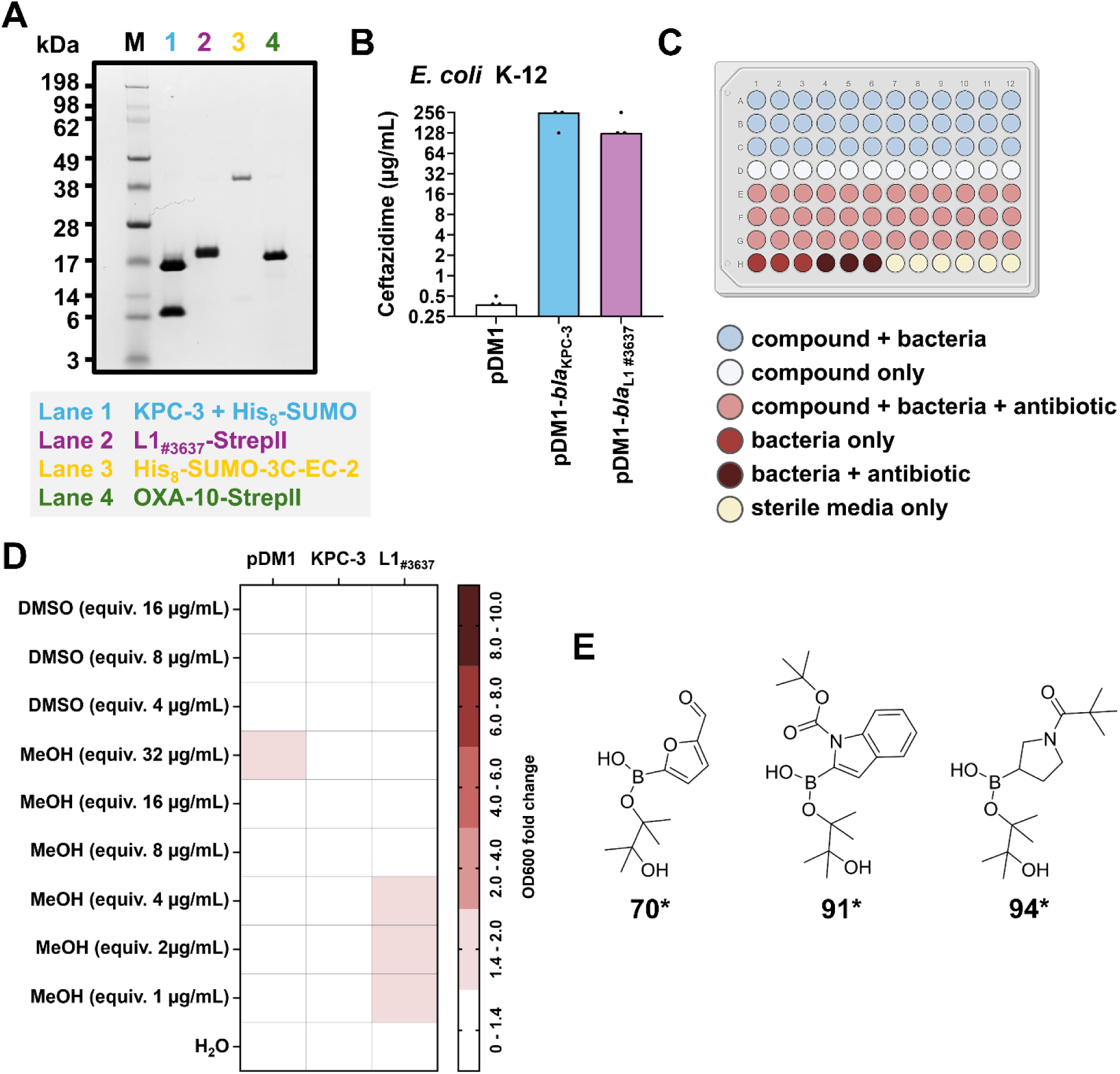
Experimental components for protein target and in-bacteria screening of the PubCheF-1 candidate compounds. **(A)** SDS-PAGE analysis of purified protein samples used for the protein target screen (**Figure 3A**, **Supplementary File S4**). Four representative β-lactamase enzymes were used: the class A serine-β-lactamase KPC-3 (lane 1; the protein was expressed as an N-terminal fusion with a His_8_-SUMO solubility tag, which was proteolytically cleaved but remained in the sample during the screen), the class B metallo-β-lactamase L1_#3637_ (lane 2; tested harboring a C-terminal StrepII tag), the class C serine-β-lactamase EC-2 (lane 3; tested as an N-terminal fusion with a His_8_-SUMO solubility tag), and the class D serine-β-lactamase OXA-10 (lane 4; tested harboring a C-terminal StrepII tag). A representative gel stained for total protein is shown; molecular weight markers (M) are on the left. **(B)** Ceftazidime MIC values for *E. coli* MC1000 (K-12 strain) carrying the empty vector pDM1 (white) or expressing KPC-3 (blue) or L1_#3637_ (purple). These MIC values were used to determine the ceftazidime concentration (∼0.3 x MIC) for the in-bacteria screen (**Figure 3B**, **Supplementary File S5**). Graphs show MIC values (μg/mL) from three biological experiments, each conducted as a single technical repeat. Raw MIC data are available in **Supplementary File S6A**. Consistent with convention, error bars and significance assessment are not shown for MIC panels, as MIC values are discrete. **(C)** The 96-well plate configuration for the in-bacteria screen. Each well is filled with 150 μL of media supplemented with candidate compounds (32 μg/mL), ceftazidime (0.1 μg/mL for MC1000 pDM1; 48 μg/mL for MC1000 pDM1-*bla*_KPC-3_ and MC1000 pDM1-*bla*_L1#3637_; the amount of ceftazidime used was ∼0.3x the MIC value of each strain; **Supplementary Figure S4B**, **Supplementary File S6A**), the carrier control (MeOH or DMSO, at equivalent volumes to those used for compound treatments), and normalized bacterial suspensions, as appropriate. Each plate was incubated for 20 hours at 37 °C and bacterial growth was determined by measuring OD_600_. **(D)** Effects of the carrier controls on the survival of *E. coli* MC1000 harboring the empty vector pDM1 or expressing KPC-3 or L1_#3637_. Data shown represent the fold change of OD_600_ of cells exposed to the carrier controls in the presence of ∼0.3x MIC for ceftazidime for each strain over the OD_600_ of cells exposed only to ∼0.3x MIC of ceftazidime (48 µg/mL ceftazidime for KPC-3 or L1_#3637_ and 0.01 µg/mL for pDM1). MeOH was added at equivalent volumes as for compound treatments of 1, 2, 4, 8, 16, 32 μg/mL and DMSO was added at equivalent volumes as for compound treatments of 4, 8, 16 μg/mL; water was used for conditions that did not require a carrier. Fold change magnitude is indicated by color intensity; the darker the red color the greater the decrease in optical density, while carriers resulting in no change in bacterial survival are shown in white. Overall, bacterial survival is not significantly affected by the presence of the carriers showing that inhibitory activities observed in the in-bacteria screen are not a result of solvent-mediated inhibition or killing. Minor decreases in viability (< 2-fold) are observed for *E. coli* MC1000 carrying pDM1 at 32 μg/mL MeOH and for *E. coli* MC1000 expressing L1_#3637_ at 1, 2, and 4 μg/mL MeOH. None of the candidate compounds hits identified in the in-bacteria screen (**Figure 3B**) dissolved in these conditions. Statistical analysis was performed in GraphPad Prism v10.6.1 using an unpaired T-test or a one-way ANOVA. Raw and analyzed data and statistical analyses can be found in **Supplementary File S5**. **(E)** Chemical structures of the three monosubstituted organoboron compounds, #70*, #91*, and #94*. Because these non-cyclized analogs were not commercially available, the closest structurally related compounds, the cyclized analogs #70, #91, and #94 (**Figure 3C**), in which a dioxaborolane scaffold protects the boronic acid, were selected for purchase and experimental testing instead.

**Figure S5.**
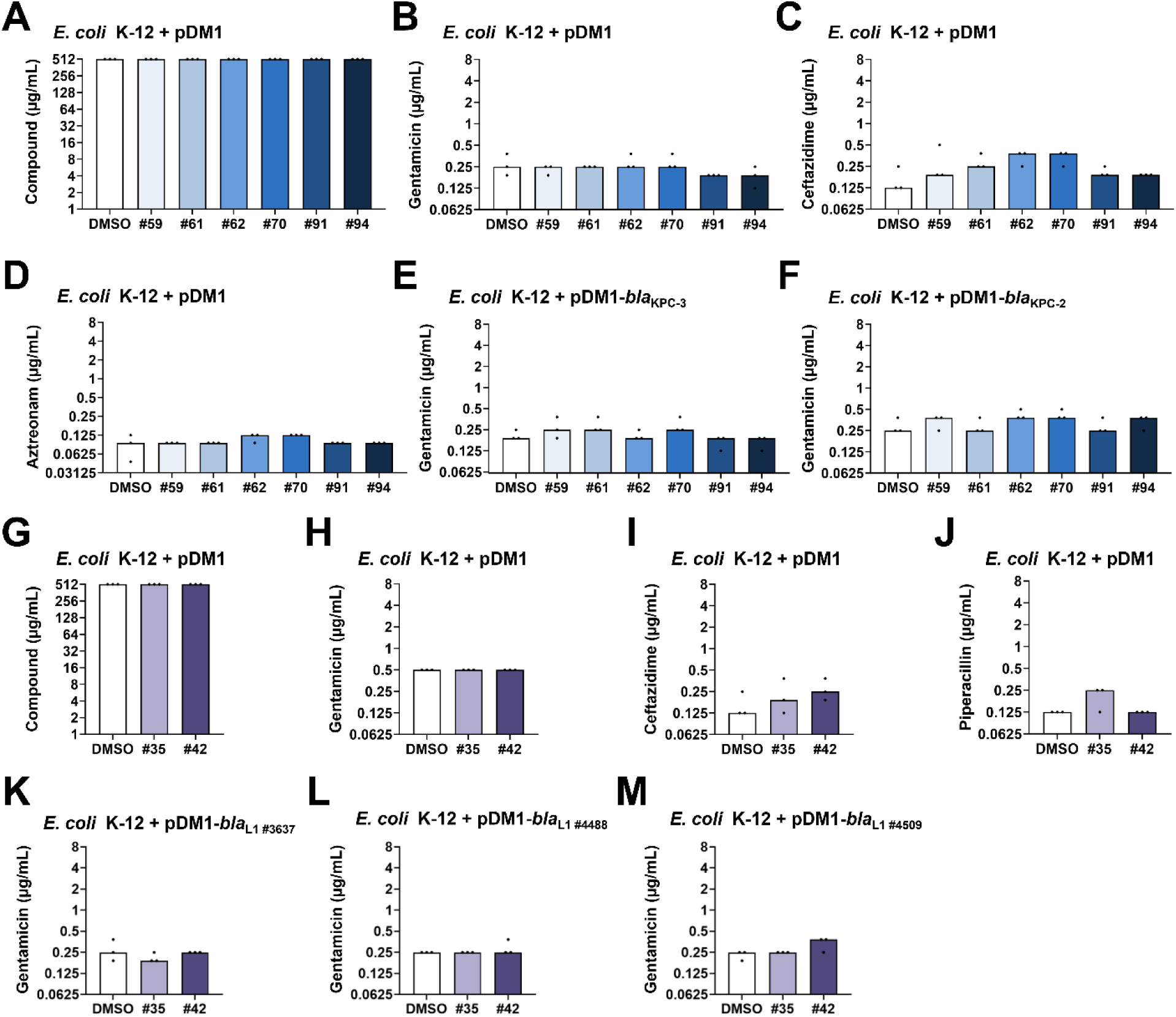
Inhibitor hits do not exhibit β-lactamase-independent activity in *E. coli* MC1000. (**A**) MIC values for candidate compound hits with activity against class-A KPC β-lactamases in *E. coli* MC1000 (K-12 strain) carrying the empty vector pDM1 (blue bars). None of the tested compounds resulted in decreased bacterial survival for concentrations up to 512 μg/mL as compared to the carrier control (DMSO, white bar). **(B)** Gentamicin, **(C)** ceftazidime, and **(D)** aztreonam MIC values for *E. coli* MC1000 carrying pDM1 in the presence of the candidate compound hits with activity against KPC β-lactamases (32 μg/mL, blue bars) or an equal volume of the carrier control (DMSO, white bar). Neither of the tested inhibitor compounds resulted in MIC changes, showing that they do not compromise the general ability of the strain to resist antibiotic stress and that observed MIC effects in the presence of a β-lactamase enzyme are specific to the protein target. Gentamicin MIC values for *E. coli* MC1000 expressing **(E)** KPC-3 or **(F)** KPC-2 remain unchanged in the presence of the tested candidate compound hits (blue bars) as compared to the carrier control (DMSO, white bar), excluding the possibility of off-target effects. **(G)** MIC values for candidate compound hits with activity against class-B L1_#3637_ metallo-β-lactamase in *E. coli* MC1000 carrying pDM1 (purple bars). Neither of the tested compounds resulted in decreased bacterial survival for concentrations up to 512 μg/mL as compared to the carrier control (DMSO, white bar). **(H)** Gentamicin, **(I)** ceftazidime, and **(J)** piperacillin MIC values for *E. coli* MC1000 carrying pDM1 in the presence of the candidate compound hits with activity against L1_#3637_ (32 μg/mL, purple bars) or an equal volume of the carrier control (DMSO, white bar). Neither of the tested inhibitor compounds resulted in MIC changes, showing that they do not compromise the general ability of the strain to resist antibiotic stress and that observed MIC effects in the presence of a β-lactamase enzyme are specific to the protein target. Gentamicin MIC values for *E. coli* MC1000 expressing **(K)** L1_#3637_, **(L)** L1_#4488_, and **(M)** L1_#4509_ remain unchanged in the presence of the tested candidate compound hits (purple bars) as compared to the carrier control (DMSO, white bar), excluding the possibility of off-target effects. Graphs show MIC values (μg/mL) from three biological experiments, each conducted as a single technical repeat. Raw MIC data are available in **Supplementary File S6B**. Consistent with convention, error bars and significance assessment are not shown for MIC panels, as MIC values are discrete.

**Figure S6.**
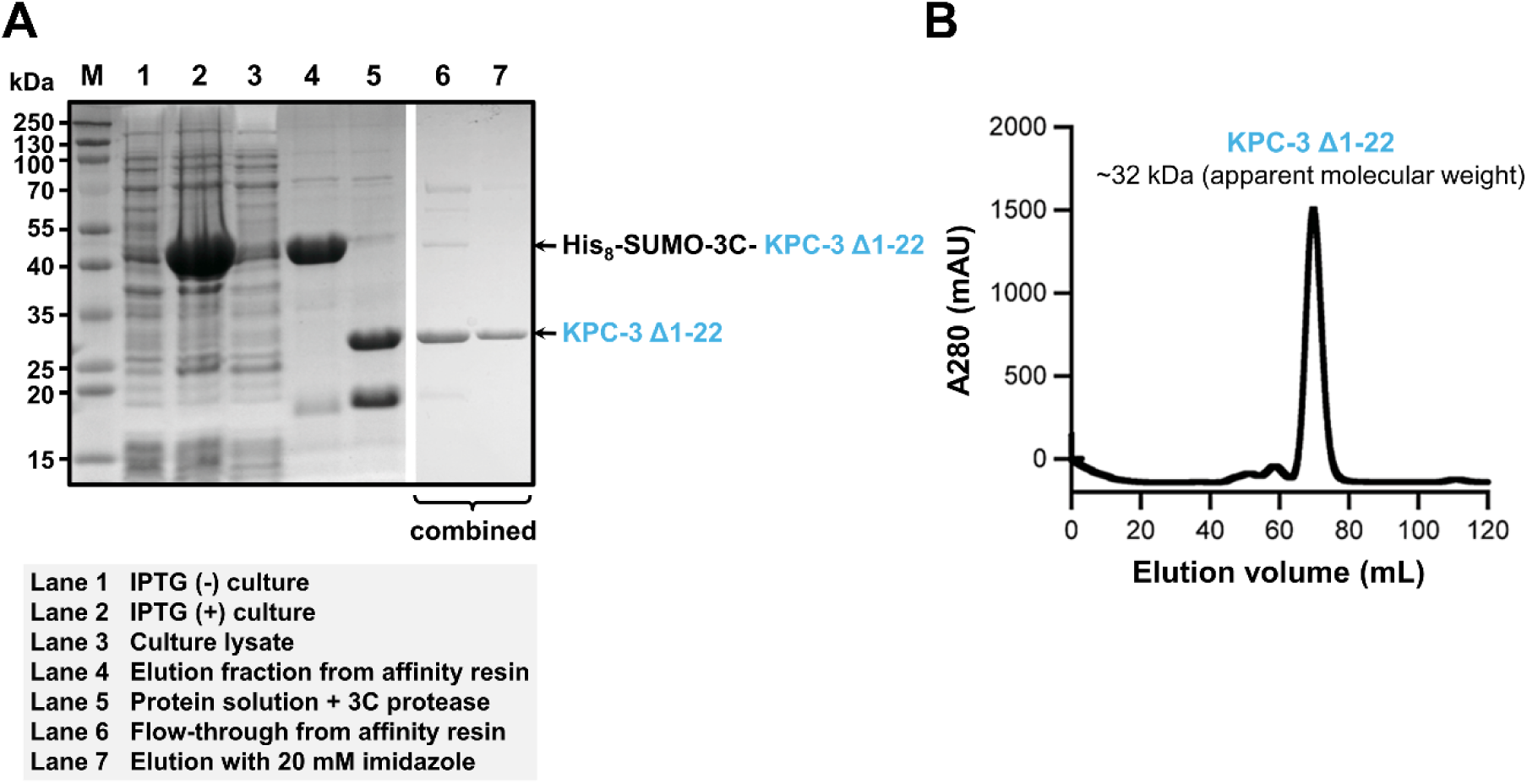
Protein expression and purification of KPC-3 for the binding and structural studies. **(A)** SDS-PAGE analysis detailing the production of recombinant class A serine β-lactamase KPC-3. The enzyme was expressed in the cytoplasm without its signal sequence (Δ1–22) as a fusion with an N-terminal His_8_-SUMO tag followed by an HRV 3C protease cleavage site. Expression (lanes 1 and 2), purification (lanes 3 and 4), proteolytic cleavage (lane 5; sample also shown in **Figure S4A** and used for the target protein screen) and further separation of the mature pure KPC-3 enzyme (lanes 6 and 7) are shown to demonstrate the purity of the final protein sample used for binding and structural studies. A representative gel stained for total protein is shown; molecular weight markers (M) are on the left and gaps indicate where gel lanes were removed. **(B)** Size-exclusion chromatography (SEC) profile of KPC-3 (sample generated from combining lanes 6 and 7 in panel (A)) showing a single, symmetric peak corresponding to monomeric, monodisperse, mature KPC-3 with an apparent molecular weight of 32 kDa.

**Figure S7.**
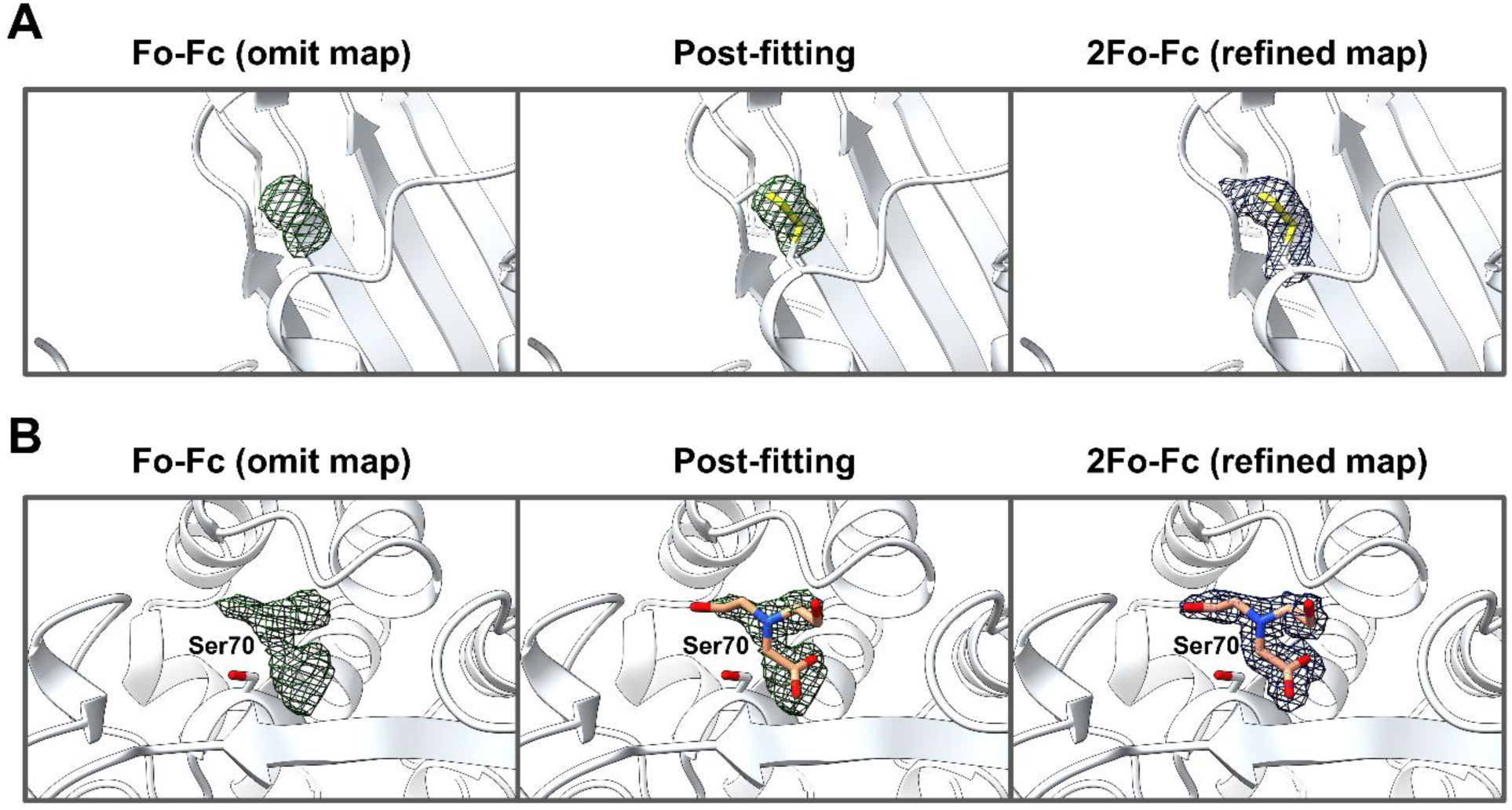
(A) Electron density around the Cys69-Cys238 disulfide bond in the apo KPC-3 structure. (left) Fo–Fc omit map (green mesh, contoured at +2.5σ) calculated before modelling the Cys69-Cys238 disulfide bridge reveals positive difference density. **(middle)** The disulfide bond was built between Cys69 and Cys238 and fitted into the density connecting the two cysteine Sγ atoms. **(right)** The final 2Fo-Fc map (grey mesh, contoured at 1.0σ) around the refined model shows continuous electron density. **(B) Electron density of bicine observed in the apo KPC-3 structure. (left)** Fo-Fc omit map (green mesh, contoured at +2.5σ) calculated prior to ligand modelling shows positive density at the active site. **(middle)** Bicine was built and fitted into the density. **(right)** The final 2Fo–Fc map (grey mesh, contoured at 1.0σ) is displayed around the refined model.

**Figure S8.**
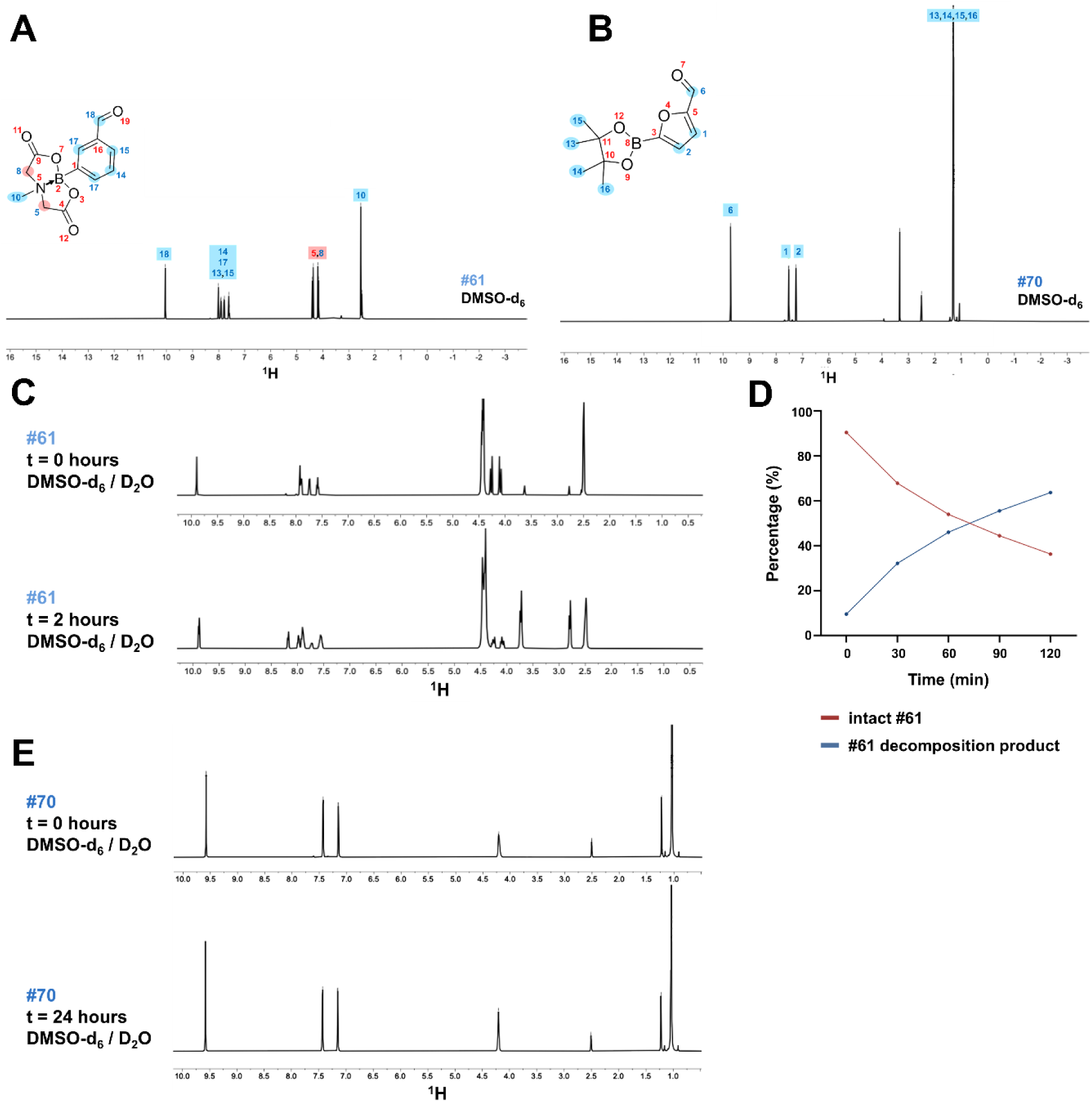
Proton NMR spectral analysis of inhibitor hits #61 and #70. **(A)** Inhibitor hit #61 in deuterated dimethyl sulfoxide (DMSO-*d*_6_): ^1^H NMR (500 MHz, DMSO-*d*_6_) δ 10.05 (s, ^1^H), 8.11 – 7.52 (m, 4H), 4.39 (d, J = 17.2 Hz, 2H), 4.18 (d, J = 17.2 Hz, 2H), 2.54 (s, 3H). **(B)** Inhibitor hit #70 in DMSO-*d*_6_: ^1^H NMR (500 MHz, DMSO-*d*_6_) δ 9.72 (s, 1H), 7.52 (d, J = 3.6 Hz, 1H), 7.25 (d, J = 3.5 Hz, 1H), 1.31 (s, 12H). **(C)** Comparison of the proton NMR spectra of inhibitor hit #61 post dissolution in a 1:1 mixture of DMSO-*d*_6_ and D_2_O at 37 °C, over time (*t*). Spectra at *t* = 0 hours (**top**) and at *t* = 2 hours (**bottom**) reveal spectral changes consistent with cleavage of the dioxazaborocane protecting group, leading us to suggest that inhibitor hit #61 is acting as a prodrug (see main text). **(D)** Time course of the inhibitor hit #61 undergoing hydrolysis to give 3-formylphenylboronic acid as monitored at 30-minute intervals over 2 hours (time (min), starting material %, product %: 0 min, 90.5%, 9.5%; 30 min, 67.9%, 32.1%; 60 min, 54.0%, 46.0%; 90 min, 44.5%, 55.5%; 120 min, 36.3%, 63.7%) **(E)** Comparison of the proton NMR spectra of inhibitor hit #70 at *t* = 0 hours (**top**) and at *t* = 24 hours (**bottom**) post dissolution in a 1:1 mixture of DMSO-*d*_6_ and D_2_O at 37 °C shows no spectral change, a finding interpreted in terms of this compound being stable under these conditions. The deprotection of #70 in the active site of KPC-3 (**Figure 4G**) is, thus inferred to be the result of an interaction between the β-lactamase enzyme and this inhibitor hit, and not due to instability of #70 (see main text for discussion). All NMR spectroscopic data were collected on a Bruker AVANCE III^TM^ HD 500 MHz spectrometer. The data was processed using the MestreNova software version 16.0.0. ^1^H NMR chemical shifts were referenced to the individual solvent residual peaks of the respective NMR solvents.

**Figure S9.**
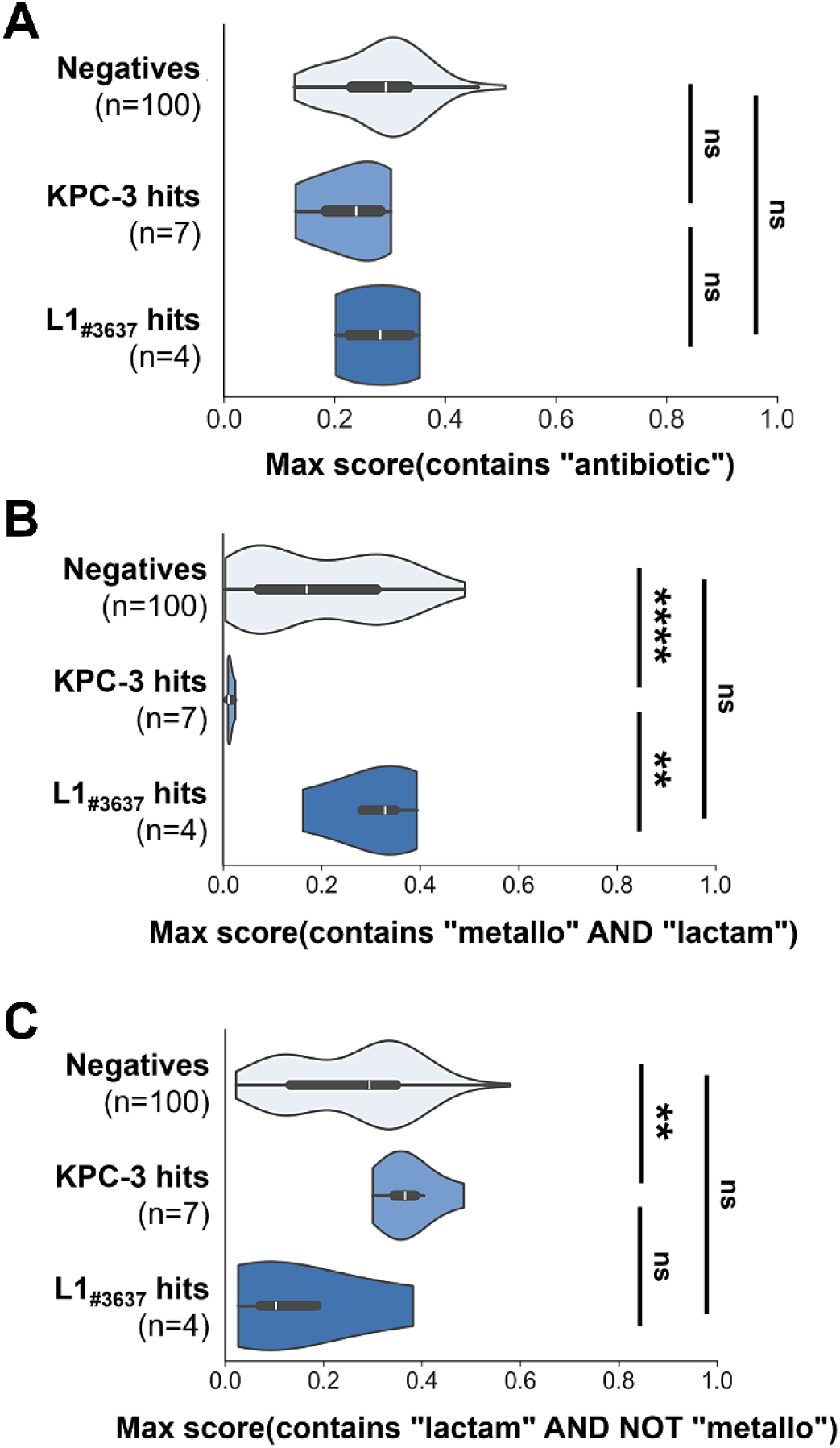
Comparison of PubCheF-1 label scores from the KPC-3 and L1_#3637_ inhibitor hits. **(A)** All purchased molecules were predicted to have the label “Antibiotic” at a PubCheF-1 score above 0.1. However, the “Antibiotic” label was not preferentially enriched among compounds that inhibited the enzymes compared with those that did not. Its prevalence among the candidate compounds may instead arise from correlations with other antibiotic resistance-related labels. **(B)** The KPC-3 serine β-lactamase hits lack PubCheF-1 score mass for labels containing both the substrings “metallo” and “lactam”, whereas the L1_#3637_ metallo-β-lactamase hits are enriched for these labels. **(C)** There is a non-significant enrichment in PubCheF-1 score mass for labels containing “lactam” but not “metallo” among the KPC-3 serine β-lactamase hits. Together, these findings suggest that the model distinguishes between different β-lactamase types, and that both classes were identified in our screen because candidate selection was based only on the “lactam” substring. Statistical analysis was performed in GraphPad Prism v10.6.1 using pairwise comparisons between groups with Mann-Whitney U tests with Benjamini-Hochberg correction for multiple comparisons; p < 0.01 (significance, **), p < 0.0001 (significance, ****); p ≥ 0.05 (non-significance, ns).

**Figure S10.**
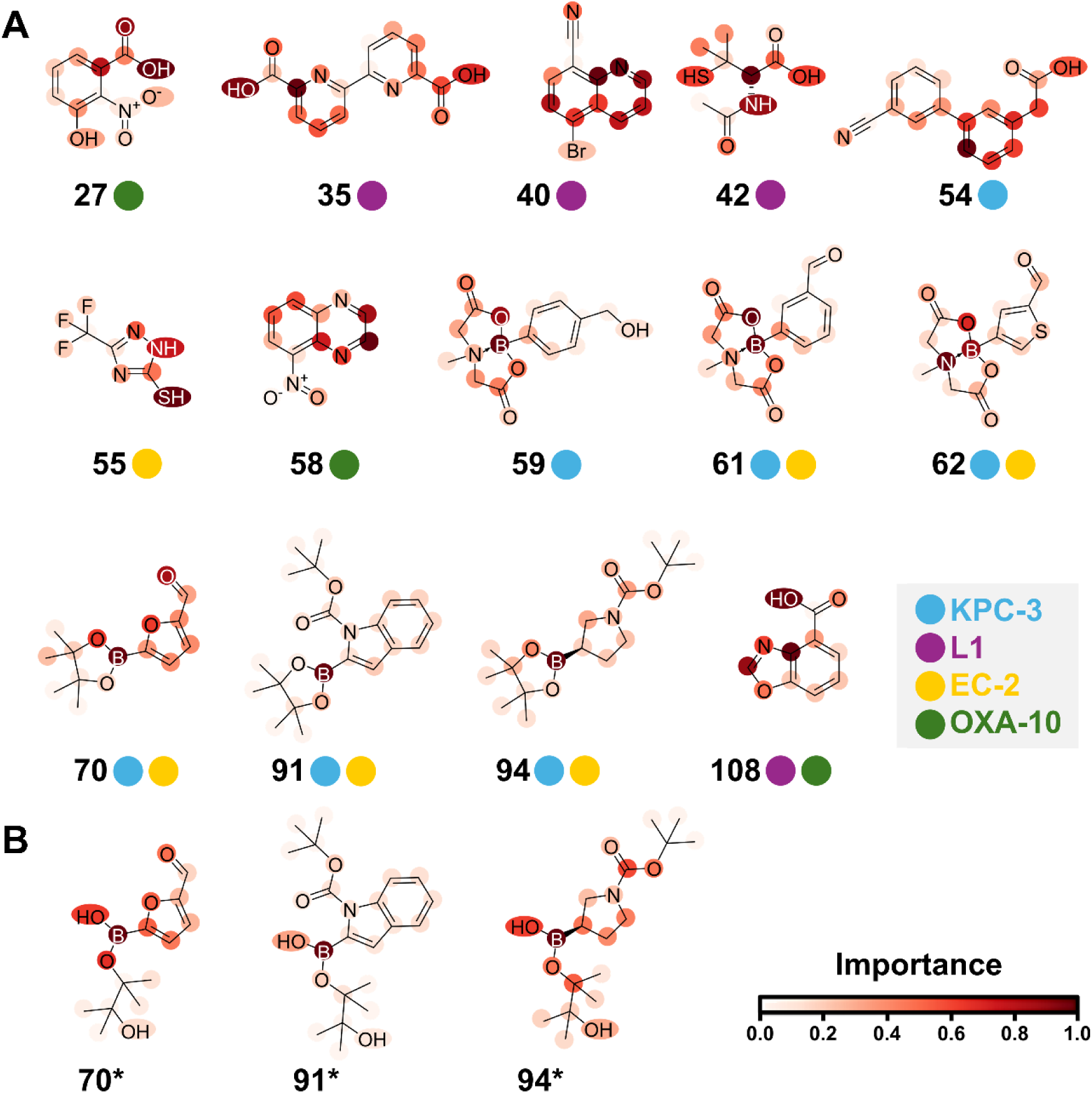
Most important atoms for PubCheF-1 predicting “Beta-Lactamase Inhibitor” or “Metallo-Beta-Lactamase Inhibitor”. LRP-derived importance is signified by color intensity; the darker the red color, the greater the weight of the token for prediction. Importance for the metallo-β-lactamase hits (#35, #40, #42, #108) are shown for the label “Metallo-Beta-Lactamase Inhibitor” label, and the rest are shown for the label “Beta-Lactamase Inhibitor”. For the majority of identified candidate inhibitors, the importance is placed on either hydroxyl, nitrogen, or aromatic ring groups. For all of the boron-containing molecules, boron is the atom with the highest assigned importance value, aligning with the understanding that the boronate groups in these molecules are central to serine β-lactamase inhibition. Panel **(A)** shows the importance for the actual candidate compounds and panel **(B)** shows the importance for the three monosubstituted organoboron compounds (#70*, #91*, and #94*) that were identified by PubChef-1 and were not commercially available; the cyclized analogs #70, #91, and #94, were selected for purchase and experimental testing instead.

**Figure S11.**
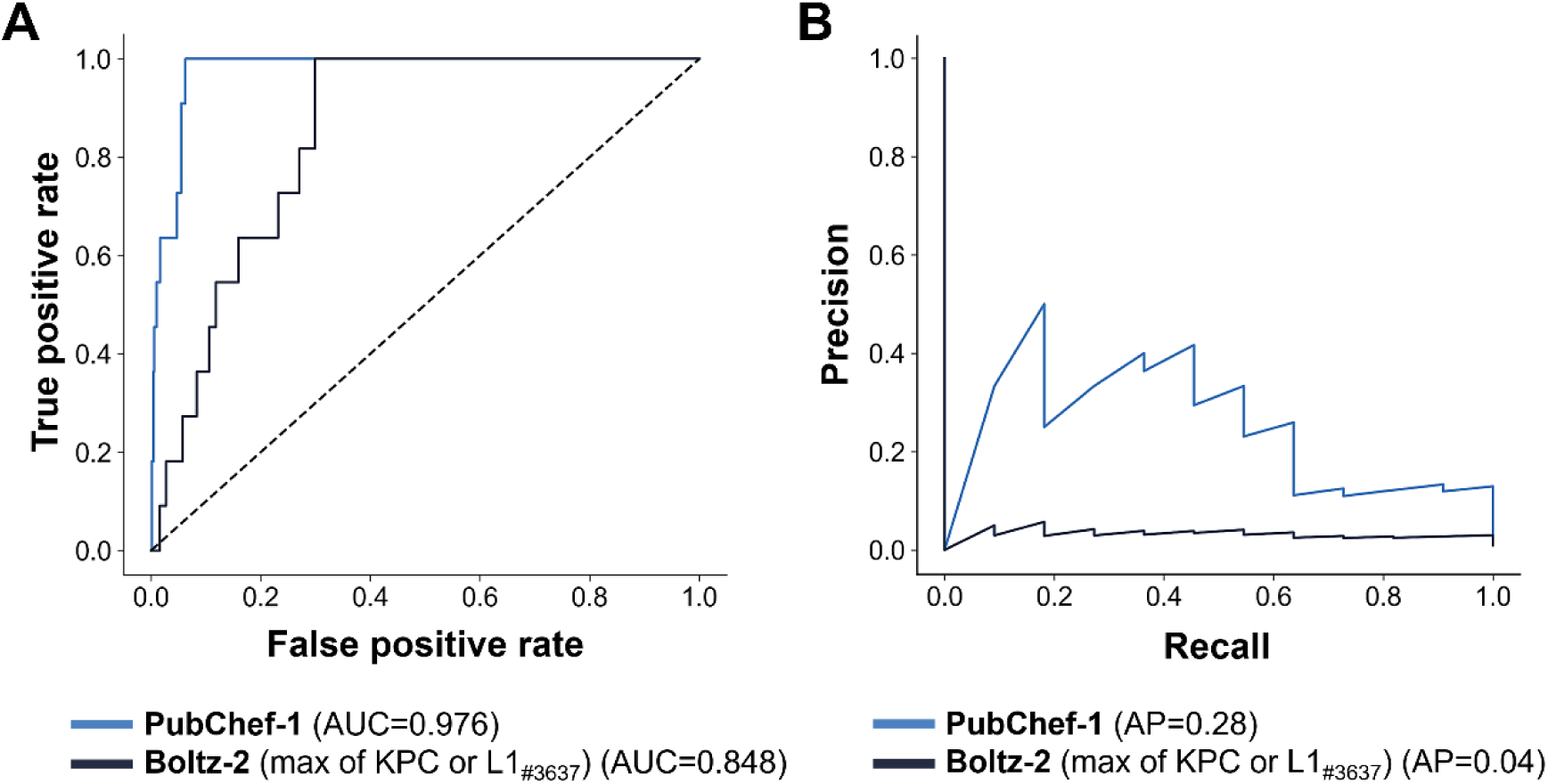
Comparison of PubCheF-1 and Boltz-2 for identifying confirmed β-lactamase inhibitor hits among confirmed negatives and assumed decoys. **(A)** ROC-AUC and **(B)** PR-AUC curves were computed using the PubCheF-1 “max_lactam” score and the Boltz-2 maximum mean “affinity_probability_binary” value across both KPC-3 and L1_#3637_-predicted complexes. Boltz-2 required 71.5 hours to predict affinities for the 2,398 KPC-3 and L1_#3637_ complexes, with 11.85 hours spent on computation on an NVIDIA A100 GPU. PubCheF-1 required 10.3 seconds to generate predictions for the same 1,084 molecules using the same GPU.

**Figure S12.**
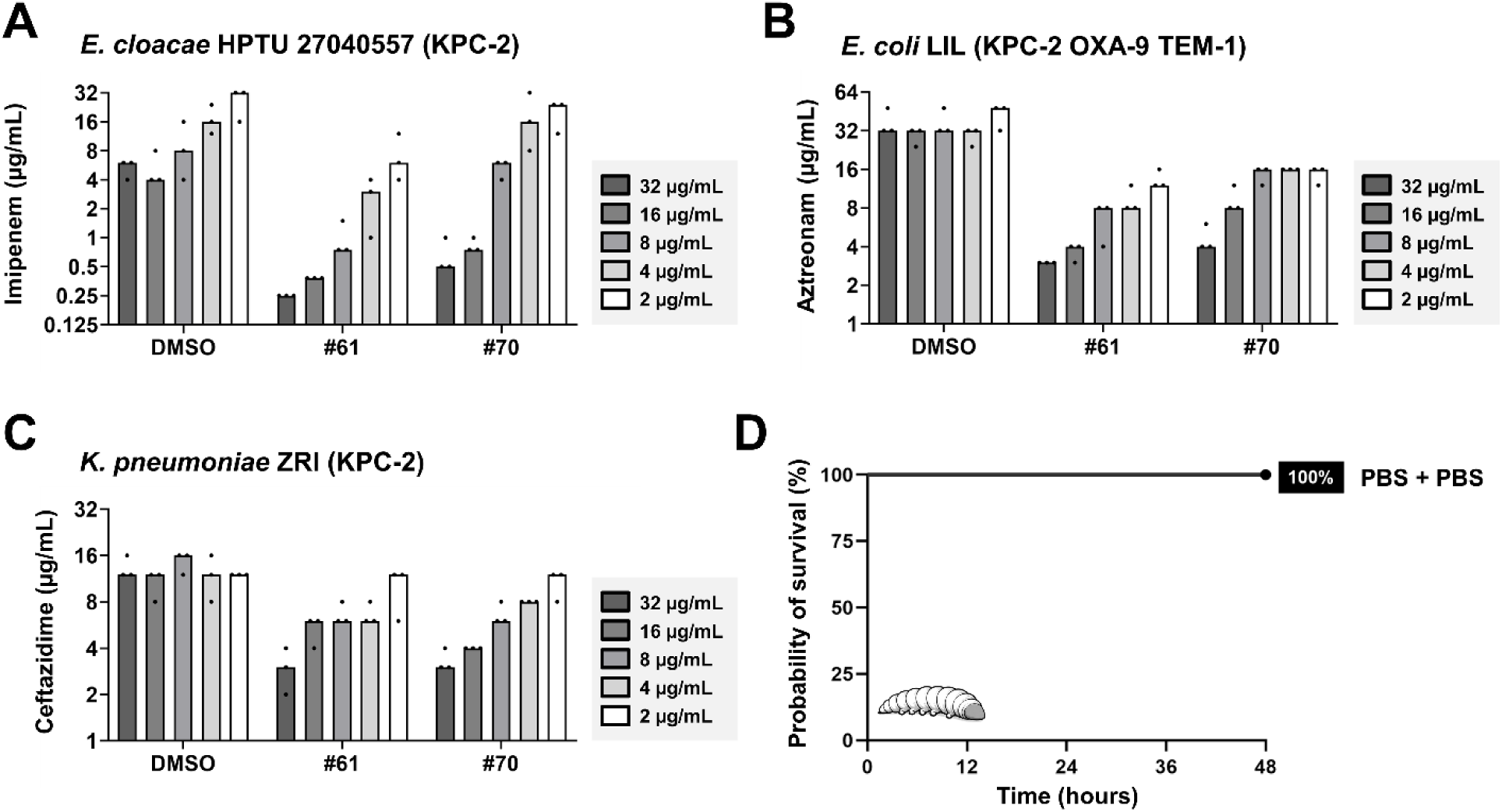
(A-C) Dose-dependent potentiation of β-lactam antibiotics by inhibitor hits #61 and #70 in representative clinical isolates. **(A**) Imipenem MIC values for *E. cloacae* HPTU 27040557 expressing KPC-2 in the presence of decreasing concentrations of inhibitor hits #61 and #70 or the DMSO carrier control (added at an equivalent volume). Both inhibitor hits retain activity down to 16 μg/mL; inhibitor hit #61 shows consistently lower MIC values than the carrier control even at the lowest tested concentration of 2 μg/mL. **(B)** Aztreonam MIC values for *E. coli* LIL expressing KPC-2 in the presence of decreasing concentrations of inhibitor hits #61 and #70 or the DMSO carrier control (added at an equivalent volume). Both inhibitor hits show lower MIC values than the carrier control at all tested concentrations. **(C)** Ceftazidime MIC values for *K. pneumoniae* ZRI expressing KPC-2 in the presence of decreasing concentrations of inhibitor hits #61 and #70 or the DMSO carrier control (added at an equivalent volume). Both inhibitor hits show lower MIC values than the carrier control at all tested concentrations except 2 μg/mL. For panels (A-C), the darkest grey color represents the highest tested concentration of each inhibitor hit (32 μg/mL), with each subsequent bar representing a 2-fold decrease in concentration; the white bars represent the lowest tested concentration (2 μg/mL). Graphs show MIC values (μg/mL) from three biological experiments, each conducted as a single technical repeat. Raw MIC data are available in **Supplementary File S6E**. Consistent with convention, error bars and significance assessment are not shown for MIC panels, as MIC values are discrete. (D) Injection control for the *G. mellonella* infection model. *G. mellonella* larvae show 100% survival when injected twice with PBS using the same procedure as during larval infection and treatment, demonstrating that larval death in **Figure 6A,B** is not a consequence of the injection process. Raw data are available in **Supplementary File S7**.

### SUPPLEMENTARY TABLES

**Table S1.** Fine-tuning improves label quality. Three ChatGPT models (two base, one fine-tuned) were validated on a holdout test set of 100 PubMed articles. The fine-tuned gpt-3.5-turbo model had substantially better precision and vocabulary size than the two base models, but slightly underperformed gpt-4-turbo in recall. The best values are shown in bold.

| MODEL | PRECISION | RECALL | VOCABULARY SIZE |
| --- | --- | --- | --- |
| gpt-3.5-turbo-0613 | 0.61 | 0.82 | 242 |
| gpt-4-turbo-0613 | 0.72 | **0.93** | 254 |
| Fine-tuned gpt-3.5-turbo-0613 | **0.98** | 0.87 | **128** |

**Table S2.** Optimal DBSCAN parameters for clustering PubCheF label embeddings.

| PARAMETER | VALUE |
| --- | --- |
| metric | euclidean distance |
| min_samples | 1 |
| epsilon | 0.410 |

**Table S3.** Model hyperparameters used to train PubCheF-1. Hyperparameter optimization was performed to minimize the validation loss after 20 epochs, resulting in the parameter values shown below.

| PARAMETER | VALUE |
| --- | --- |
| split_type | scaffold |
| dropout_rate | 0.1 |
| max_epochs | 20 |
| batch_size | 128 |
| opt_lr | 1e-3 |
| opt_eps | 1e-8 |
| opt_weight_decay | 1e-2 |
| num_warmup_steps | 2000 |
| pretrained_checkpoint | ChemBERTa-77M-MLM |
| loss_type | focal loss |
| ensemble seeds | 42, 43, 44 |

**Table S4.**
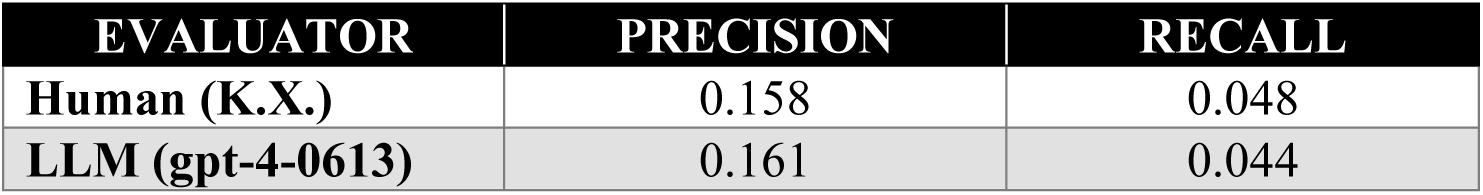
Comparison of large language model (LLM) and human evaluation of chemical function benchmark predictions. For both benchmark sets (PubCheF-test-reduced and OpenTargets-indications), 50 random molecules and their predictions from PubCheF-1, CheF, and MolT5 were obtained (300 examples total).

| EVALUATOR | PRECISION | RECALL |
| --- | --- | --- |
| Human (K.X.) | 0.158 | 0.048 |
| LLM (gpt-4-0613) | 0.161 | 0.044 |

**Table S5.** Benchmark comparison for PubCheF-1, CheF, MolT5, and a ChEMBL-trained PubCheF-1-style model on two chemical-function datasets. Models were evaluated on PubCheF-test-reduced and OpenTargets-indications using an LLM judge to score predictions regardless of output format (**see Materials and Methods**). Hit-rate is the fraction of molecules with at least one correct predicted label. Macro precision is the mean across molecules of the fraction of each molecule’s predicted labels that are correct; because models predict different numbers of labels per molecule (mean labels predicted), precision is computed over each model’s own predicted set rather than at a fixed cutoff. F1 is depressed on OpenTargets-indications because many molecules carry more indications than the number of labels predicted. Molecules receiving no prediction from a given model were assigned a precision of 0 (treated as failures rather than excluded). Molecules for which the judge’s confusion matrix could not be parsed were excluded, yielding 10,282 (PubCheF-test-reduced) and 6,654 (OpenTargets-indications) evaluated molecules from the original 10.3K and 6.7K, respectively. Train test overlaps were computed for both exact molecule and Bemis-Murcko scaffold matches. The high overlap in OpenTargets-indications shows that this benchmark should be interpreted as recall of known pharmacology rather than generalization to novel chemistry. The best values for each benchmark are shown in bold.

| MODEL | BENCHMARK | HIT-RATE | MACRO PRECISION | F1 | MEAN LABELS PREDICTED | TRAIN TEST OVERLAP (Exact) | TRAIN TEST OVERLAP (Scaffold) |
| --- | --- | --- | --- | --- | --- | --- | --- |
| MolT5 | PubCheF-test-reduced | 0.083 | 0.038 | 0.047 | 3.061 | 0.30% | 1.70% |
| CheF | PubCheF-test-reduced | 0.150 | 0.102 | 0.152 | 0.655 | 0.07% | 6.49% |
| PubCheF-1 | PubCheF-test-reduced | **0.501** | **0.455** | **0.530** | 0.907 | 0.00% | 0.00% |
| PubCheF-1 (ChEMBL) | PubCheF-test-reduced | 0.154 | 0.127 | 0.158 | 0.608 | 0.00% | 0.00% |
| MolT5 | OpenTargets-indications | 0.183 | 0.084 | 0.071 | 3.143 | 22.87% | 28.2% |
| CheF | OpenTargets-indications | 0.136 | 0.064 | 0.048 | 1.374 | 0.03% | 33.95% |
| PubCheF-1 | OpenTargets-indications | **0.295** | **0.169** | **0.115** | 2.756 | 43.36% | 61.46% |
| PubCheF-1 (ChEMBL) | OpenTargets-indications | 0.067 | 0.048 | 0.033 | 1.917 | 46.64% | 62.70% |

**Table S6.** Overview of the β-lactamase enzymes used in this study. The “Activity Spectrum” column refers to the hydrolytic spectrum of each tested enzyme; tested enzymes are extended spectrum β-lactamases (ESBL) or carbapenemases. The “Inhibition” column refers to classical inhibitor susceptibility i.e., susceptibility to inhibition by clavulanic acid, tazobactam or sulbactam. Finally, the “Organism” column refers to the bacterial species that most commonly express the tested β-lactamase enzymes. Due to the evolving nomenclature and classification of L1 and L2 enzymes from *S. maltophilia* at the time of this study’s publication, the naming of L1 proteins is based on their strain of origin (see **Tables S7** and **S9**).

| ENZYME | AMBLER CLASS | ACTIVITY SPECTRUM | INHIBITION | ORGANISM |
| --- | --- | --- | --- | --- |
| KPC-3 | A | Carbapenemase | no [1] | *Klebsiella pneumoniae* |
| KPC-2 | A | Carbapenemase | no [1] | *Klebsiella pneumoniae* |
| L1<sub>#3637</sub> | B3 | Carbapenemase | no [2] | *Stenotrophomonas maltophilia* |
| L1<sub>#4488</sub> | B3 | Carbapenemase | unknown | *Stenotrophomonas maltophilia* |
| L1<sub>#4509</sub> | B3 | Carbapenemase | unknown | *Stenotrophomonas maltophilia* |
| EC-2 | C | ESBL | no [3] | *Escherichia coli* |
| OXA-10 | D | ESBL | no [4] | *Pseudomonas aeruginosa* |

**Table S7.**
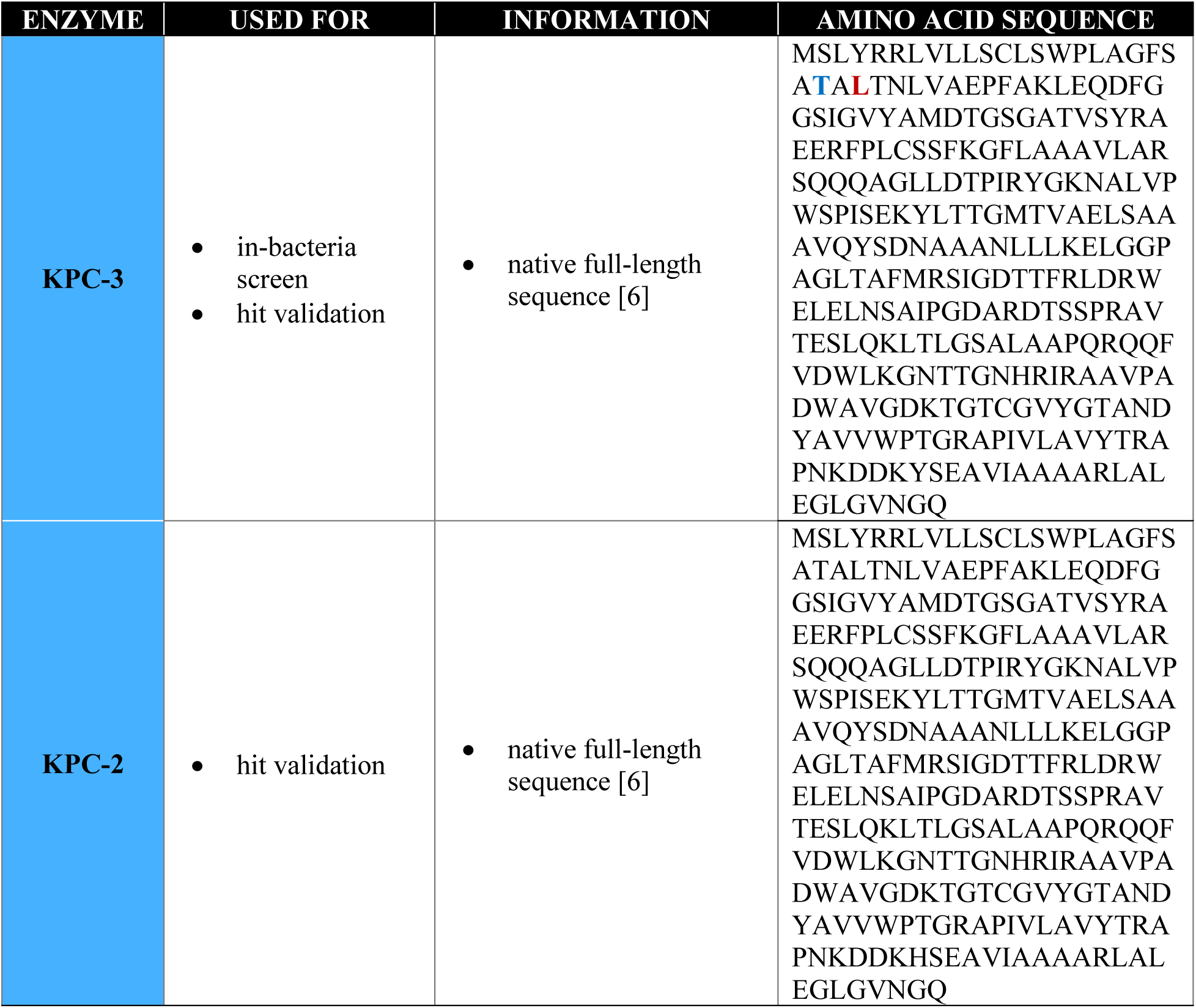
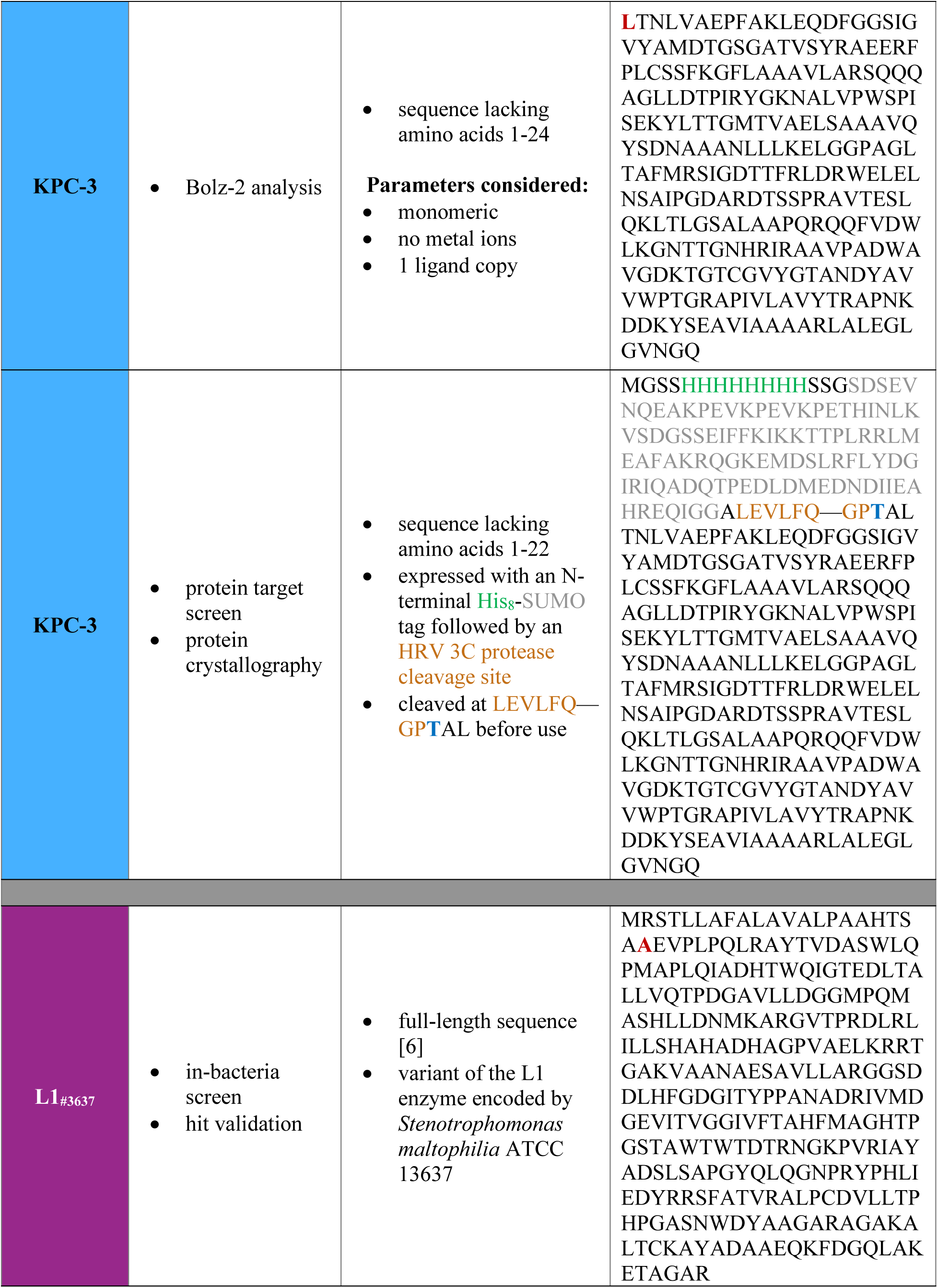
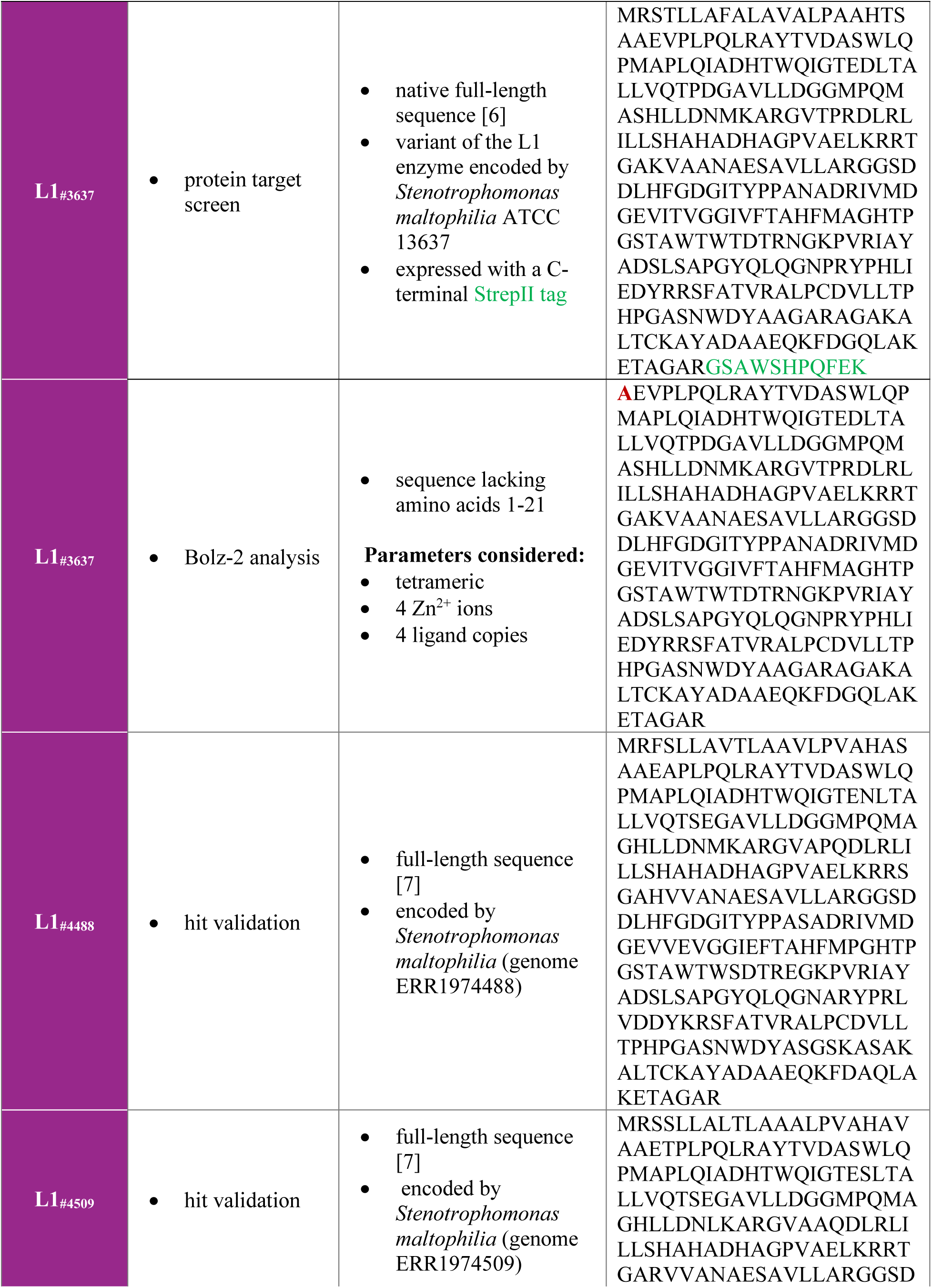
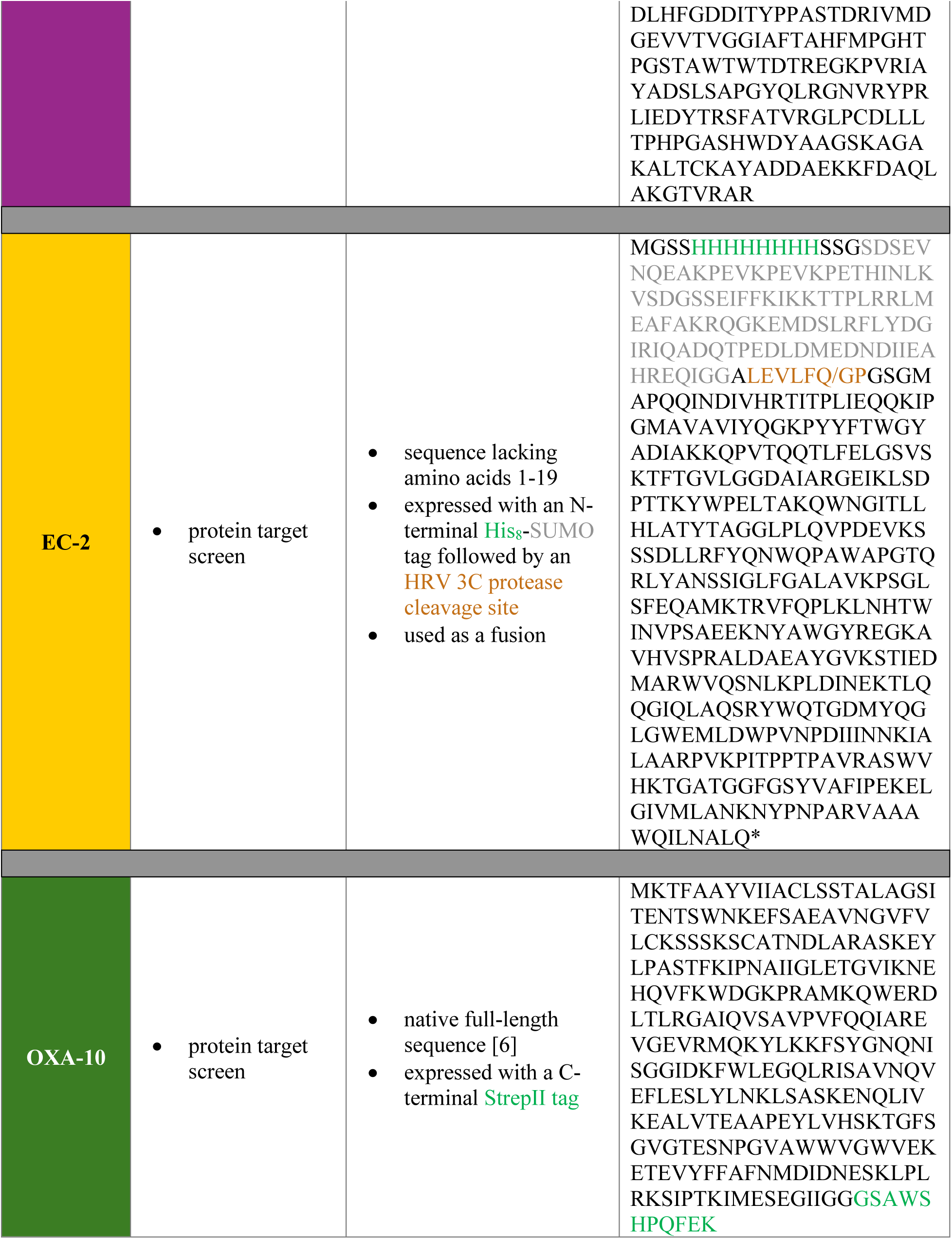
Amino acid sequences for β-lactamase proteins used in this study. The “Used for” column describes the computational or experimental analyses for which each protein sequence was used. The “Information” column details sequence characteristics or computational parameters that were considered during analysis. Enzymes used for in-bacteria screening and hit validation were expressed as full-length sequences including their N-terminal signal sequence that is removed after Sec system translocation [5]. For some enzymes (KPC-3, L1_#3637_ and EC-2), the mature protein sequence (lacking 22 (or 24), 21, and 19 amino acids, respectively) was used for computational analyses and for protein production to be used for the protein target screen or for crystallization. The start of the mature sequence is indicated within the full-length protein in red or blue letters. For constructs expressed with affinity or solubility tags these are color-coded accordingly. Due to the evolving nomenclature and classification of L1 and L2 enzymes from *S. maltophilia* at the time of this study’s publication, the naming of L1 proteins is based on their strain of origin, which is stated in the “Information” column.

**Table S8.** Bacterial strains used in this study. All listed isolates are clinical strains. “FNRCAR” refers to the French National Reference Centre for Antibiotic Resistance in Le Kremlin-Bicêtre, France.

| NAME | DESCRIPTION | SOURCE |
| --- | --- | --- |
| ***Escherichia coli*** |  |  |
| DH5α | F<sup>-</sup> *endA1 glnV44 thi-1 recA1 relA1 gyrA96 deoR nupG purB20* ϕ80*dlacZ*ΔM15 Δ(*lacZYA-argF*)U169 *hsdR17*(r<sub>K</sub><sup>-</sup>m<sub>K</sub><sup>+</sup>) λ<sup>-</sup> | [8] |
| CC118λpir | F<sup>-</sup> *araD139* Δ(*ara, leu*)7679 Δ*lacX74 phoA20 galE thi rspE rpoB argE*(Am) *recA1* λpir | [9] |
| MC1000 | F<sup>-</sup> *araD139* Δ(*ara-leu*)7697 Δ*lacX74 galU galK strA* λ<sup>-</sup> | [10] |
| BL21 (DE3) | F<sup>-</sup> *ompT11 galB dcm-6 lon-11 hsdS<sub>B</sub>*(r<sub>B</sub><sup>-</sup>m<sub>B</sub><sup>-</sup>) λ(DE3 [*lacI lacUV5-T7p07 ind1 sam7 nin5*]) [*malB*<sup>+</sup>]<sub>K-12</sub>(λ<sup>S</sup>) | New England Biolabs |
| BL21 SHuffle T7 Express | F<sup>-</sup> *ompT11 galB dcm-6 lon-11 hsdS<sub>B</sub>*(r<sub>B</sub><sup>-</sup>m<sub>B</sub><sup>-</sup>) *trxB1 gor522 ahpC\** Δ*attB::pnprt-dsbC* λ(DE3 [*lacI lacUV5-T7p07 ind1 sam7 nin5*]) [*malB*<sup>+</sup>]<sub>K-12</sub>(λ<sup>S</sup>) | New England Biolabs |
| **Clinical isolates** |  |  |
| *Enterobacter cloacae* BM15 | *bla*<sub>KPC-2</sub> *bla*<sub>TEM-1</sub> *bla*<sub>OXA-9</sub> | [11] |
| *Enterobacter cloacae* HGM 83048 | *bla*<sub>KPC-2</sub> *bla*<sub>TEM-1</sub> | [11] |
| *Enterobacter cloacae* HPTU 27040557 | *bla*<sub>KPC-2</sub> | [11] |
| *Enterobacter cloacae* CFVL | *bla*<sub>KPC-2</sub> *bla*<sub>TEM-3</sub> | [11] |
| *Klebsiella pneumoniae* S22 | *bla*<sub>KPC-3</sub> | FNRCAR |
| *Klebsiella pneumoniae* COQ | *bla*<sub>KPC-2</sub> | FNRCAR |
| *Klebsiella pneumoniae* ZRI | *bla*<sub>OKP-A</sub> *bla*<sub>OXA-1</sub> *bla*<sub>TEM-1</sub> *bla*<sub>KPC-2</sub> | FNRCAR |
| *Klebsiella pneumoniae* S32 | *bla*<sub>KPC-2</sub> | FNRCAR |
| *Klebsiella pneumoniae* S31 | *bla*<sub>KPC-2</sub> | FNRCAR |
| *Klebsiella pneumoniae* S5 | *bla*<sub>KPC-2</sub> | FNRCAR |
| *Klebsiella pneumoniae* S15 | *bla*<sub>KPC-2</sub> | FNRCAR |
| *Klebsiella pneumoniae* ST234 | *bla*<sub>KPC-2</sub> *bla*<sub>SHV-27</sub> | [12] |
| *Citrobacter freundii* BM19 | *bla*<sub>KPC-2</sub> | [11] |
| *Citrobacter freundii* CNR27B5 | *bla*<sub>KPC-3</sub> | FNRCAR |
| *Citrobacter freundii* HPTU | *bla*<sub>KPC-2</sub> *bla*<sub>TEM-1</sub> | FNRCAR |
| *Escherichia coli* BM16 | *bla*<sub>KPC-2</sub> *bla*<sub>TEM-1b</sub> | [11] |
| *Escherichia coli* COL | *bla*<sub>KPC-2</sub> *bla*<sub>TEM-1</sub> *bla*<sub>CTX-M-9</sub> | [11] |
| *Escherichia coli* DIN | *bla*<sub>KPC-2</sub> | [11] |
| *Escherichia coli* IFI | *bla*<sub>KPC-2</sub> | FNRCAR |
| *Escherichia coli* LIL-1 | *bla*<sub>KPC-2</sub> *bla*<sub>TEM-1</sub> *bla*<sub>OXA-9</sub> | [11] |
| *Escherichia coli* S34 1F7 | *bla*<sub>KPC-2</sub> | FNRCAR |
| *Stenotrophomonas maltophilia* AMM | *bla*<sub>L2</sub> *bla*<sub>L1</sub> | [13] |
| *Stenotrophomonas maltophilia* AMM Δ*bla*<sub>L2</sub> | *bla*<sub>L1</sub> | This study |

**Table S9.** Plasmids used in this study.

| NAME | DESCRIPTION | SOURCE |
| --- | --- | --- |
| pDM1 | pDM1 vector (GenBank MN128719), p15A *ori*, *Ptac* promoter, MCS, Tet<sup>R</sup> | Mavridou lab stock |
| pDM1-*bla*<sub>KPC-2</sub> | *bla*<sub>KPC-2</sub> cloned into pDM1, Tet<sup>R</sup> | [6] |
| pDM1-*bla*<sub>KPC-3</sub> | *bla*<sub>KPC-3</sub> cloned into pDM1, Tet<sup>R</sup> | [6] |
| pDM1-*bla*<sub>KPC-3</sub>-StrepII | *bla*<sub>KPC-3</sub> encoding KPC-3 with a C-terminal StrepII tag cloned into pDM1, Tet<sup>R</sup> | [6] |
| pDM1-*bla*<sub>L1</sub> #3637 | *bla*<sub>L1</sub> #3637, a variant of the L1 enzyme encoded by *Stenotrophomonas maltophilia* ATCC 13637 cloned into pDM1, Tet<sup>R</sup> | [6] |
| pDM1-*bla*<sub>L1</sub> #3637-StrepII | *bla*<sub>L1</sub> #3637 encoding L1#3637 with a C-terminal StrepII tag cloned into pDM1, Tet<sup>R</sup> | [6] |
| pDM1-*bla*<sub>L1</sub> #4488 | *bla*<sub>L1</sub> #4488 encoding the L1 enzyme of *Stenotrophomonas maltophilia* (genome ERR1974488) cloned into pDM1, Tet<sup>R</sup> | This study |
| pDM1-*bla*<sub>L1</sub> #4509 | *bla*<sub>L1</sub> #4509 encoding the L1 enzyme of *Stenotrophomonas maltophilia* (genome ERR1974509) cloned into pDM1, Tet<sup>R</sup> | This study |
| pDM1-*bla*<sub>OXA-10</sub>-StrepII | *bla*<sub>OXA-10</sub> encoding OXA-10 with a C-terminal StrepII tag cloned into pDM1, Tet<sup>R</sup> | [6] |
| pET28a(+)-His8-SUMO-HRV3C-*bla*<sub>KPC-3</sub> Δ1-22 | *bla*<sub>KPC-3</sub> encoding KPC-3 lacking residues 1-22 cloned in a pET28a(+)-derived vector with an N-terminal His8-SUMO tag followed by an HRV 3C protease cleavage site, Kan<sup>R</sup> | This study |
| pET28a(+)-His8-SUMO-HRV3C-*bla*<sub>EC-2</sub> Δ1-19 | *Bla*<sub>EC-2</sub> encoding EC-2 lacking residues 1-19 cloned in a pET28a(+)-derived vector with an N-terminal His8-SUMO tag followed by an HRV 3C protease cleavage site, Kan<sup>R</sup> | This study |
| pKNG101 | Gene replacement suicide vector, *ori*R6K, *ori*TRK2, *sacB*, (template for the *strAB* cassette), Str<sup>R</sup> | [14] |
| pKNG101-*bla*<sub>L2</sub> | PCR fragment containing the regions upstream and downstream *S. maltophilia* AMM *bla*<sub>L2</sub> gene cloned in pKNG101; when inserted into the chromosome, the strain is a merodiploid for *bla*<sub>L2</sub> mutant, Str<sup>R</sup> | This study |
| pRK600 | Helper plasmid, ColE1 *ori*, *mob*RK2, *tra*RK2, Cam<sup>R</sup> | [15] |

**Table S10.**
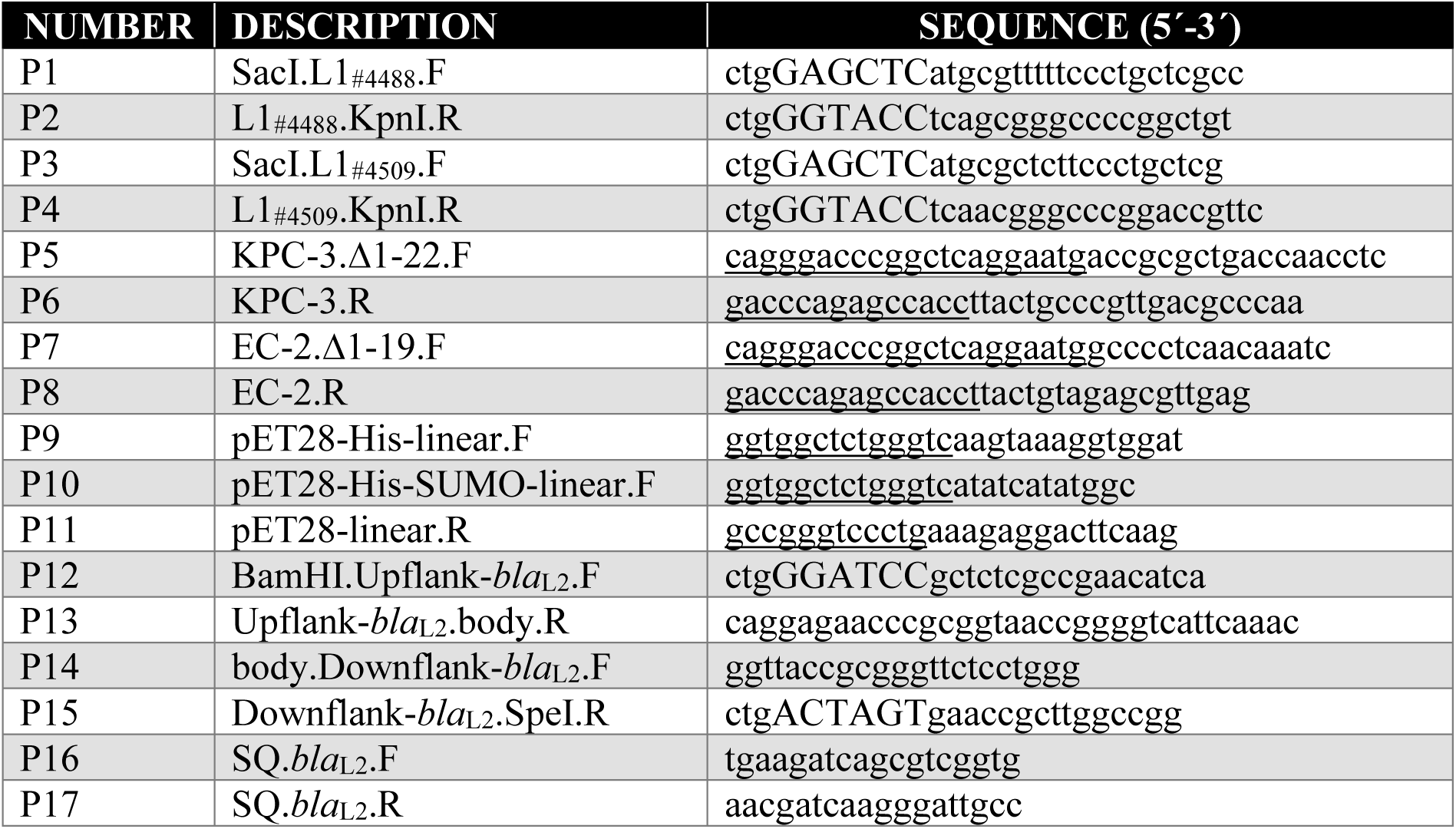
Oligonucleotide primers used in this study. The “Description” column provides basic information on the primer design (restriction enzyme used for cloning, encoded protein or deleted gene, forward or reverse orientation of the primer (F or R); SQ stands for sequencing primers). Restriction sites are indicated in capital letters while underlined sequences indicate Gibson assembly overlap regions.

**Table S11.** X-ray crystallographic data collection and refinement statistics. * Values for the corresponding parameters in the outermost shell are shown in parentheses. ^ϒ^CC1/2 is the Pearson correlation coefficient for a random half of the data, the two numbers represent the lowest and highest resolution shell, respectively. ^±^R_free_ is the R_work_ value calculated for about 10% of the reflections randomly selected and omitted from refinement. ^MolProbity score is calculated by combining clashscore with rotamer and Ramachandran percentage and is scaled based on X-ray resolution. The percentage is calculated with 100^th^ percentile as the best and the 0^th^ percentile as the worst among structures of comparable resolution.

| PDB ENTRY | KPC-3 apo<br>36PH | KPC-3·+ #61<br>36ZC | KPC3·+ #70<br>36ZD |
| --- | --- | --- | --- |
| **Data Collection** |  |  |  |
| Space group | P 3<sub>1</sub> | P 3<sub>1</sub> | P 3<sub>1</sub> |
| **Cell Dimension** |  |  |  |
| Resolution (Å) | 58.01 – 2.40 (2.44 – 2.40)* | 57.79 – 2.52 (2.56 – 2.52) | 58.00 – 2.32 (2.36 – 2.32) |
| Unit cell: a, b, c (Å) | 116.02, 116.02, 51.78 | 115.59, 115.59, 52.50 | 116.00, 116.00, 51.56 |
| Unit cell: $\alpha$, $\beta$, $\gamma$ (°) | 90, 90, 120 | 90, 90, 120 | 90, 90, 120 |
| $R_{\text{sym}} / R_{\text{pim}}$ | 0.186 (0.977) / 0.068 (0.369) | 0.123 (0.911) / 0.060 (0.440) | 0.135 (0.709) / 0.064 (0.332) |
| $CC1/2^Y$ | 0.989 (0.672) | 0.995 (0.611) | 0.991 (0.769) |
| I / $\sigma$ | 8.4 (1.0) | 14.5 (1.5) | 10.2 (1.3) |
| Completeness (%) | 100.0 (99.9) | 100.0 (99.9) | 100.0 (100.0) |
| Redundancy | 8.4 (8.0) | 5.2 (5.2) | 5.35 (5.5) |
| **Refinement** |  |  |  |
| Resolution (Å) | 58.01 – 2.40 (2.46 – 2.40) | 57.79 – 2.52 (2.58 – 2.52) | 58.00 – 2.32 (2.38 – 2.32) |
| No. reflections | 30404 (2145) | 26482 (1869) | 33591 (2388) |
| $R_{\text{work}}$ | 0.1565 (0.2451) | 0.1614 (0.2310) | 0.2000 (0.2700) |
| $R_{\text{free}}^{\pm}$ | 0.1898 (0.2426) | 0.1997 (0.2593) | 0.2350 (0.2878) |
| **No. Atoms** |  |  |  |
| Protein | 5808 | 5808 | 5808 |
| Ligand | 33 | 22 | 10 |
| Solvent | 3 | 7 | 5 |
| **B- factors (Å<sup>2</sup>)** |  |  |  |
| Protein | 52.5 | 56.1 | 41.0 |
| Ligand | 41.9 | 36.2 | 23.0 |
| Solvent | 37.6 | 40.3 | 27.7 |
| **R.m.s deviations** |  |  |  |
| Bond lengths (Å) | 0.009 | 0.011 | 0.009 |
| Bond angles (°) | 1.15 | 1.47 | 1.12 |
| **Ramachandran plot** |  |  |  |
| Favored (%) | 98.6 | 98.2 | 98.4 |
| Allowed (%) | 1.4 | 1.8 | 1.6 |
| Outliers (%) | 0.00 | 0.00 | 0.00 |
| **Molprobity score<sup>^</sup>** | 1.35 (100<sup>th</sup> percentile) | 1.50 (99<sup>th</sup> percentile) | 1.59 (99<sup>th</sup> percentile) |

**Table S12.** Real-space correlation coefficients (RSCC) and average B-factors (Å^2^) of inhibitors from PDB validation reports.

| COMPLEX | CHAIN | LIGAND RSCC | LIGAND B-FACTORS ($\text{\AA}^2$) | PROTEIN B-FACTORS ($\text{\AA}^2$) |
| --- | --- | --- | --- | --- |
| KPC-3·#61 | A | 0.99 | 32.8 | 42.7 |
| KPC-3·#61 | B | 0.99 | 37.1 | 50.6 |
| KPC-3·#70 | B | 0.98 | 24.8 | 31.5 |

### LEGENDS FOR SUPPLEMENTARY DATA FILES

**File S1. Validation datasets, benchmark test sets, and raw data used for summarized metrics.** This file contains the following worksheets, in order: the set of 200 PubMed articles and corresponding functional labels used to fine-tune the summarizing large language model (LLM); the test set of 100 examples and the performance of GPT-3.5.turbo-0613, GPT-4-turbo, and GPT-3.5-turbo-0613 fine-tuned on the aforementioned training set; the confusion matrix; a set of 92 manually-curated clusters of labels that should be grouped together, used as the DBSCAN validation set; the performance of DBSCAN across different parameter settings on this validation set; the set of 100 clusters used as a test set to evaluate the final DBSCAN cluster-based label-mapping method, along with its performance; a 186-molecule sample of PubCheF used to validate its label-assignment quality; a performance comparison of a judge LLM and a human evaluator (author K.X.) for assigning confusion-matrix outcomes to predictions across the PubCheF-test-reduced and OpenTargets-indications benchmark sets, using predictions from MolT5, CheF, and PubCheF-1; CheF-v1 predictions and performance on the PubCheF-test-reduced benchmark set; CheF-v1 predictions and performance on the OpenTargets-indications benchmark set; PubCheF-1 predictions and performance on the PubCheF-test-reduced benchmark set; PubCheF-1 predictions and performance on the OpenTargets-indications benchmark set; MolT5 predictions and performance on the PubCheF-test-reduced benchmark set; MolT5 predictions and performance on the OpenTargets-indications benchmark set; PubCheF-1 (ChEMBL) predictions and performance on the PubCheF-test-reduced benchmark set; PubCheF-1 (ChEMBL) predictions and performance on the OpenTargets-indications benchmark set.

**File S2. PubCheF dataset.** The entire PubCheF dataset comprising ∼1.2 million molecules, containing PubChem CIDs, SMILES strings, associated PubMed IDs (PMIDs), and functional labels generated by the dataset creation pipeline. The file is available for download at https://doi.org/10.5281/zenodo.21108754.

**File S3. Predicted bioactive molecules and β-lactamase inhibitors in the ZINC20 in-stock dataset, candidate compound down-selection, and associated ordering and screening parameters.** The “PubCheF-1 5-target analysis” tab contains 100 top and top novel (Tc < 0.50) predictions for each of five targets: SARS-CoV-2 Mpro, 5-HT2A receptor, Taq DNA polymerase, KPC-3 β-lactamase, and Hepatitis C Virus NS5B RNA-dependent RNA polymerase. The “PubCheF-1 predict. bla inhib.” tab contains molecules in the ZINC20 purchasable set with a PubCheF-1 predicted scores greater than 0.05 for any label containing the substring “lactam”. For access to all PubCheF-1 predictions (across the entire ZINC20 database and for all labels), see the PubCheF website, https://www.pubchef.org or https://doi.org/10.5281/zenodo.21108754. The “Filtered compounds” tab contains the molecules that satisfied filtering for structural novelty, low predicted toxicity to HepG2, HSkMC, and IMR-90 cells, absence of Pan-Assay Interference Substructures [16], and availability from preferred vendors, from which 111 molecules were ordered (“Compounds ordered” tab). The “Protein-target screen solvents” tab lists the solvents used to dissolve the ordered compounds (compound #90, highlighted in grey, arrived damaged and was not tested) for the protein target screen. The “In-cell screen solvents” tab lists the solvents used to dissolve the ordered compounds for the in-bacteria screen and all subsequent microbiology experiments.

**File S4. Raw and analyzed data and statistical analyses used to generate Figure 3A**. The “Results summary” tab summarizes the percentage difference in nitrocefin hydrolysis by each tested β-lactamase when comparing a reaction containing a candidate compound with a reaction containing the carrier control. Conditions where nitrocefin hydrolysis significantly decreased are highlighted in pink. Data shown represent three independent experiments using the same purified protein sample. Compound #90 arrived damaged and was not tested, thus it is not included in this analysis. “KPC-3, “L1_#3637_”, “EC-2”, “OXA-10” tabs show the raw OD_490_ measurements at 0 and 10 minutes used to calculate the percentage inhibition for the serine β-lactamases KPC-3, EC-2 and OXA-10 and the metallo-β-lactamase L1#3637. The “Statistical analysis” tabs lists the results of the one-way ANOVA followed by an uncorrected Fisher’s LSD test used to determine significance in this screen. Greyed out cells represent conditions in which increase in nitrocefin hydrolysis was observed and highlighted in pink are conditions where percentage inhibition exceeded 20% and showed statistical significance.

**File S5. Raw and analyzed data and statistical analyses used to generate Figure 3B**. The “Results summary” tab summarizes the fold change of OD_600_ of cells exposed to 32 µg/mL of the candidate compounds in the presence of ∼0.3x MIC of ceftazidime for each tested strain (*E. coli* MC1000 carrying the empty vector pDM1 or expressing either the serine-β-lactamase KPC-3 or the metallo-β-lactamase L1_#3637_) relative to the OD_600_ of cells exposed only to ∼0.3x MIC of ceftazidime (48 µg/mL ceftazidime for KPC-3 or L1_#3637_ and 0.01 µg/mL for pDM1). Conditions where OD_600_ significantly decreased are highlighted in pink. Data shown represent three biological experiments performed as a single technical repeat. Compound #90 arrived damaged and was not tested, thus it is not included in this analysis. The “Carrier controls” tab shows the effects of the carrier controls (DMSO and MeOH) at different concentrations on each of the three tested strains; this data was used to generate Supplementary Figure S4D and represents three biological experiments performed as a single technical repeat. For each tested strain, three worksheets are included. As an example, for *E. coli* MC1000 carrying the empty vector pDM1, the tabs “pDM1 raw data”, “pDM1 analysis” and “pDM1 statistics” are provided. These worksheets show the raw OD_600_ measurements for each 96-well plate used to screen this strain according to the schematic provided in **Supplementary Figure S4C**, the analysis of these data to generate the fold changes presented in the “Results summary” tab, and the statistical analysis of the generated data, respectively. All statistical analyses were performed using a one-way ANOVA followed by an uncorrected Fisher’s LSD test. Greyed out cells represent conditions in which increase in OD_600_ was observed when the compound was present, cell with black text represent conditions in which OD_600_ decreased or remained the same, and pink cells represent conditions in which OD_600_ decreased by 1.4-fold or more and was statistically significant compared to the relevant carrier-control condition.

**File S6. Data used to generate Figure 3B**, **Figure 4A-D**, **Figure 5, Supplementary Figure S4B, Supplementary Figure S5, and Supplementary Figure S12 A-C. (A)** Ceftazidime MIC values for *E. coli* MC1000 (K-12 strain) carrying the empty vector pDM1 or expressing either KPC-3 or L1_#3637_ used to generate **Supplementary Figure S4B** and to determine the ceftazidime concentration (∼0.3 x MIC) for the in-bacteria screen (**Figure 3B**). **(B)** MIC values for *E. coli* MC1000 strains harboring pDM1 or expressing KPC (KPC-3 and KPC-2) or L1 variants (L1_#3637_, L1_#4488_, and L1_#4509_) in the presence of all compound hits identified in **Figure 3B**, used to generate **Figure 4A-D** and **Supplementary Figure S5**. **(C)** Ceftazidime, imipenem, and aztreonam MIC values for clinical isolates of *E. cloacae*, *K. pneumoniae*, *C. freundii* and *E. coli* in the presence of inhibitor hits #61 and #70 or the DMSO carrier control (added at an equivalent volume) used to generate **Figure 5A-C**. **(D)** Ceftazidime MIC values for a clinical *S. maltophilia* isolate (strain AMM) that harbors a deletion of its *bla_L2_* gene in the presence of inhibitor hits #35 and #42 or the DMSO carrier control (added at an equivalent volume) used to generate **Figure 5D**. **(E)** Imipenem, aztreonam, and ceftazidime MIC values for clinical isolates expressing KPC-2 (*E. cloacae* HPTU 27040557, *E. coli* LIL, and *K. pneumoniae* ZRI) in the presence of decreasing concentrations of inhibitor hits #61 and #70 or the DMSO carrier control (added at an equivalent volume) used to generate **Supplementary Figure S12A-C**. For all MIC values, experiments were performed as three biological experiments, each conducted as a single technical repeat.

**File S7. Data used to generate Figure 6 and Supplementary Figure S12D. (A)** Number and timing of *Galleria mellonella* larval deaths used to generate the Kaplan-Meier plots in **Figure 6A**,**B**. **(B)** CFU values per thigh for the *K. pneumoniae* ZRI clinical strain isolated from neutropenic mice at 2 hours post-infection (no treatment); these data were used to generate **Figure 6C,D**. **(C)** CFU values per thigh for the *K. pneumoniae* ZRI clinical strain isolated from neutropenic mice after treatment with **(1)** the carrier control (PBS + DMSO), **(2)** antibiotic only (ceftazidime + DMSO), **(3)** inhibitor compound only (PBS + #61), or **(4)** combination treatment (ceftazidime + inhibitor hit #61); these data were used to generate **Figure 6C**. **(D)** CFU values per thigh for the *K. pneumoniae* ZRI clinical strain isolated from neutropenic mice after treatment with **(1)** the carrier control (PBS + DMSO), **(2)** antibiotic only (ceftazidime + DMSO), **(3)** inhibitor compound only (PBS + #70), or **(4)** combination treatment (ceftazidime + inhibitor hit #70); these data were used to generate **Figure 6D**.

## Notes

https://www.pubchef.org

https://doi.org/10.5281/zenodo.21108754

